# Targeted Protein Acetylation in Cells Using Heterobifunctional Molecules

**DOI:** 10.1101/2021.07.27.454011

**Authors:** Wesley W. Wang, Li-Yun Chen, Jacob M. Wozniak, Appaso M. Jadhav, Hayden Anderson, Taylor E. Malone, Christopher G. Parker

## Abstract

Protein acetylation is a central event in orchestrating diverse cellular processes. However, current strategies to investigate protein acetylation in cells are often non-specific or lack temporal and magnitude control. Here, we developed an acetylation tagging system, AceTAG, to induce acetylation of targeted proteins. The AceTAG system utilizes bifunctional molecules to direct the lysine acetyltransferase p300/CBP to proteins fused with the small protein tag FKBP12^F36V^, resulting in their induced acetylation. Using AceTAG, we induced targeted acetylation of a diverse array of proteins in cells, specifically histone H3.3, the NF-κB subunit p65/RelA, and the tumor suppressor p53. We demonstrate that targeted acetylation with the AceTAG system is rapid, selective, reversible, and can be controlled in a dose-dependent fashion. AceTAG represents a useful strategy to modulate protein acetylation and will enable the exploration of targeted acetylation in basic biological and therapeutic contexts.

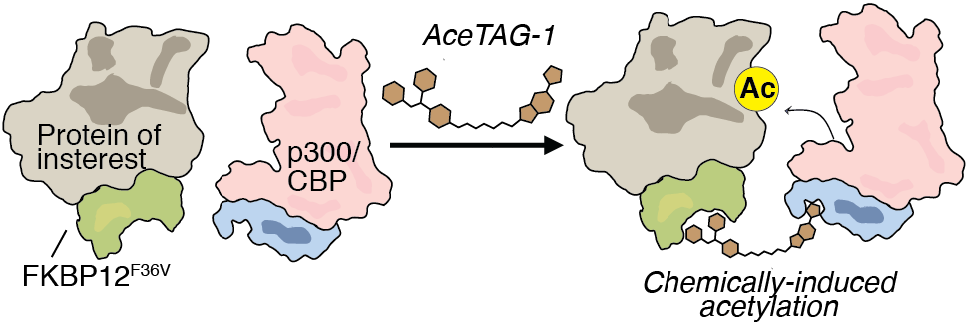

## INTRODUCTION

Lysine acetylation is one of the most frequent post-translational modification (PTM), occurring on >10,000 sites on human proteins, and plays critical roles in human biology^1^. This covalent modification is reversible, and its dynamic equilibrium is mediated by a combined ∼40 lysine deacetylases (KDACs) and acetyltransferases (KATs)^2–3^. Acetylation not only imparts direct functional consequences on proteins, but can also interplay with other PTMs, such as phosphorylation, ubiquitination, methylation and SUMOylation, and its dysregulation has been implicated in diverse human diseases, including various cancers,^4–5^ neurodegenerative disorders ^6–7^, autoimmune conditions^8–9^ and metabolic disease^5, 10–11^. Despite its frequency and diverse roles, a substantial hurdle for accurate functional investigations of acetylation is the dearth of tools to selectively modulate protein acetylation in cells. Although conventional methods, such as genetic or chemical manipulation of KDACs or KATs, have yielded much insight into the myriad roles of acetylation,^12–13^ they induce global alterations to their substrates, potentially complicating the interpretation of the biological effects relating to a single protein.^14–15^ Alternatively, methods for site-specific modification of substrates, either to mimic or block acetylation,^16–17^ could have undesirable effects on protein structure, obfuscates analysis of alternate PTMs on the same residues, and do not permit acute or graded investigations of acetylation in cells. To overcome these obstacles, we envisioned a potentially generalizable, chemical-based strategy to dynamically and selectively induce acetylation on a target protein of interest (POI) directly in cells.

Encouraged by the transformative utility of chemical inducers of dimerization,^18–19^ ubiquitin-inducing small molecules,^20^ and the recent examples of phosphorylation modifying bifunctional molecules^21–22^, we hypothesized that a similar heterobifunctional system could be employed to modulate the acetylation of any POI on-demand by proximity-induced acetylation. Acetylation can impart inhibitory effects on protein as well as activating or gain-of-function outcomes, thus a potential hurdle for the development of such heterobifunctional small molecules is the identification of functionally ‘silent’ ligands that selectively bind to target proteins without overt effects on protein function to prevent counterproductive or complicated outputs. Considering this, and inspired by various technologies that enable the selective targeting of genetically-tagged proteins with small molecules,^23–25^ we conceptualized a strategy to direct protein acetylation to targeted POIs, even in the absence of available small molecule ligands.

Here, we report the development of an ‘acetylation tagging’ system, or AceTAG, which consists of a heterobifunctional molecule formed by covalently linking a KAT-binding ligand to an FKBP12^F36V^-binding ligand, resulting in chemically induced proximity between cellular acetylation machinery and a targeted protein. Using AceTAG, we demonstrate that the KAT p300/CBP can be recruited directly to histone H3.3, p65/RELA and p53, resulting in their targeted acetylation at functionally relevant sites in cells. We demonstrate AceTAG-mediated acetylation is dose-controlled, reversible, rapid, occurring within minutes, and selective for tagged protein targets.

## RESULTS AND DISCUSSION

### Bifunctional AceTAG molecules mediate binding between p300/CBP and FKBP12^F36V^

Among KATs, the E1A-binding protein (p300) and its paralog CREB-binding protein (CBP) are two of the most prominent members, playing key roles as transcriptional co-activators essential for myriad cellular processes, including growth and development, stress response, oncogenesis, and DNA damage^26^. p300/CBP collectively regulate >2/3 of the known acetylation sites in humans^27^, indicative of a broad substrate tolerability and therefore represent high-priority candidates for the design of generalizable chemically-induced acetylation strategies. Recently, inhibitors targeting the conserved p300/CBP bromodomain (BRD)^28^ have been co-opted into bifunctional molecules to target chromatin machinery. This includes their conjugation to DNA minor groove targeting polyamides^29^ to coordinate histone acetylation *in vitro* and alter gene expression *in cellulo* as well as their use to recruit p300/CBP to gene loci for targeted gene transcription via catalytically inactive Cas9^30^, the latter presumably through proximity-induced histone acetylation. However, it remained an open question as to whether a potentially generalizable strategy might be developed to recruit p300/CBP directly to protein targets, resulting in their targeted acetylation in living cells.

Recently, in an approach called degradation TAG (dTAG), it has been shown that synthetic ligands that recognize the engineered *FKBP12* variant FKBP12^F36V^ can be conjugated to E3 ligands,^31^ resulting in selective ubiquitination and proteasome-mediated degradation of FKBP12^F36V^-tagged proteins^25, 32^. We hypothesized a similar strategy could be adopted for targeted protein acetylation (Figure 1a). To this end, we appended the previously reported FKBP12^F36V^-binding ligand^25^ to the 5-isoxazoly-benzimidizole p300/CBP BRD-inhibitor^28^, to generate a series compounds, AceTAG 1-3, with different linker compositions (Figure 1b, Supplementary Figure 1a). We first assessed the ability of AceTAG1-3 to mediate complex formation between soluble recombinant FKBP12^F36V^ and the BRD domain of p300 (BRD-p300) *in vitro* using an AlphaScreen assay^33^ (Supplementary Figure 1b). We observed characteristic bell-shaped auto-inhibitory curves for all AceTAG analogs, the result of AceTAG molecules saturating both FKBP12^F36V^ and BRD-p300, effectively outcompeting ternary complex formation^34^. We noted that AceTAG-1 has increased potency relative to other analogs, with maximum complex formation occurring at ∼1 μM (Figure 1c). Further, the luminescence signal is considerably diminished upon co-treatment of either terminal binding molecules (FKBP-c and p300-c) in a concentration-dependent fashion (Figure 1d-e, Supplementary Figure 1c), confirming that observed ternary complex formation is dependent on the simultaneous binding of both BRD-p300 and FKBP12^F36V^. To verify that AceTAG molecules can engage their protein binding partners in cells, we constructed ‘fully-functionalized’ photoaffinity probes of both the FKBP12^F36V^ (FKBP-p) and p300/CBP (p300-p) ligands and confirmed respective binding to recombinantly expressed p300 and FKBP12^F36V^ can be efficiently competed when cells are co-treated with increasing concentrations AceTAG-1 (Figure 1f-g). Together, these biochemical data indicate that AceTAG 1-3 effectively engage FKBP12^F36V^ and p300/CBP and are capable of inducing ternary complex formation.

**Figure 1.**
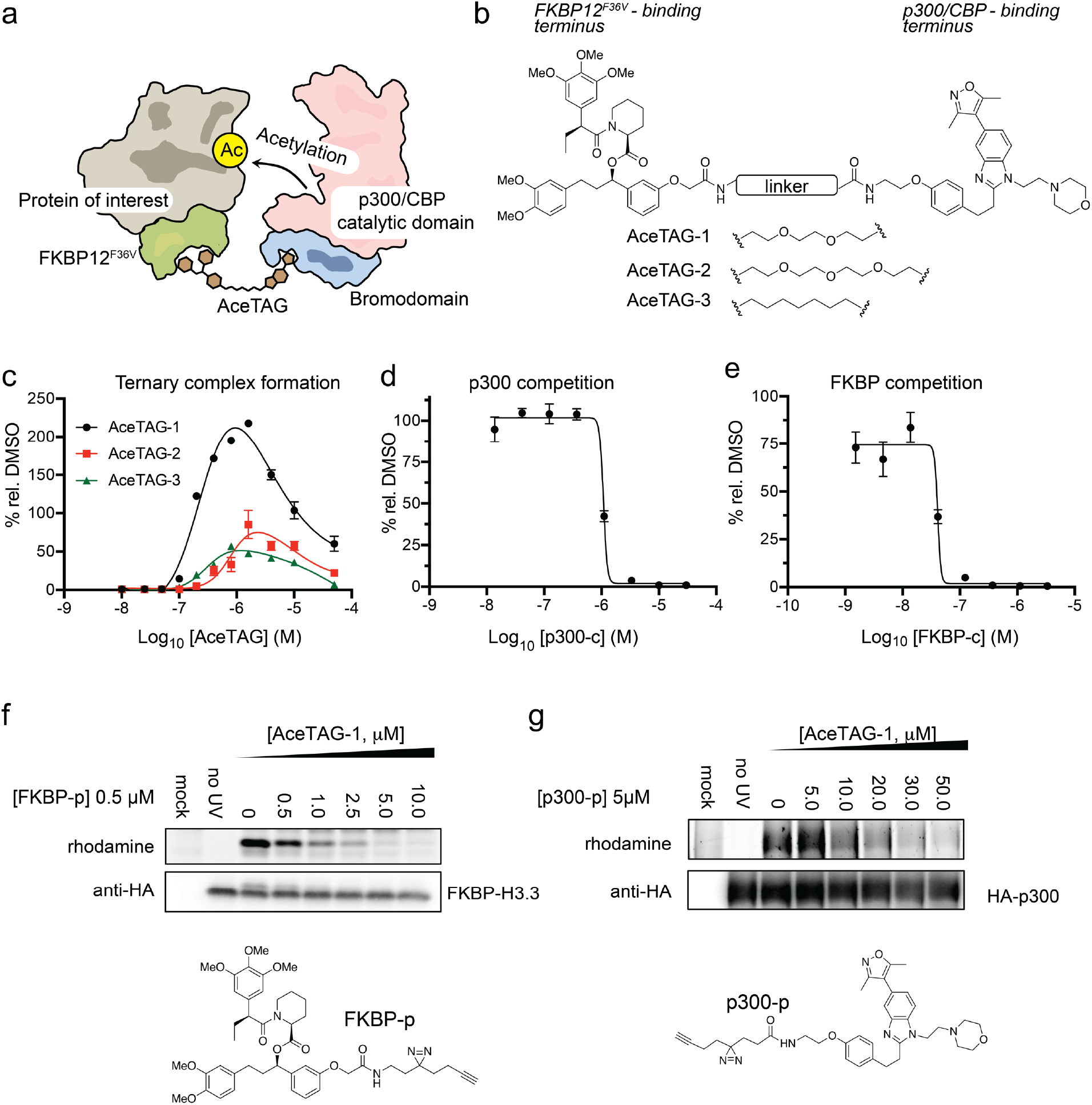
AceTAG molecules bind p300/CBP and FKBP12^F36V^ and mediate ternary complex formation. **a,** Schematic overview of targeted acetylation tagging (AceTAG) strategy. Hetereobifunctional AceTAG molecules induce acetylation of FKBP12^F36V^-POI fusions through recruitment of KAT p300/CBP. **b**, Structures of AceTAG molecules composed of an FKBP12^F36V^ binding ligand and a p300/CBP-binding ligand connected by a linker. **c**, AceTAG molecules induce ternary complex between FKBP12^F36V^ and the bromodomain of p300 (BRD-p300) as determined by AlphaScreen (Supplementary Figure 1b). **d-e**, AceTAG-1 (300 nM) mediated ternary complex formation between FKBP12^F36V^ and BRD-p300 can be blocked upon coincubation of increasing concentrations of p300-binding (**d**) or FKBP12^F36V^-binding ligands (**e,** Supplementary Figure 1c). Data in **c-e** represent as mean ± s.d. of *n* = 3 replicates. **f-g**, Confirmation of AceTAG-1 target engagement in cells. FKBP12^F36V^-H3.3 (**f**) and (**g**) were recombinantly expressed with HA epitope tags by transient transfection in HEK293T cells, which were then co-treated with a photoaffinity probe derived from the FKBP12^F36V^ ligand (**f**, FKBP-p) or p300/CBP ligand (**g**, p300-p) and increasing concentrations of AceTAG-1, photocrosslinked and lysed, and proteomes were conjugated to an azide-rhodamine tag by CuAAC chemistry and analyzed by in-gel fluorescence staining. The results in **f**-**g** are representative of three independent biological replicates (*n* = 3). Full images of gels are shown in Supplementary Figure 6.

### AceTAG Molecules Induce Targeted Protein Acetylation in Cells

We next evaluated whether AceTAG molecules could induce acetylation of FKBP12^F36V^-tagged proteins in cells. We chose first to target histone H3.3 given that p300/CBP is a predominant histone acetyl transferase. Although the specific sites on H3 targeted by p300/CBP are not firmly established, recent proteomic studies have indicated that pharmacological inhibition and genetic ablation of p300/CBP results in decreased levels of H3K18ac, H3K27ac and H3K36ac^15^. In addition, *in vitro* studies suggest H3K14, H3K18, H3K23, H3K64 and H3K122 are also major acetylation targets^35–37^. We first stably transfected HeLa cells with H3.3-FKBP12^F36V^-HA and confirmed that the H3.3-FKBP12^F36V^ chimera localizes primarily to chromatin along with endogenous H3.3 (Supplementary Figure 2a). We next evaluated AceTAG compounds AceTAG 1-3 for their relative ability to induce acetylation at H3.3 K18 using acetyl-histone H3 K18 selective antibodies. We observed AceTAG-mediated K18 acetylation for all analogs, with AceTAG-1 inducing modestly higher levels of acetylation relative to the other analogs (Figure 2a, Supplementary Figure 2b).

**Figure 2.**
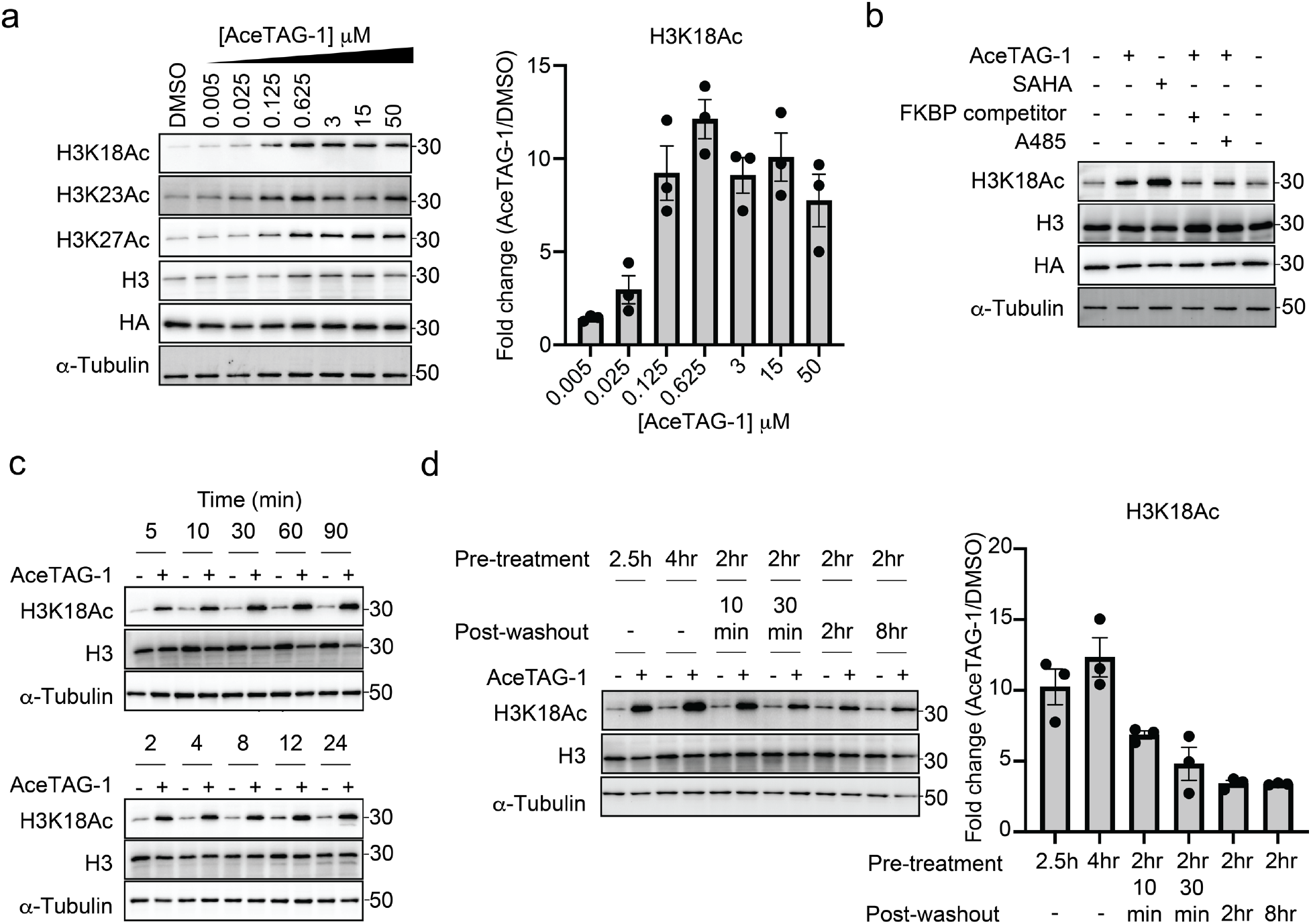
AceTAG molecules induce rapid, targeted acetylation of H3.3-FKBP12^F36V^ in cells. **a**, Concentration dependent acetylation of H3.3-FKBP12^F36V^ lysine residues by AceTAG-1. H3.3-FKBP12^F36V^ HeLa cells were treated with increasing concentrations of AceTAG-1 for 2 h and acetylation of K18, K27, and K23 were monitored by immunoblot. Shown in the right panel is quantitation of immunoblot signal of K18Ac relative to α-tubulin as the mean ± s.e.m. of *n* = 3 biologically independent experiments. **b**, Immunoblot analysis of controls for AceTAG-mediated acetylation of H3.3. Cells were treated with the histone deacetylase inhibitor SAHA (5 μM) or co-treated with AceTAG-1 (625 nM) and DMSO, an FKBP12^F36V^ binding ligand (50 μM, Supplemental Figure 1c), or the p300/CBP KAT domain inhibitor A-485 (1 μM) for 2 h which block AceTAG-1-induced acetylation. The results in **a**-**b** are representative of two independent biological replicates (*n* = 2). **c**, Immunoblot analysis of AceTAG-1 (625 nM) treated H3.3-FKBP12^F36V^ HeLa cells over the indicated time course. **d**, Washout experiments showing decreasing acetylation upon removal of AceTAG-1 from cells. Immunoblot of H3.3-FKBP12^F36V^ HeLa cells pretreated with AceTAG-1 (625 nM) or vehicle for the indicated time, washed with DPBS and resuspended in fresh media (without AceTAG-1) for the indicated time. Shown in the right panel is quantitation of immunoblot signal of K18Ac relative to α-tubulin as the mean ± s.e.m. of *n* = 3 biologically independent experiments. The results in **c-d** are representative of three independent biological replicates (*n* = 3). Full images of blots are shown in Supplementary Figures 7-8.

We observe that AceTAG-mediated acetylation of H3.3-FKBP12^F36V^ is dose-dependent, with highest levels of acetylation achieved between 625 nM and 3 μM, near the concentration needed for maximum ternary complex formation *in vitro* (Figure 1c), and observed decreased acetylation levels at the highest AceTAG-1 concentrations, consistent with acetylation being a consequence of ternary complex induction in cells (Figure 2a). We also confirmed increased H3.3-FKBP12^F36V^ acetylation upon incubation of cells with the broad histone deacetylase (HDAC) inhibitor suberoylanilide hydroxamic acid (SAHA), indicating the fusion protein is a substrate of endogenous HDACs (Figure 2b). We next examined levels at other known H3.3 sites using available antibodies, including K9, K14, K23, K27, and K79 (Figure 2a, Supplementary Figure 2c), detecting induced acetylation only at K18, K23 and K27 (Figure 2a). Notably, we did not observe induced acetylation of un-tagged, endogenous H3.3 after AceTAG-1 treatment (Supplementary Figure 2d). Further, acetylation can be blocked both by coincubation with either A-485, a p300/CBP KAT inhibitor^38^, or with excess FKBP12^F36V^ ligand (FKBP-c, Figure 2b, Supplementary Figure 1c). Collectively, these data indicate that AceTAG-induced acetylation in cells is dependent on p300/CBP enzymatic activity and is targeted to the FKBP12^F36V^ tagged protein.

We next set out to assess the kinetics of AceTAG-mediated acetylation in cells by monitoring acetylation levels of H3.3 K18 over multiple time points. Surprisingly, we observed that induced acetylation occurs almost immediately, after only ∼5min exposure to AceTAG-1 (Figure 2c) as well as after continuous incubation (up to 24 hrs). Further, when AceTAG-1 is removed from cells, H3.3 K18 acetylation steadily dissipates, reaching near basal levels within 2hr after compound removal (Figure 2d). This re-equilibration is likely the result of endogenous KDAC activity, together indicating that the targeted acetylation is reversible and dependent on the heterobifunctional compound.

### AceTAG Strategy Extendable to Multiple Proteins

Satisfied with the successful compound-induced acetylation of H3.3, we next used our AceTAG system to assess chemically mediated acetylation of other targeted proteins. To avoid potential obfuscation by endogenous target proteins, we generated FKBP12^F36V^-RelA and FKBP12^F36V^-p53 constructs, two proteins wherein p300/CBP-mediated acetylation is known to impart consequences^39–40^ on their transcriptional activity. Subsequently, FKBP12^F36V^-RelA was stably transfected in a HeLa *RelA*^-/-^ cell line while FKBP12^F36V^-p53 was stably transfected in H1299 non-small cell carcinoma (NSCLC) cells, which have a homozygous partial deletion of *TP53* and lack p53 protein expression. Treatment of FKBP12^F36V^-RelA expressing cells with AceTAG-1 resulted dose-dependent acetylation at K310, a site previously proposed to be a predominant target of p300/CBP (Figure 3a-b),^41–42^ with minimal effects on neighboring K314/315 (Supplementary Figure 3a). In FKBP12^F36V^-p53 H1299 cells, we monitored acetylation of the N-terminal domain of p53, specifically at K305, K373, and K382, sites previously suggested to be substrates of p300/CBP,^43–46^ where strong AceTAG-1-dependent acetylation was also observed (Figure 3c-d).

**Figure 3.**
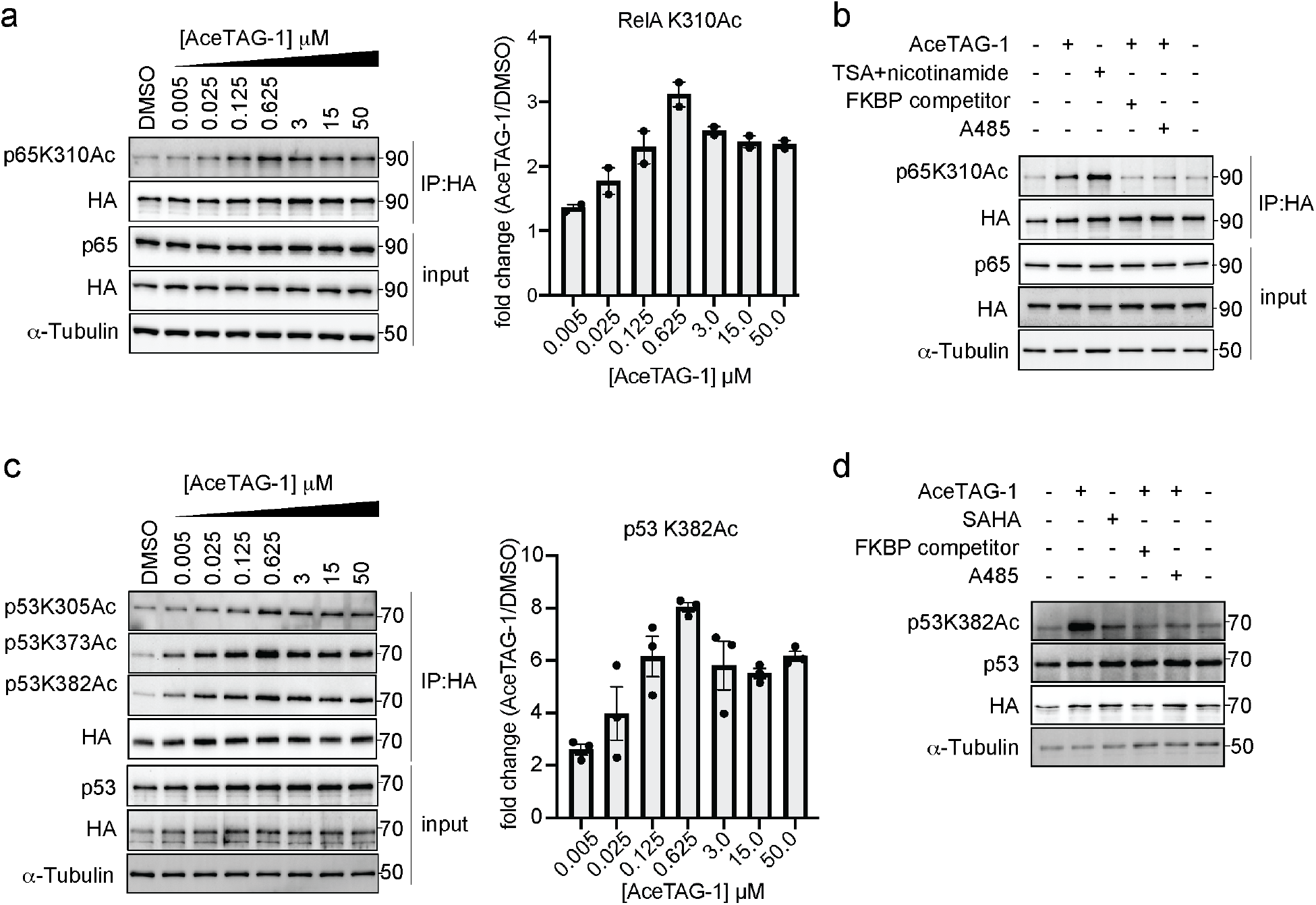
AceTAG targeted acetylation can be applied to multiple POIs. **a**, AceTAG-1 induces acetylation FKBP12^F36V^-p65/RelA fusion in HeLa cells. FKBP12^F36V^-RelA-HA RelA^-/-^ HeLa cells were treated with increasing concentrations of AceTAG-1 (2 h), lysed and FKBP12^F36V^-RelA was enriched and blotted with noted antibodies. Shown in the right panel is quantitation of immunoblot signal for p65 K310Ac relative to FKBP12^F36V^-RelA-HA as the mean ± s.e.m. of *n* = 2 biologically independent experiments. **b**, Immunoblot analysis of controls for AceTAG-mediated acetylation of RelA. Cells were treated with TSA (2 μM) and nicotinamide (10 mM) as a positive control for p65 acetylation^47^ or co-treated with AceTAG-1 (625 nM) and DMSO, an FKBP12^F36V^ binding ligand (Supplementary Figure 1c), or the p300/CBP KAT domain inhibitor A485 for 2 h which block AceTAG-1-induced acetylation. **c**, AceTAG-1 induces acetylation FKBP12^F36V^-p53 fusion in H1299 cells. FKBP12^F36V^-p53-HA H1299 were treated with increasing concentrations of AceTAG-1 (2 h), lysed and FKBP12^F36V^-p53 was enriched and blotted with noted antibodies. Shown in the right panel is quantitation of immunoblot signal for p53 K382Ac relative to FKBP12^F36V^-p53-HA as the mean ± s.e.m. of *n* = 3 biologically independent experiments. **d,** Immunoblot analysis of controls for AceTAG-mediated acetylation of p53. The results in **b,d** are representative of two independent biological replicates (*n* = 2). Full images of blots are shown in Supplementary Figures 9-10.

Consistent with our previous observations, we observe apparent acetylation autoinhibition for both targets at higher AceTAG-1 concentrations (Figure 3a, c) and blockade of induced acetylation upon co-treatment of cells with p300/CBP KAT inhibitor A-485 or competing FKBP ligand (Figure 3b, d). Further, a significant level of p53 acetylation is detectable after ∼10 minutes of AceTAG compound treatment (Supplementary Figure 3b), similar to the kinetics observed for H3.3. Finally, we note that AceTAG-1 induced acetylation does not appear dependent upon the positioning of the FKBP12^F36V^ tag, as similar acetylation effects were observed with C-terminally tagged p53 (Supplementary Figure 3c).

### Selectivity of AceTAG-Induced Acetylation

To more robustly characterize the site selectivity of induced acetylation using AceTAG, we monitored AceTAG-1 induced acetylation of a targeted protein in cells by quantitative mass spectrometry (MS). In these experiments, H3.3-FKBP12^F36V^-HA was enriched from stably transfected HeLa cells treated with (1) DMSO, (2) AceTAG-1, or (3) SAHA and trypsinized for protein identification and quantitation using tandem mass tags (TMT) (Supplementary Figure 4a). In line with immunoblotting experiments (Figure 2a, Supplementary Figure 2c), we measured substantial increases in acetylation at K18, K23, and K27, but detected no acetylation at other identified lysines on H3.3 (Figure 4a, Supplementary Figure 4b). Together, these data suggest that when p300/CBP is recruited to H3.3 in cells using our AceTAG system, the preferred acetylation sites are K18, K23, and K27.

**Figure 4.**
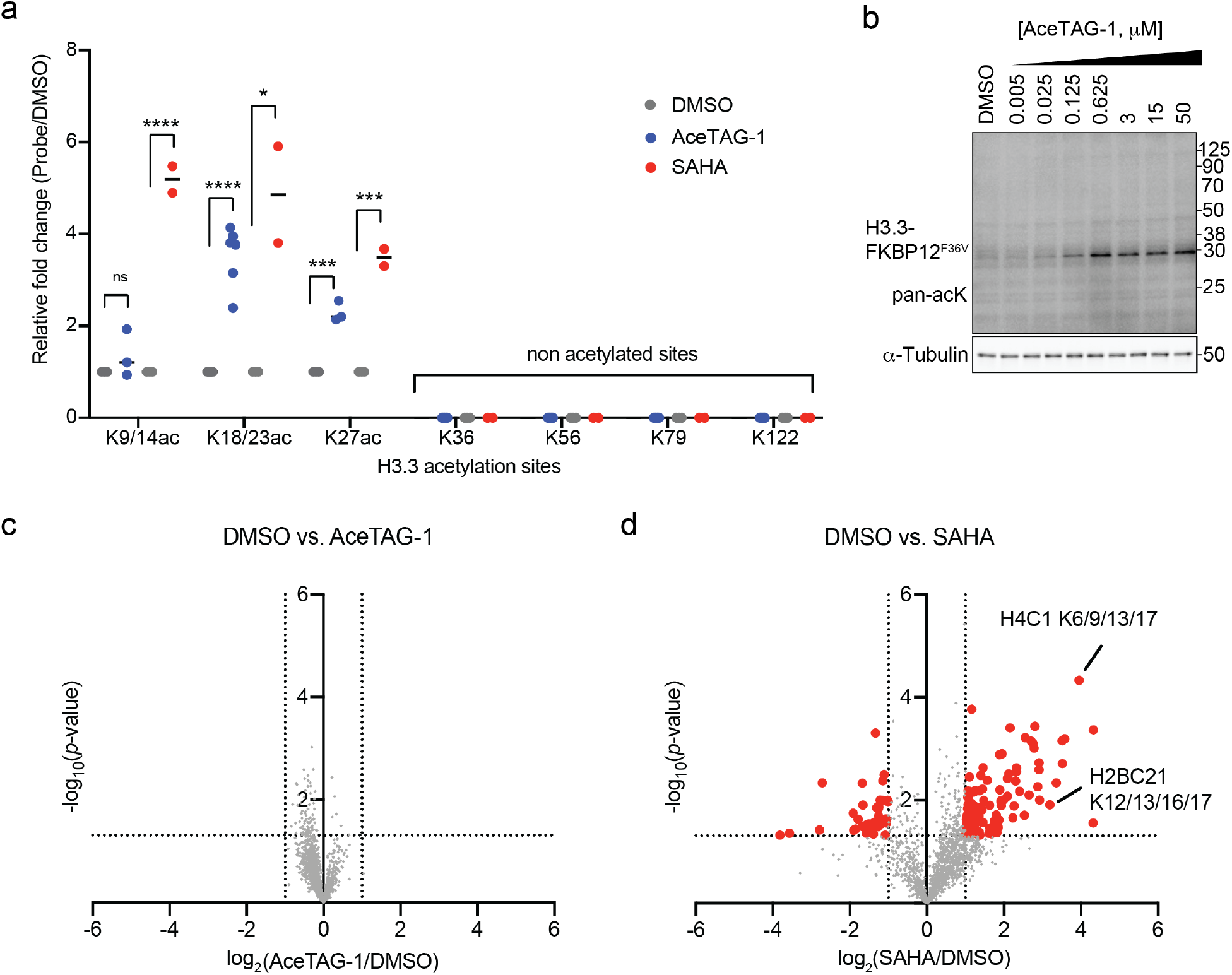
Chemically induced acetylation selectivity assessment using quantitative proteomics. **a**, Fold changes in H3.3 acetylated lysine abundances of H3.3-FKBP12^F36V^ HeLa cells treated with AceTAG-1. Cells were treated with DMSO, AceTAG-1 (625 nM) or SAHA (5 μM) for 2 or 24 h respectively, lysed, and H3.3 was enriched, trypsinized and labeled with isobaric TMT reagents, then subjected to quantitative MS (outlined in Supplementary Figure 3a, Supplementary Table 1). Data represents median of *n* = 6 biological replicates for AceTAG-1 treated cells and median of *n* = 2 biological replicates for SAHA-treated cells. Note, peptides may not have been detected in every replicate, but are required to be detected and quantified in at least two biological replicates to be included in this analysis. No acetylation detected for K36, K56, K79, and K122. Statistical significance was calculated with unpaired two-tailed Student’s *t*-tests comparing DMSO- to AceTAG-1 or SAHA-treated samples. \**P* < 0.05; \*\*\**P* < 0.001; \*\*\*\**P* < 0.0001. See Supplementary Figure 4b for representative spectra. **b**, AceTAG-1 does not induce broad changes in acetylation as indicated by immunoblot analysis of H3.3-FKBP12^F36V^ HeLa cells treated with increasing concentrations AceTAG-1 using a pan acetyl-lysine antibody. The results are representative of three independent biological replicates (*n* = 3). Full images of blots are shown in Supplementary Figure 11. **c,d**, Volcano plots showing that AceTAG-1 does not induce broad changes in acetylation in the proteome as determined by quantitative MS. H3.3-FKBP12^F36V^ HeLa cells treated with DMSO, AceTAG-1 (600 nM, **c**) or SAHA (2 μM, **d**) for 2 or 16 h respectively, lysed, trypsinized and acetylated peptides enriched and labeled with TMT reagents, then subjected to quantitative MS (Supplementary Table 1). The vertical dashed lines correspond to 2-fold change in enrichment relative to DMSO and the horizontal line corresponds to a *P*-value of 0.05 for statistical significance. Red circles correspond to protein targets with >2-fold change (*P* < 0.05) relative to DMSO. Each point represents an individual acetylated peptide plotted as a mean of *n* = 3 biological replicates for DMSO and AceTAG-1 treatments and *n* = 2 biological replicates for SAHA treatments combined in one TMT 10-plex experiment. Independent replicated TMT experiment shown in Supplementary Figure 5. Examples of histone-derived sites with increased acetylation upon SAHA treatment are noted in **d**.

We next evaluated the selectivity of AceTAG-induced acetylation across the human proteome. Treatment of HeLa cells with AceTAG-1 resulted in no observable acetylation changes in well-established p300/CBP substrates, including c-Myc and STAT3 (Supplementary Figure 5a) or broader acetylation perturbation via immunoblot assays (Figure 4b). To assess targeted acetylation selectivity more globally and quantitatively, we performed MS-based acetyl-proteomic analysis of AceTAG-treated H3.3-FKBP12^F36V^ HeLa cells. Briefly, acetylated peptides from H3.3-FKBP12^F36V^ HeLa cells treated either with (1) DMSO, (2) AceTAG-1, or (3) SAHA were digested, enriched, identified and quantified using TMT (Supplementary Figure 5b). Relative to DMSO, we observed no substantial changes in acetylation of any detected protein in AceTAG-treated cells while SAHA treatment resulted in increases across ∼30 proteins (Figure 4c-d, Supplementary Figure 5c-e, Supplementary Table 1). We note that no substantial changes in endogenous histone lysine acetylation was observed upon AceTAG treatment, including H3.3, likely due to the substantially lower H3.3-FKBP12^F36V^ acetylation and expression levels (∼25 fold) relative to endogenous histones (Supplementary Figure 5f). Collectively, these data suggest that AceTAG-1 mediated acetylation is endowed with exquisite selectivity for FKBP12^F36V^-tagged proteins.

## CONCLUSION

Here, we describe a method for the selective acetylation of targeted proteins in live cells using heterobifunctional small molecules. We demonstrate that AceTAG molecules can bind to and recruit the lysine acetyltransferase p300/CBP to multiple protein targets genetically fused with FKBP12^F36V^, effectively inducing acetylation. We show AceTAG-mediated acetylation is rapid, occurring within minutes of molecule addition, selective, reversible, and dependent upon KAT enzymatic activity. Further, AceTAG molecules possess submicromolar efficacy in live cells, and as expected, possess characteristics associated with heterobifunctional small molecules, including autoinhibition at increased concentrations both *in vitro* and *in celluo* as well as reduced ternary complex formation with competing terminal ligands.

Despite the consequential roles of protein acetylation, methods to study the effects of acetylation on specific protein targets in cells are limited. Common approaches include genetic or pharmacological ablation of acetylation machinery which affects global substrates, potentially complicating any downstream analyses, or substrate mutagenesis, which lacks dynamic control and can perturb substrate structure or potentially interferes with competing PTMs. Thus, the ability to selectively induce acetylation of targeted proteins overcomes many of these challenges. Towards this end, ‘all-chemical’ based approaches have yielded ligand-directed acetyl-donating reagents for stoichiometric, non-enzymatic transfer of acyl groups to lysine residues on specific proteins like androgen receptor^48^, histone H2B^49–50^, dihydrofolate reductase^51–52^, and phosphoglycerate mutase 1^53^. Though capable of targeting endogenous POIs, such reagents are ‘single use,’ allowing for the transfer of one acetyl group per molecule and, further, are limited to proteins with available ligands that bind within proximity to targeted recipient lysine residues. In addition, recent approaches have established that recruitment of p300/CBP to specific DNA sequences with bifunctional molecules can induce transcriptional modulation^29–30, 54^. However, these approaches rely on DNA-targeting polyamides or sequence-targeting using guide RNA’s via engineered Cas9. Our AceTAG system enables the direct targeting of tagged proteins for acetylation, even in the absence of available targeting ligands, in principle enabling the study p300/CBP-mediated acetylation on a wide range of targets.

There are some important points when considering implementation of AceTAG to study protein acetylation. First, we recognize that our current studies have focused on three protein targets. Although current evidence deems p300/CBP to have the largest substrate of all KATs,^1^ it is unclear how generalizable this approach might be to induce acetylation of other substrates or even *neo*-substrates. Thus, future attention should be given to exploring a broader subset of targets from diverse functional classes. Further, even though we conducted a brief exploration of linker length and composition, we do observe linker-dependent ternary complex formation and targeted acetylation (Figure 1c-e, Supplementary Figure 2b). Like other bifunctional molecules,^55–57^ we suspect that each protein target will likely have unique linker preferences, necessitating thorough linker optimization studies. In addition, we recognize that the functional effects of acetylation on protein targets are often site-specific and it is not yet clear if site-selective acetylation can be achieved using heterobifunctional molecule systems. In our studies, we note consistent acetylation of observable sites on p53 using both C- and N-terminal FKBP12^F36V^ fusion constructs, suggesting that selectivity is largely driven by the substrate recognition of the recruited acetylation machinery. Furthermore, AceTAG should be fully compatible with CRISPR-knock in technologies, which would benefit functional studies of target proteins. Lastly, the utility of our AceTAG system is limited by the availability and reliability of methods to monitor acetylation. Currently, the majority of proteins with evidence of acetylation lack site-specific antibodies, and acetylation detection by MS-based proteomics typically requires pan anti-acetyl lysine antibodies for enrichment and detection, which themselves have limited coverage.^58^ However, as we demonstrate in the case of H3.3, the use of the FKBP12^F36V^-HA fusion constructs enables the direct enrichment and monitoring of target proteins by quantitative MS, partially alleviating the dependence on antibodies to explore new target areas.

Finally, we believe that AceTAG-based strategies can be applied to investigate functional consequences of acetylation in cells. Towards this end, have shown that targeted acetylation occurs at functional residues on proteins such as H3.3, p65/RelA, and p53, providing evidence that AceTAG represents an attractive tool to study the various roles of acetylation in cells. Looking forward, we envision similar strategies can be extended to target endogenous proteins for acetylation, bypassing the requirement of genetic manipulation altogether. Given the manifold roles acetylation can have on protein function, we suspect that the identification of functionally “silent” ligands or binders^59^ will prove necessary for the generation of such acetylation-targeting chimeric small molecules. In this regard, powerful ligand discovery technologies, including chemoproteomic-based methods^60–61^ and DNA-encoded libraries (DELs)^62^, should prove fruitful. In conclusion, we envision AceTAG will not only serve as useful tool to study basic biology but could also enable the exploration of chemically-induced acetylation as a therapeutic strategy.

## ASSOCIATED CONTENT

### Supporting Information

Supplementary figures, tables and experimental details can be found in attached supporting documents. The mass spectrometry proteomics data has been deposited to the ProteomeXchange Consortium via the PRIDE partner repository, which will be made public upon acceptance of manuscript.

## ACKNOWLEDGMENTS

The authors acknowledge the Scripps NMR, MS, Genetic Perturbation Screening and Genomics core facilities. We thank Dr. Michael Erb (Scripps Research) and Timothy Bishop (Scripps Research) for their useful discussions.

## Supplementary Figures

**Supplementary Figure 1.**
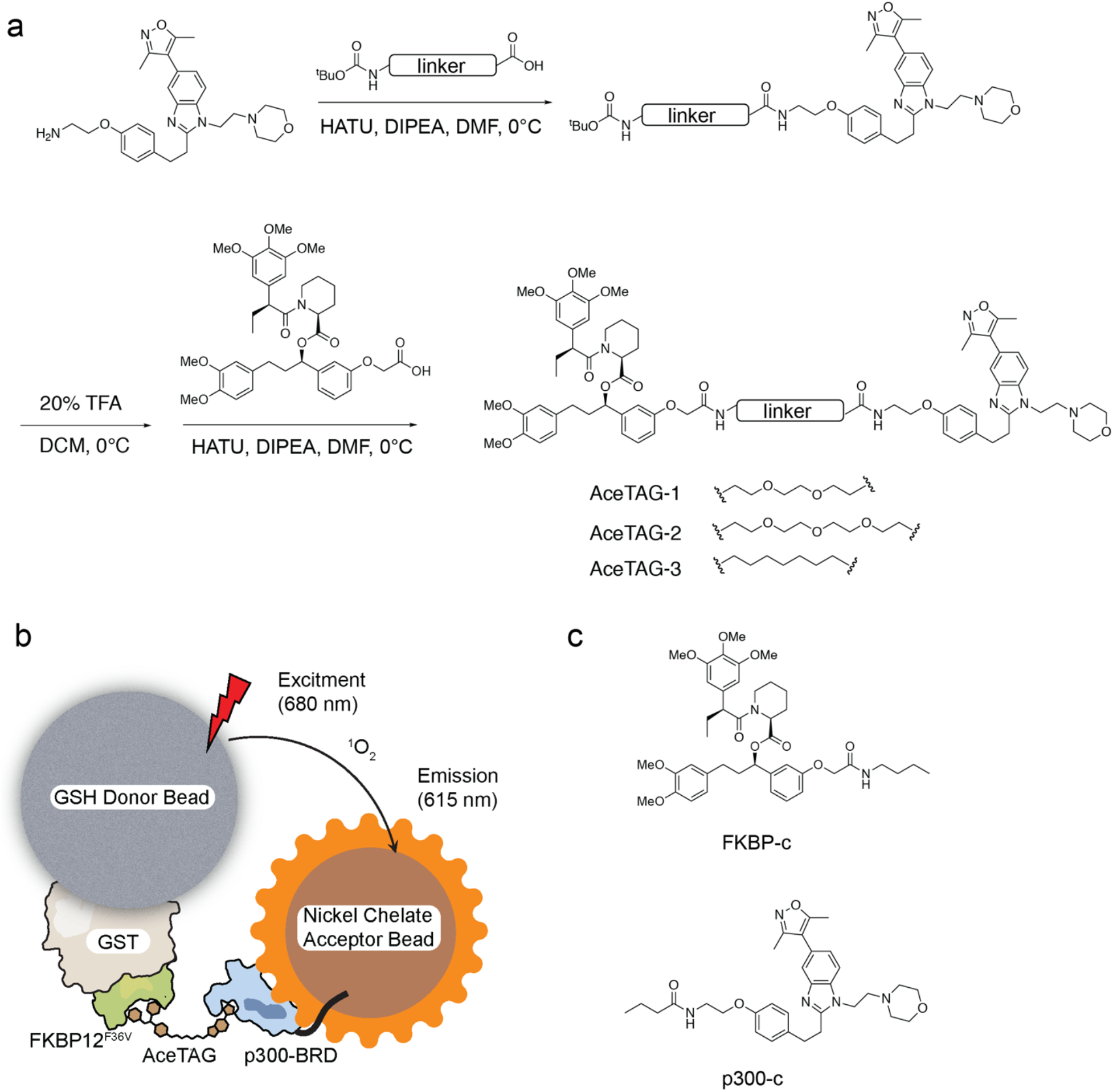
Chemical structures of indicated molecules and AlphaScreen assay design. **a**, General synthetic route of AceTAG molecules. **b**, Schematic depiction of a luminescence-based AlphaScreen assay, in which AceTAG molecule induce dimerization of GST-FKBP12^F36V^ and His-p300-BRD. **c,** Chemical structures of terminal binding molecules. FKBP-c is used as a competitive binder FKBP12^F36V^ and p300-c is used as a competitive binder for p300/CBP.

**Supplementary Figure 2.**
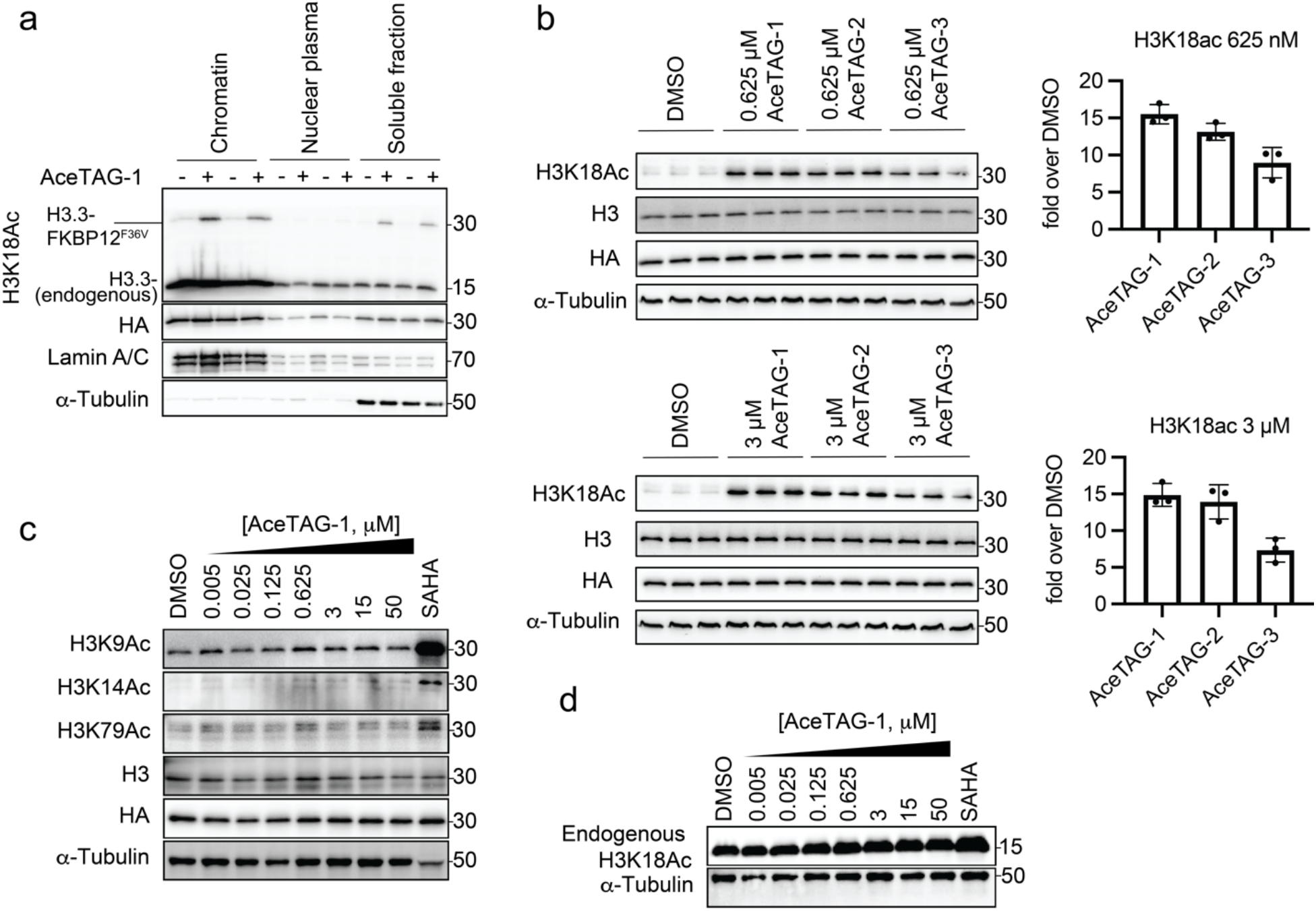
Confirmation of Histone H3.3-FKBP12^F36V^ acetylation induced by AceTAGs. **a,** Cell fractionation confirms localization of Histone H3.3-FKBP12^F36V^ to chromatin. H3.3-FKBP12^F36V^ HeLa cells were treated with AceTAG-1 (625 nM) or DMSO for 2h. Acetylation of K18 was monitored by immunoblot. **b,** Comparison of K18ac level induced by different AceTAGs. H3.3-FKBP12^F36V^ HeLa cells were treated with 625 nM or 3 μM AceTAG1-3 for 2h. Acetylation of K18 was monitored by immunoblot analysis of lysate. Shown in the right panel is quantitation of immunoblot signal of K18Ac relative to α-tubulin as the mean ± s.e.m. of *n*=3 biologically independent experiments. **c,** Acetylation of K9, K14 and K79 were not induced by AceTAG-1. H3.3-FKBP12^F36V^ HeLa cells were treated with increasing concentrations of AceTAG-1 for 2h. Acetylation of K9, K14 and K79 were monitored by immunoblot analysis of lysate. **d,** Endogenous Histone H3K18 acetylation was not affected by AceTAG-1. H3.3-FKBP12^F36V^ HeLa cells were treated with increasing concentrations of AceTAG-1 for 2h. Acetylation of K18 was monitored by immunoblot analysis of lysate. See Supplementary Figure 12 and 13 for full blot images.

**Supplementary Figure 3.**
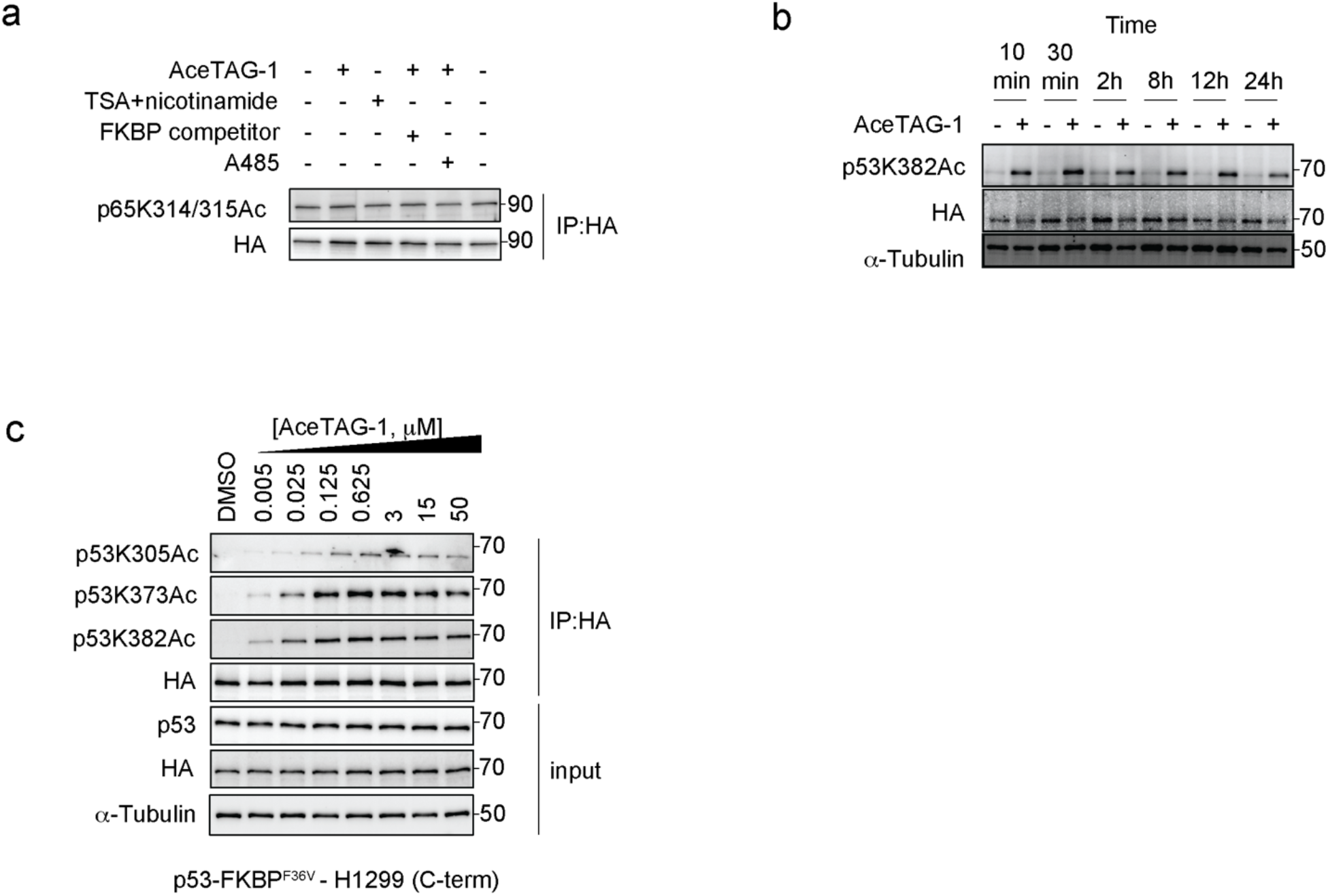
AceTAG-1 p53 kinetics and induction of acetylation of C-terminally tagged p53. **a**, Immunoblot analysis of controls for AceTAG-mediated acetylation of RelA. Cells were treated with TSA (2 μM) and nicotinamide (10 mM) as a positive control for p65 acetylation^45^ or co-treated with AceTAG-1 (625 nM) and DMSO, an FKBP12^F36V^ binding ligand (Supplementary Figure 1c), or the p300/CBP KAT domain inhibitor A485 for 2 h which block AceTAG-1-induced acetylation. **b,** Kinetics of AceTAG-1 induced acetylation on FKBP12^F36V^-p53. FKBP12^F36V^-p53 H1299 cells were treated with AceTAG-1 (625 nM) or DMSO for indicated time. Acetylation of K382 was monitored by immunoblot analysis of cell lysate. **c,** AceTAG-1 induces dose-dependent acetylation on K382, K373 and K305 of p53-FKBP12^F36V^. p53-FKBP12^F36V^ H1299 cells were treated with increasing concentrations of AceTAG-1 for 2h, lysed and p53-FKBP12^F36V^ was enriched and blotted with noted antibodies. The results in **a-c** are representative of two independent biological replicates (*n*=2). See Supplementary Figure 14 for full blot images.

**Supplementary Figure 4.**
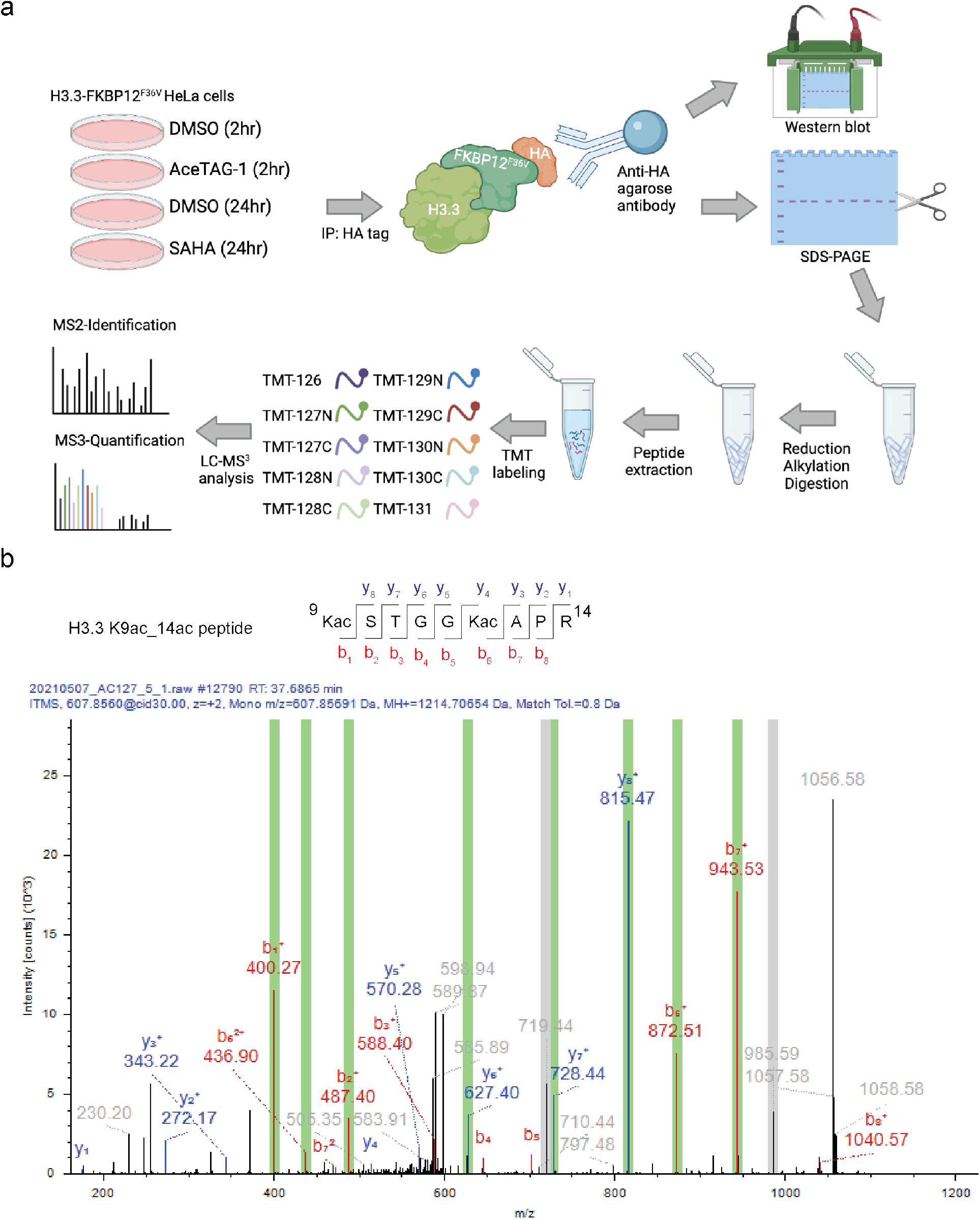
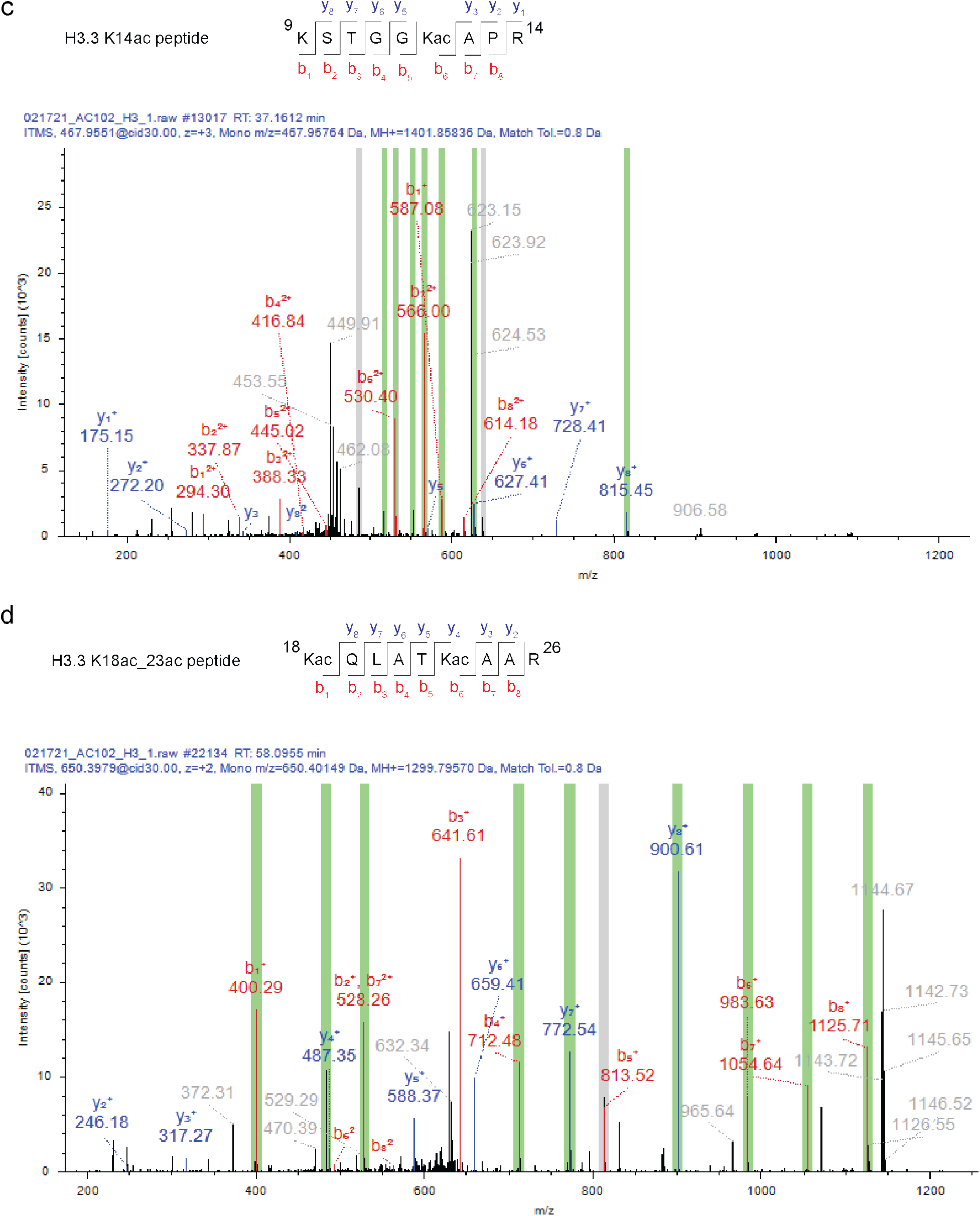
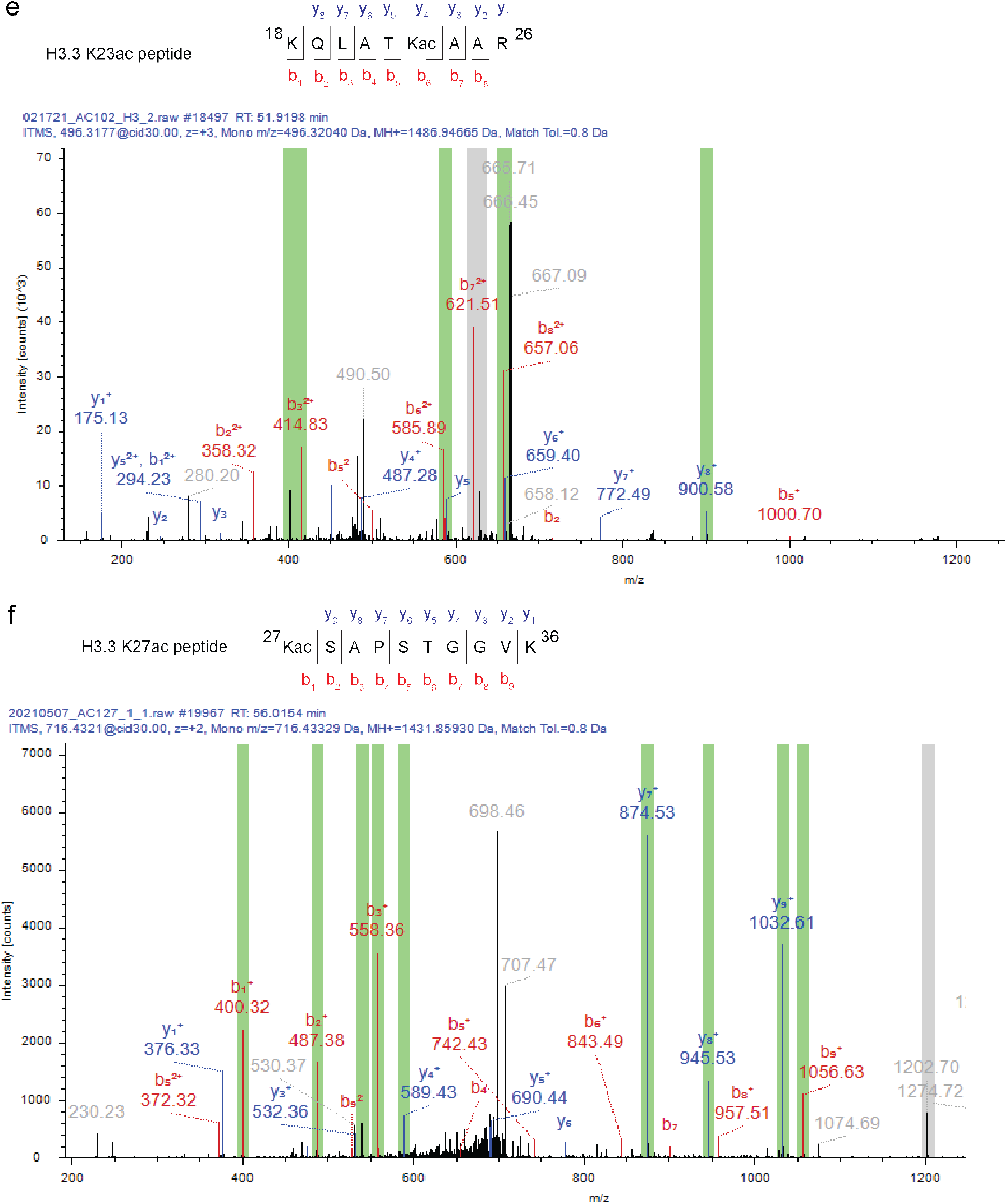
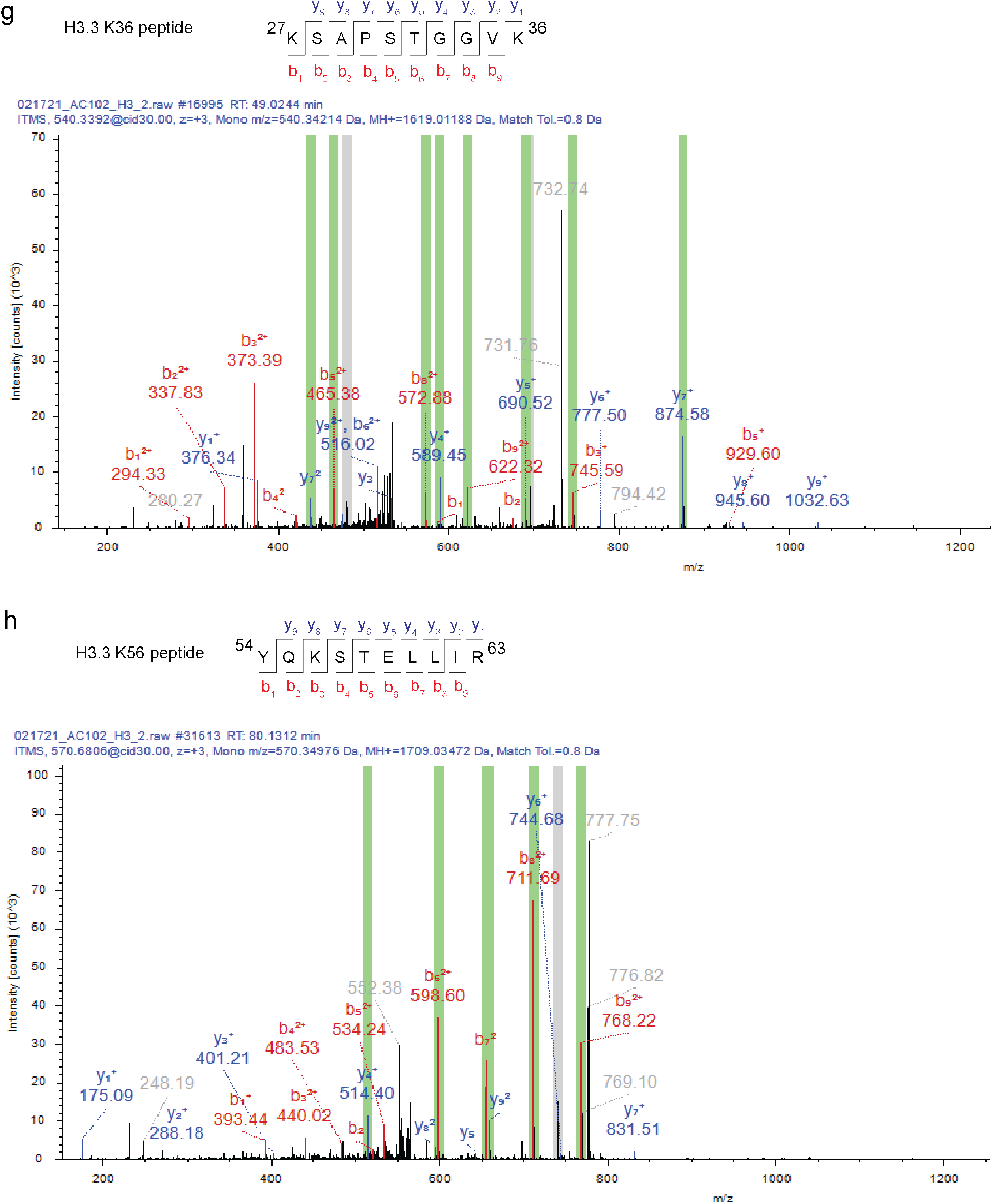
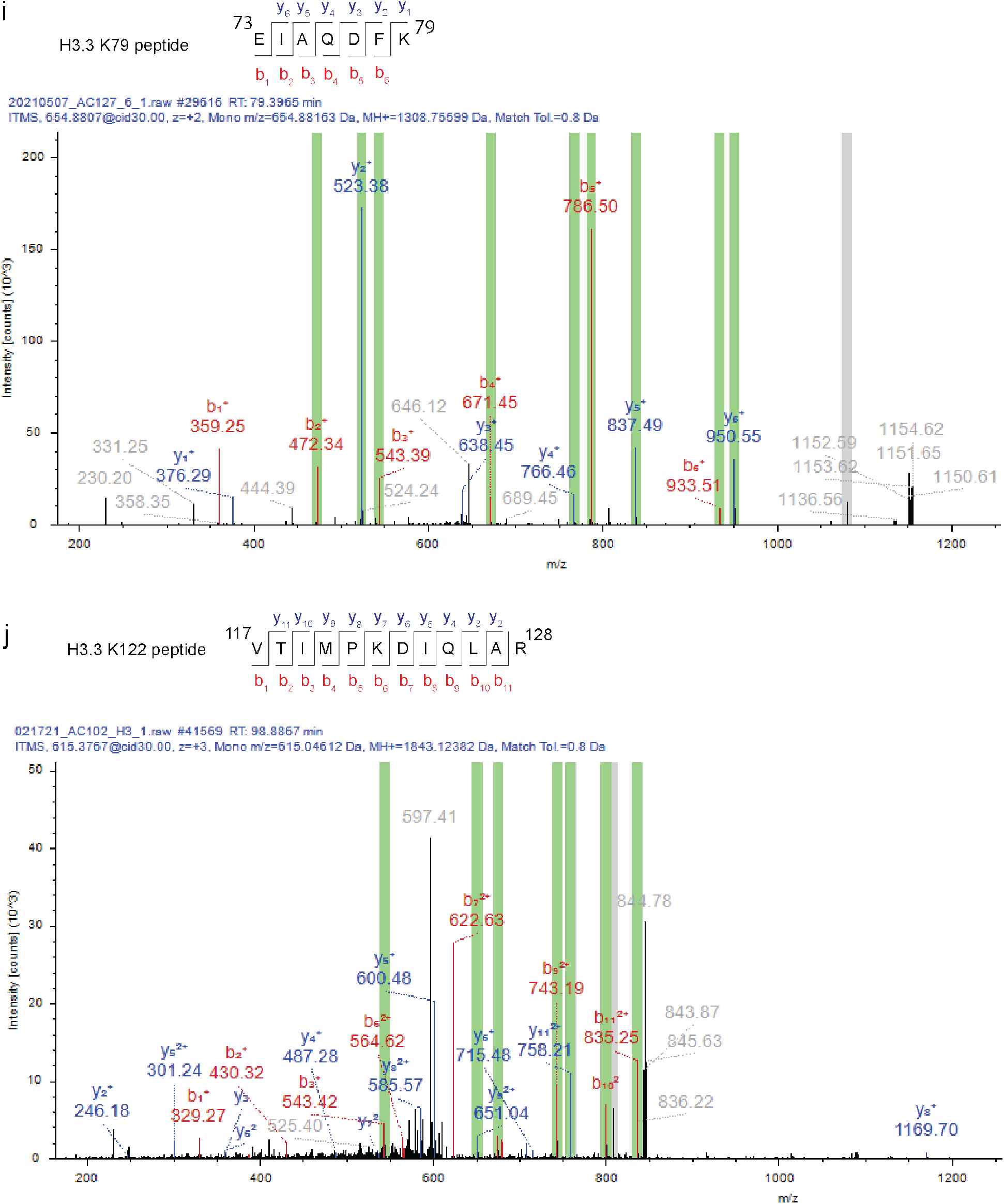
Confirmation of H3.3-FKBP12^F36V^ acetylation in cells by quantitative mass spectrometry. **a,** Schematic depiction of H3.3-FKBP12^F36V^ acetylation determination using TMT-based quantitative proteomics and immunoblot. H3.3-FKBP12^F36V^ HeLa cells were treated with DMSO, 625 nM AceTAG-1 and 5 μM SAHA for 2h or 24h, respectively. H3.3-FKBP12^F36V^ was enriched by immunoprecipitation with HA antibody conjugated beads. Eluted protein was resolved by immunoblot analysis or SDS-PAGE. Gel slices containing H3.3-FKBP12^F36V^ were subjected for in-gel trypsin digestion. Recovered peptides were labeled with TMT10plex Isobaric Label Reagent and analyzed by LC-MS/MS/MS proteomics. Data were processed and analyzed with Proteome Discoverer 2.4. **b,** MS2 spectrum of H3.3K9ac/K14ac peptide. **c,** MS2 spectrum of H3.3K14ac peptide. **d,** MS2 spectrum of H3.3K18ac/K23ac peptide. **e,** MS2 spectrum of H3.3K23ac peptide. **f,** MS2 spectrum of H3.3K27ac peptide. **g,** MS2 spectrum of H3.3K36 peptide. **h,** MS2 spectrum of H3.3K56 peptide. **i,** MS2 spectrum of H3.3K79 peptide. **j,** MS2 spectrum of H3.3K122 peptide.

**Supplementary Figure 5.**
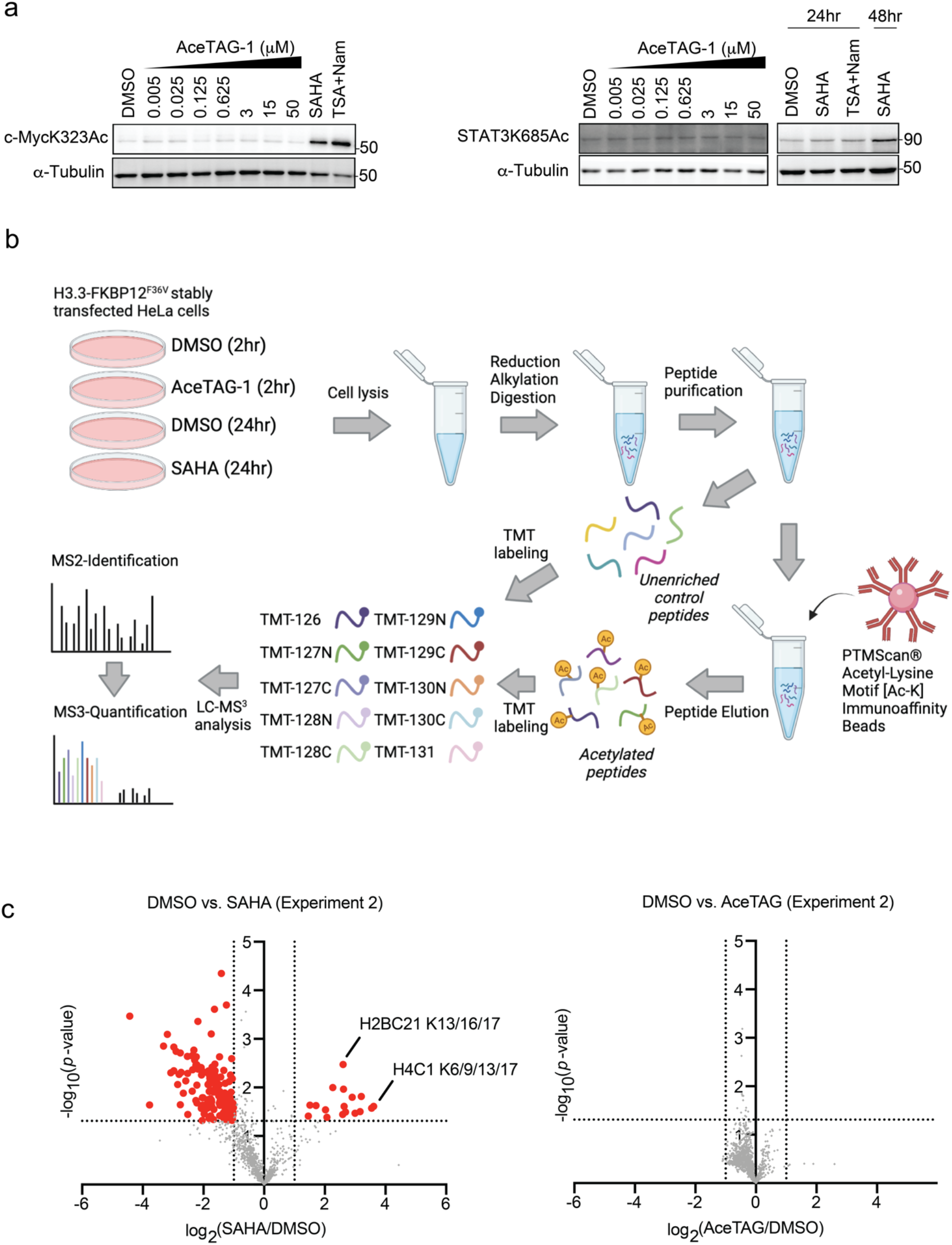
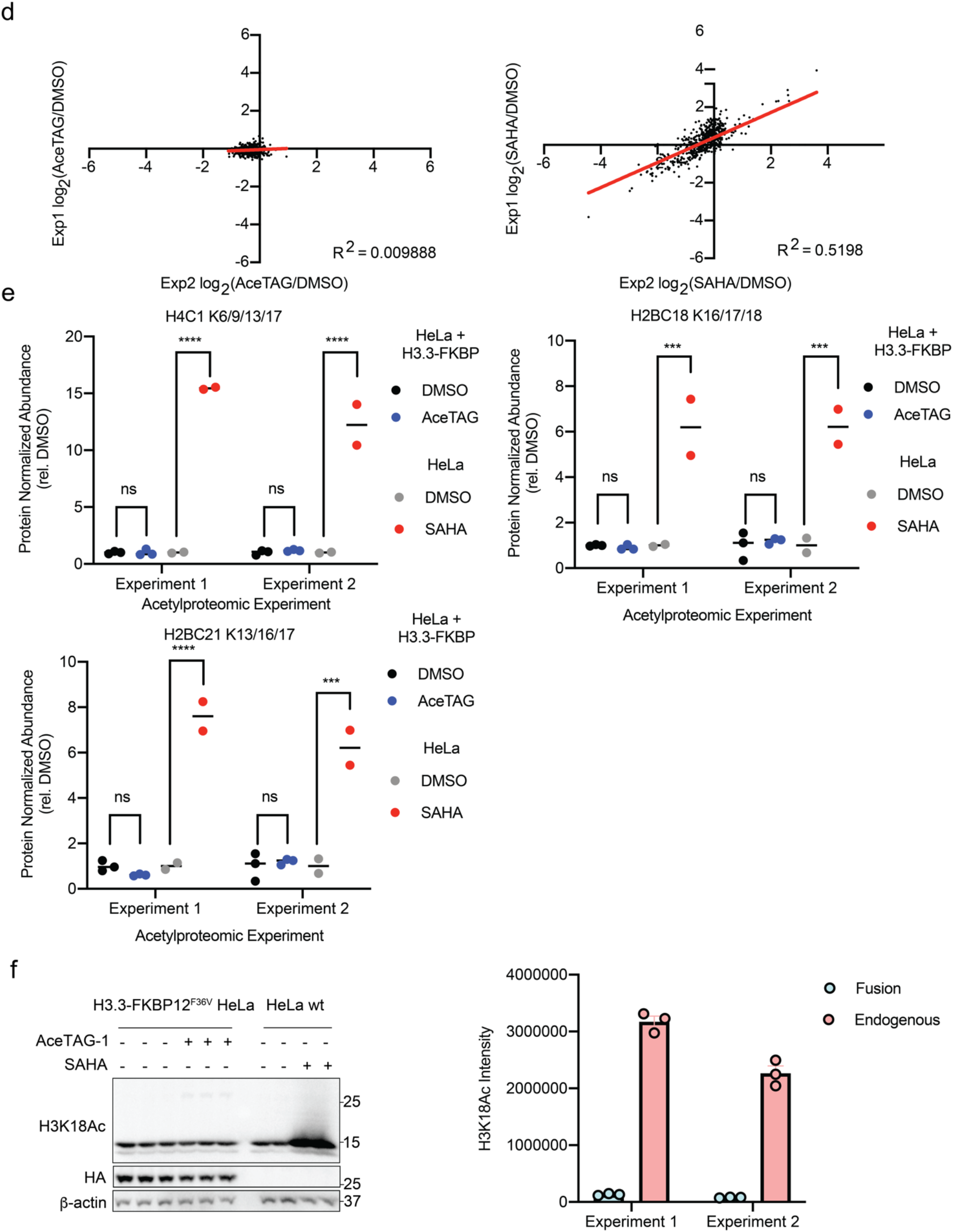
Confirmation of proteome-wide selectivity of AceTAG induced acetylation. **a,** No acetylation increase was observed for established p300/CBP targets including c-Myc and STAT3. H3.3-FKBP12^F36V^ HeLa cells were treated with increasing concentrations of AceTAG-1 for 2h. Acetylation of c-myc K323ac and STAT3 K685ac was monitored by immunoblot analysis. **b,** Schematic depiction of quantitative proteomics experiments to profile global acetylation changes. H3.3-FKBP12^F36V^ HeLa cells were treated with (1) DMSO, (2) 600 nM AceTAG-1 for 2h or (3) 2 μM SAHA for 16h. Acetyl peptides were enriched from cell lysates using PTMscan acetyl-lysine motif immunoaffnity beads. Both acetyl peptides and unenriched peptides were labeled with TMT10plex isobaric tags and analyzed by LC-MS/MS/MS quantitative proteomics. Data were processed and analyzed with Proteome Discoverer 2.4. **c,** Volcano plot of independent TMT experiment of AceTAG-1 or SAHA induced acetylproteome changes. H3.3-FKBP12^F36V^ HeLa cells were treated with DMSO, AceTAG-1 (600 nM) or SAHA (2 μM) for 2 or 16 h respectively, lysed, trypsinized and acetylated peptides enriched and labeled with TMT reagents, then subjected to quantitative MS. The vertical dashed lines correspond to 2-fold change in enrichment relative to DMSO and the horizontal line corresponds to a *P*-value of 0.05 for statistical significance. Red circles correspond to protein targets with >2-fold change (*P* < 0.05) relative to DMSO. **d,** Scatter plot of independent experiments 1 and 2 of AceTAG or SAHA induced acetylproteome changes. **e,** Selected histone acetylpeptides demonstrating minimal off-target effects of AceTAG. Significance was determined using two-way ANOVA with Šídák’s multiple comparisons test (ns - not significant, \*\*\**p*<0.001, \*\*\*\**p* < 0.0001). **f,** H3.3-FKBP12^F36V^ acetylation and expression levels are lower relative to endogenous histones. H3.3-FKBP12^F36V^ HeLa cells or HeLa cells were treated with DMSO, AceTAG (600 nM) or SAHA (2 μM) for 2 or 16 h respectively. Acetylation of H3.3K18ac-FKBP12^F36V^ and endogenous H3K18ac were monitored by immunoblot analysis. Shown in the right panel is quantitation of immunoblot signal of H3.3K18ac-FKBP12^F36V^ and endogenous H3K18ac as the mean ± s.e.m. of *n*=3 biologically independent experiments. See Supplementary Figure 15 for full blot images.

**Supplementary Figure 6.**
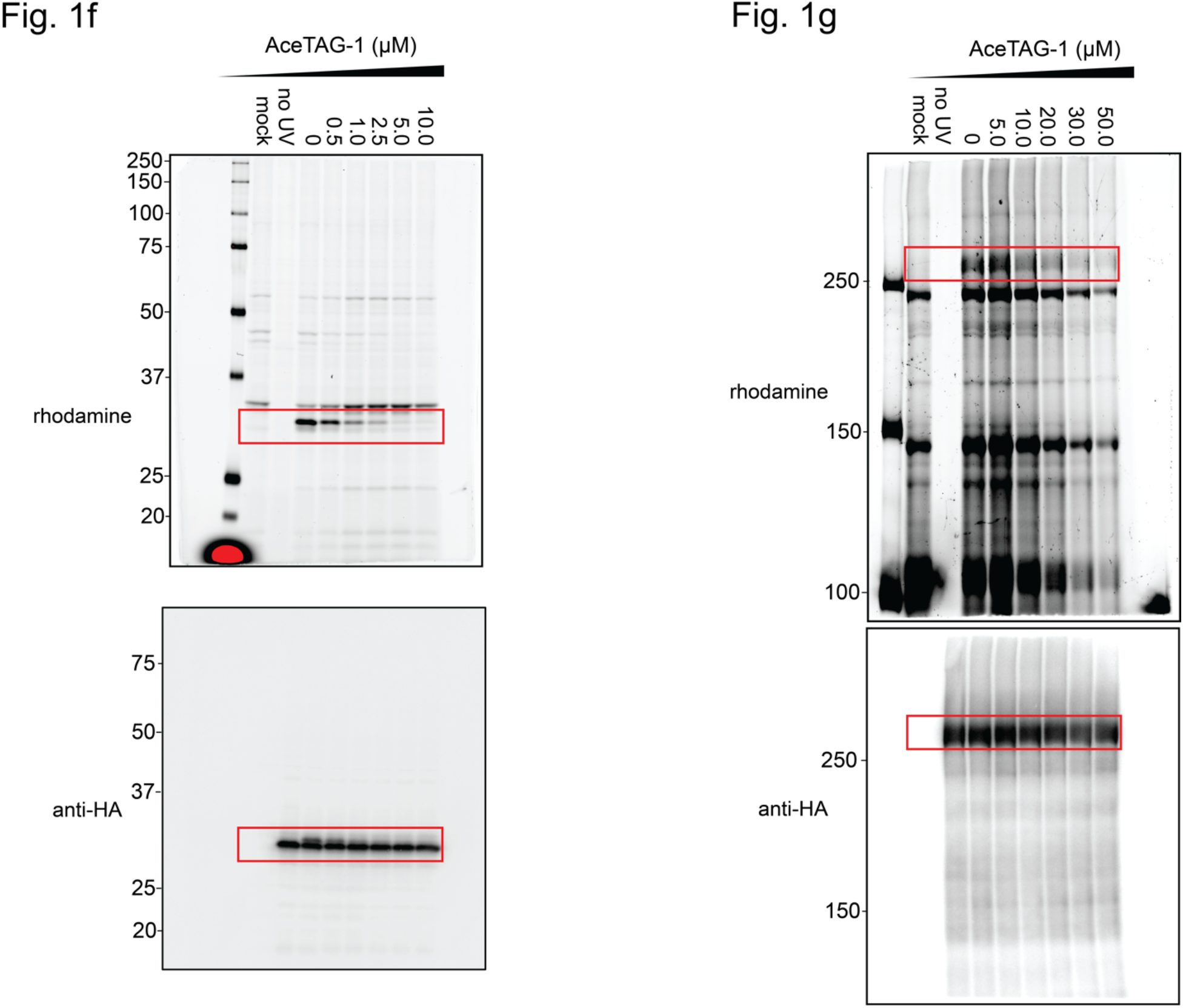
Uncropped gels and immunoblots for Figure 1f and Figure 1g.

**Supplementary Figure 7.**
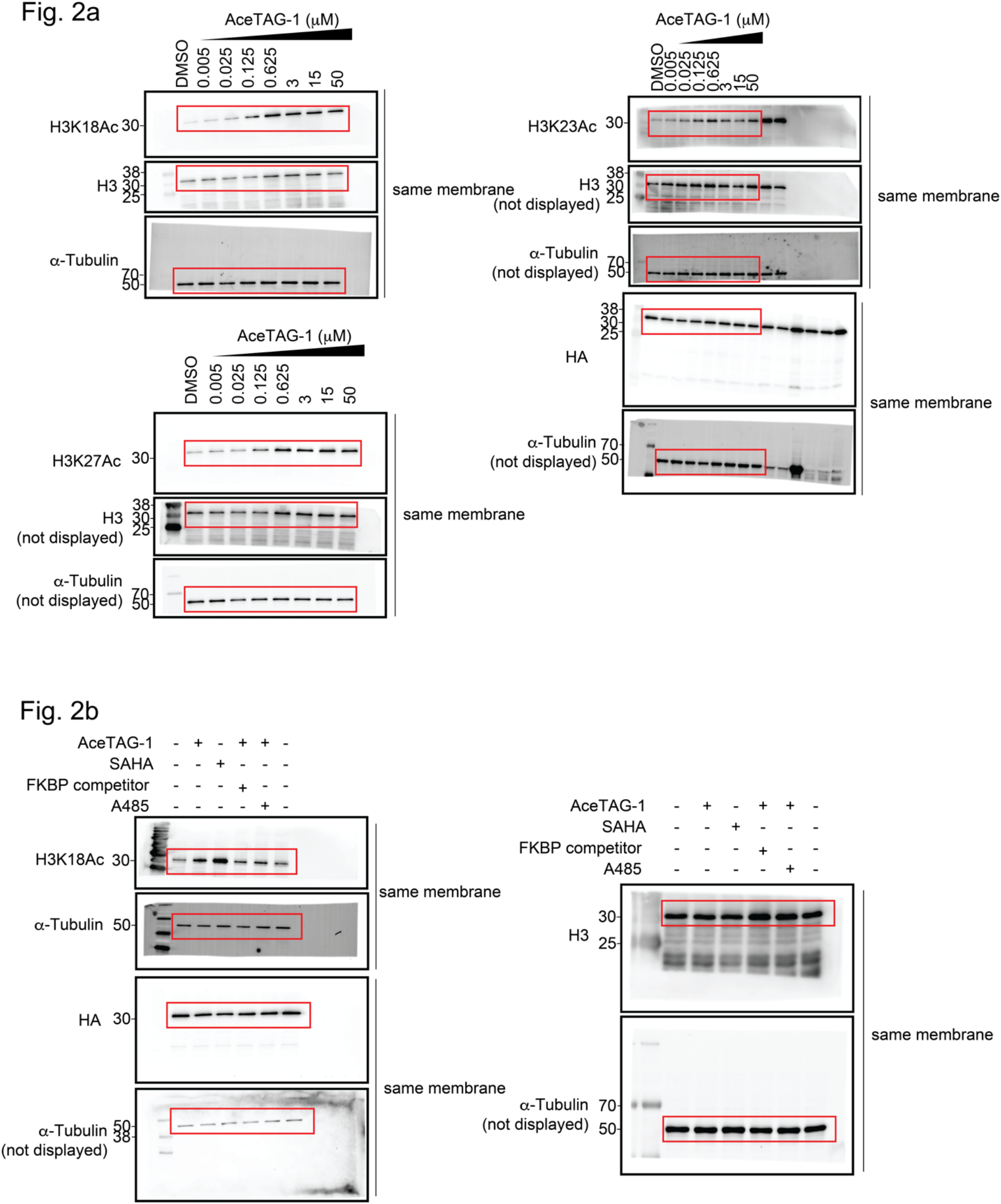
Uncropped immunoblots for Figure 2a and Figure 2b.

**Supplementary Figure 8.**
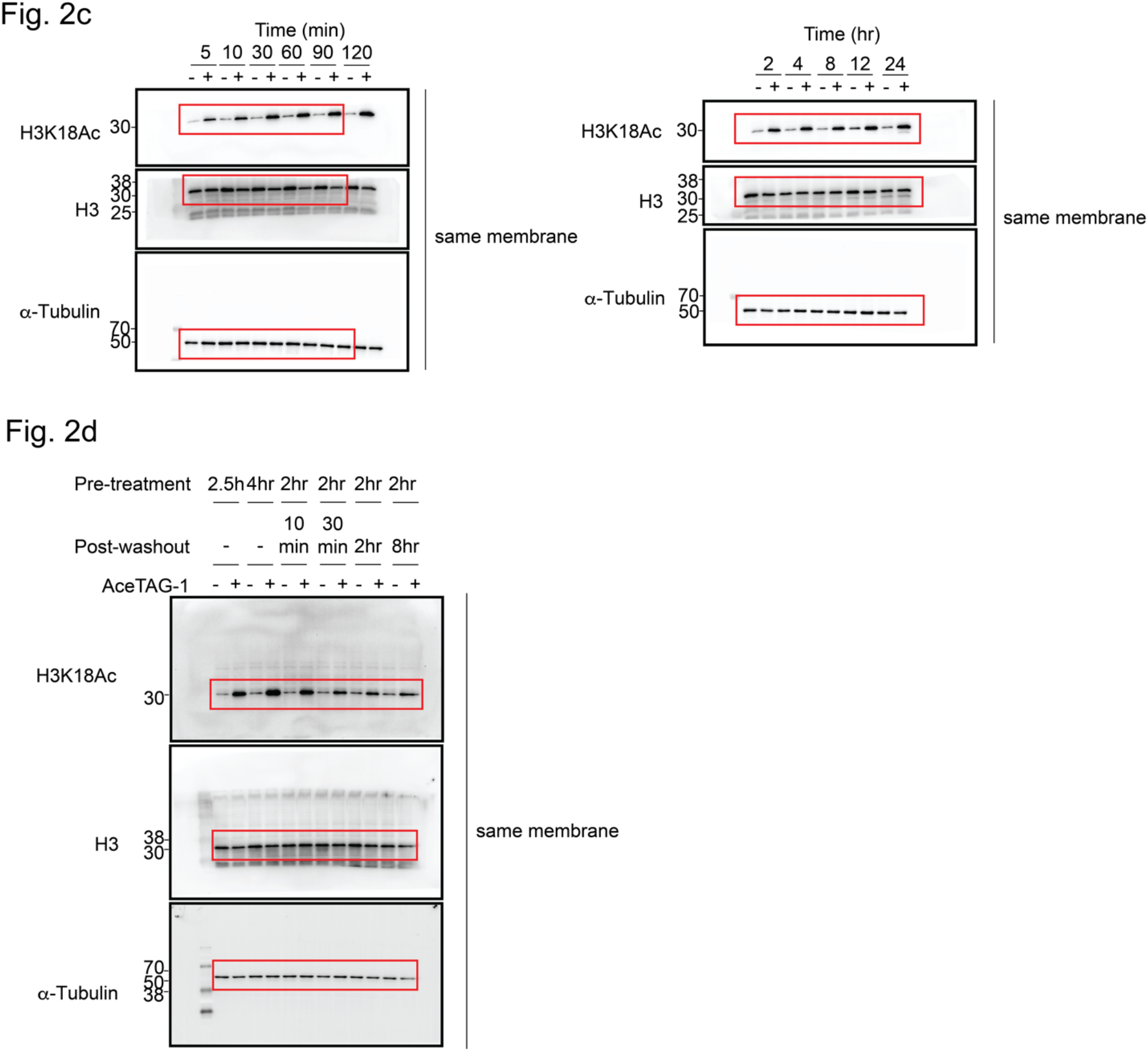
Uncropped immunoblots for Figure 2c and Figure 2d.

**Supplementary Figure 9.**
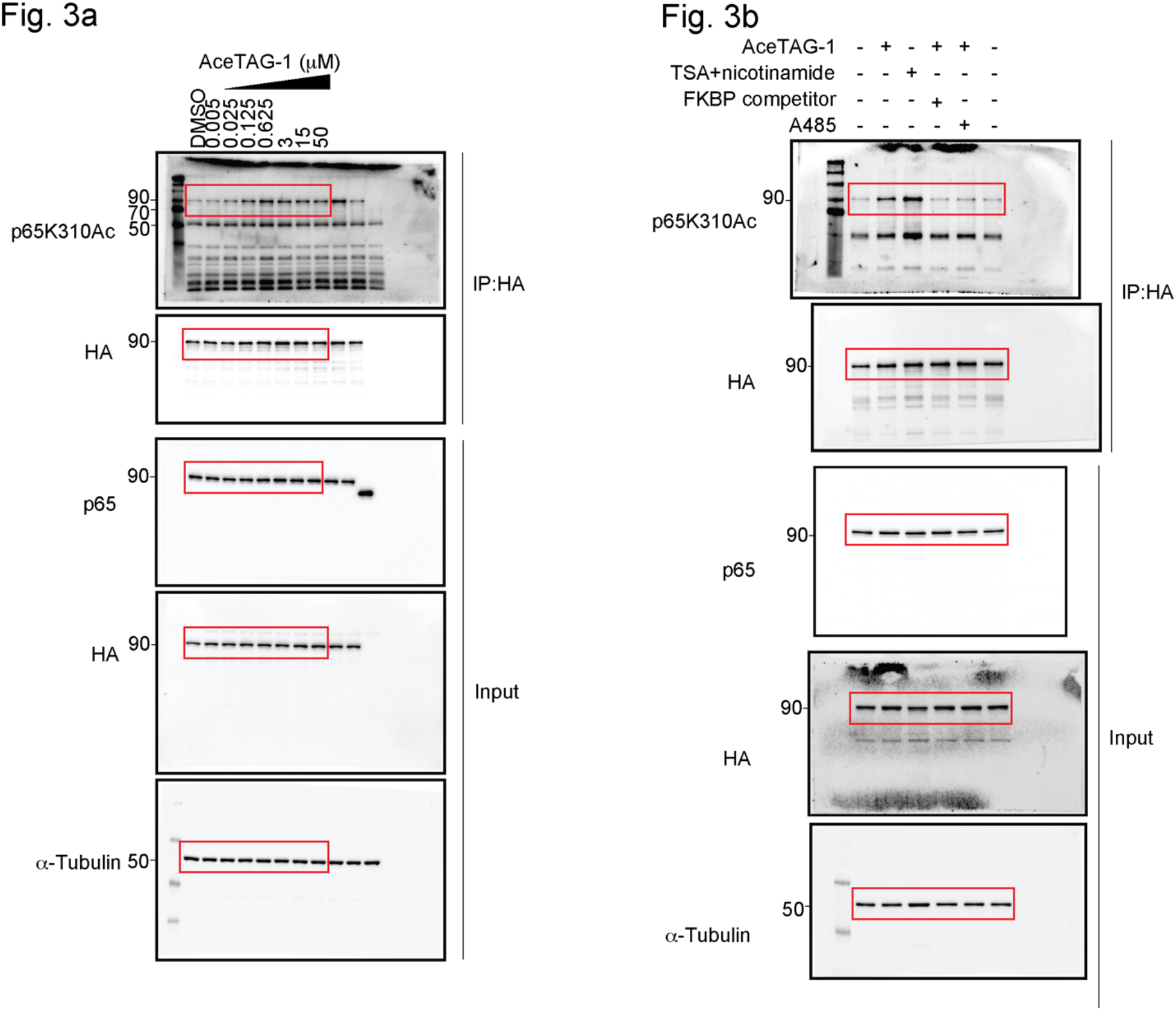
Uncropped immunoblots for Figure 3a and Figure 3b.

**Supplementary Figure 10.**
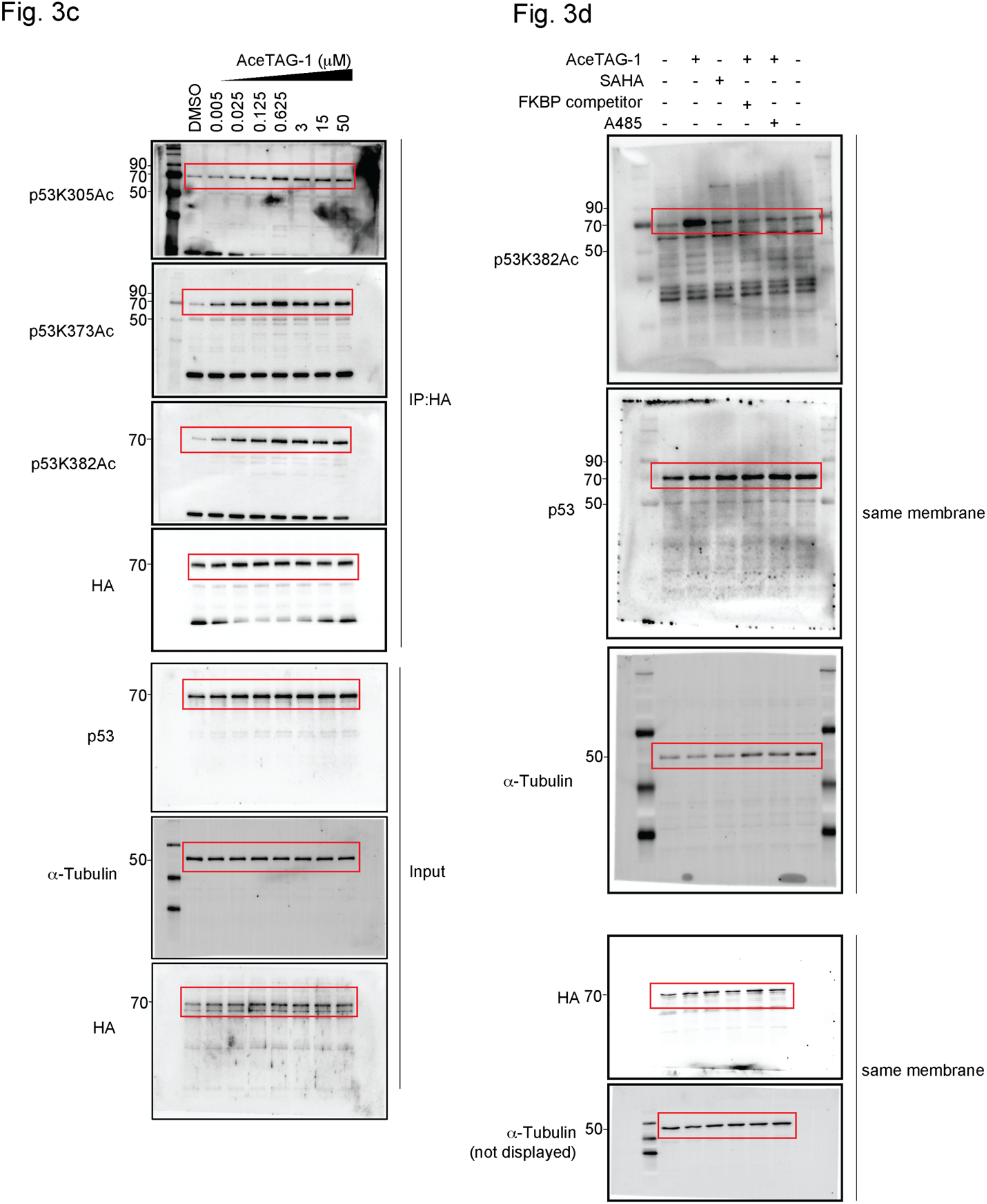
Uncropped immunoblots for Figure 3c and Figure 3d.

**Supplementary Figure 11.**
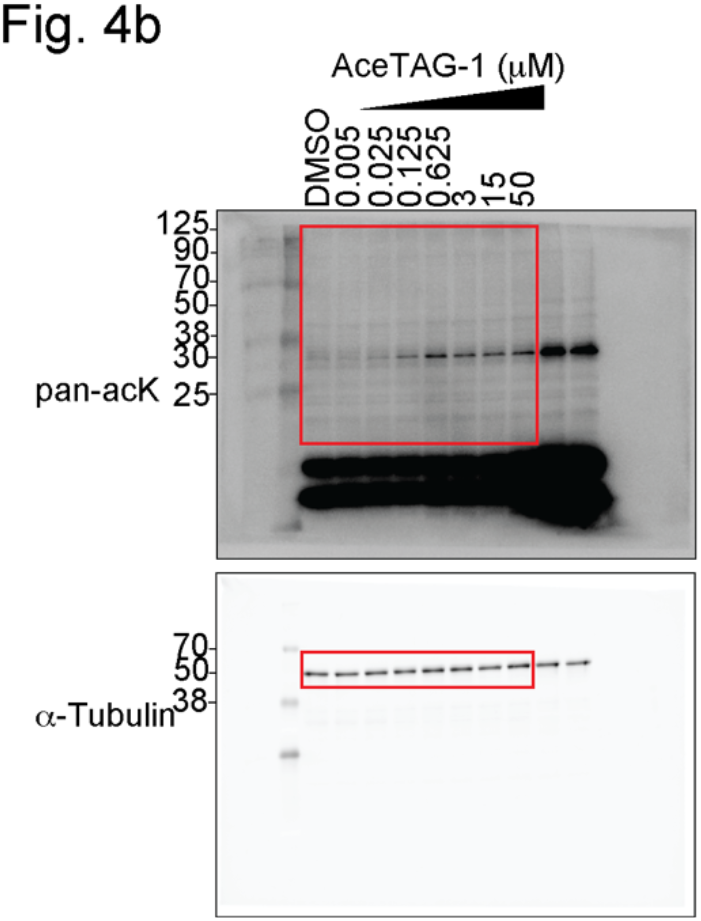
Uncropped immunoblots for Figure 4b.

**Supplementary Figure 12.**
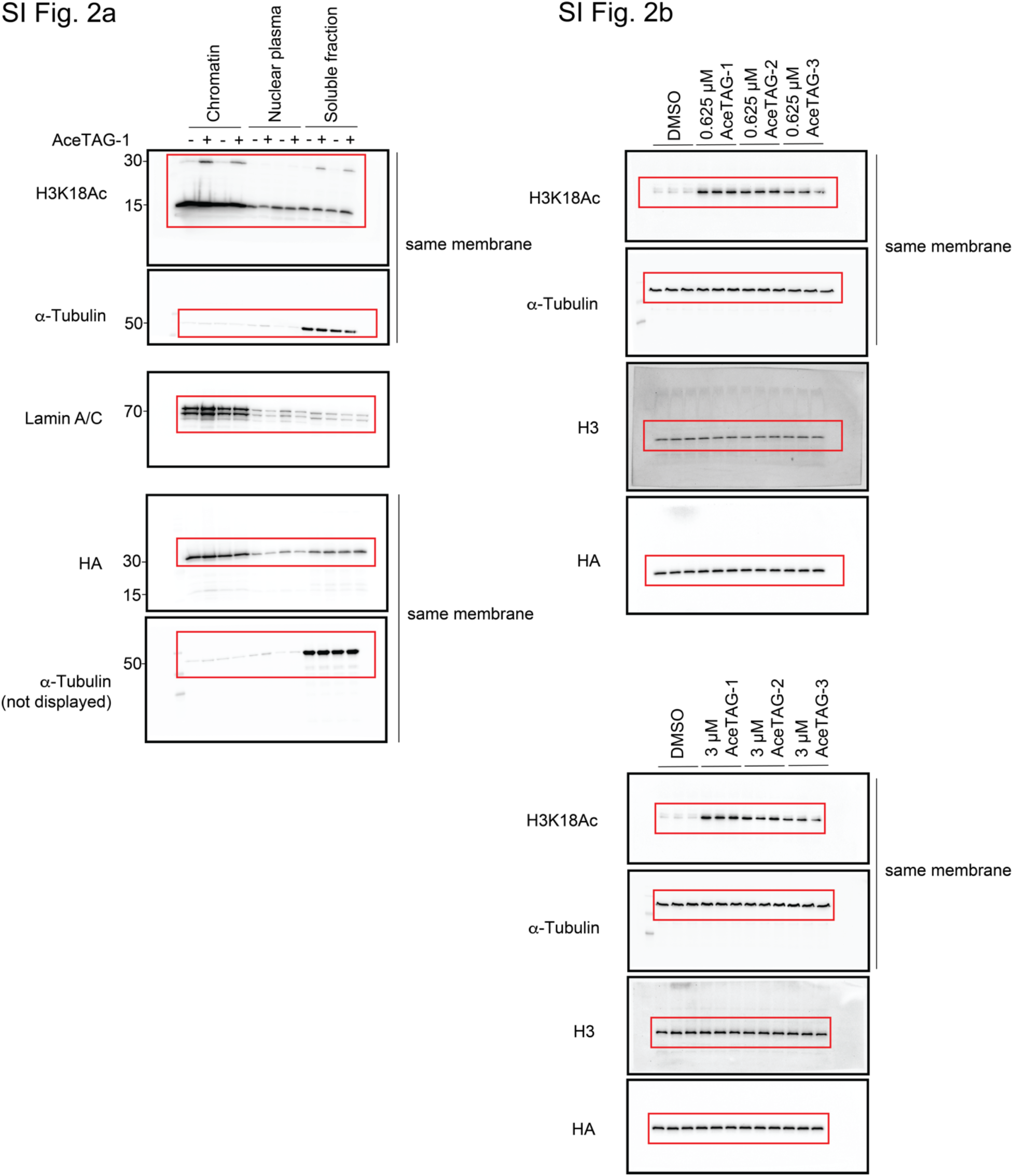
Uncropped immunoblots for SI Figure 2a and SI Figure 2b.

**Supplementary Figure 13.**
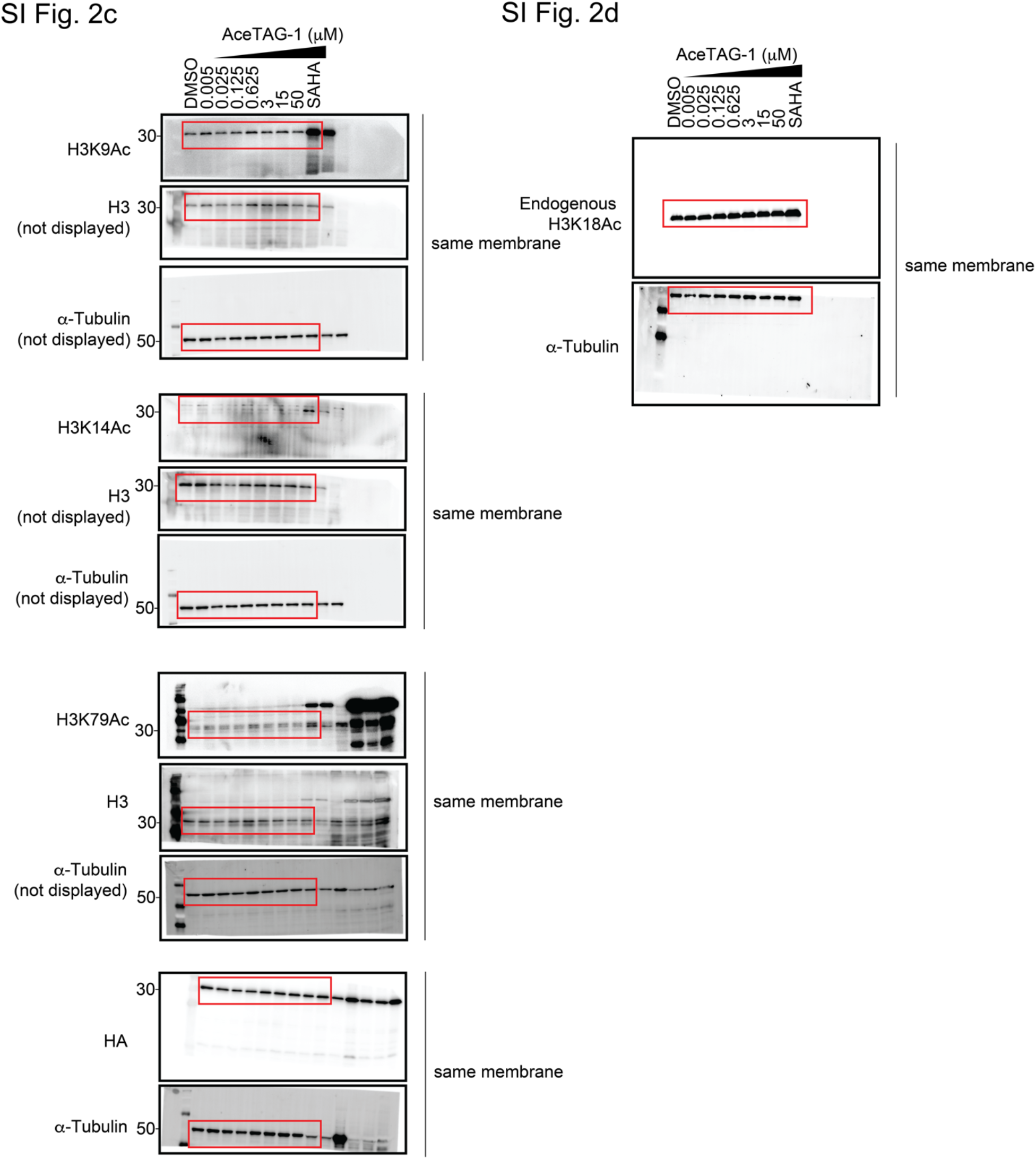
Uncropped immunoblots for SI Figure 2c and SI Figure 2d.

**Supplementary Figure 14.**
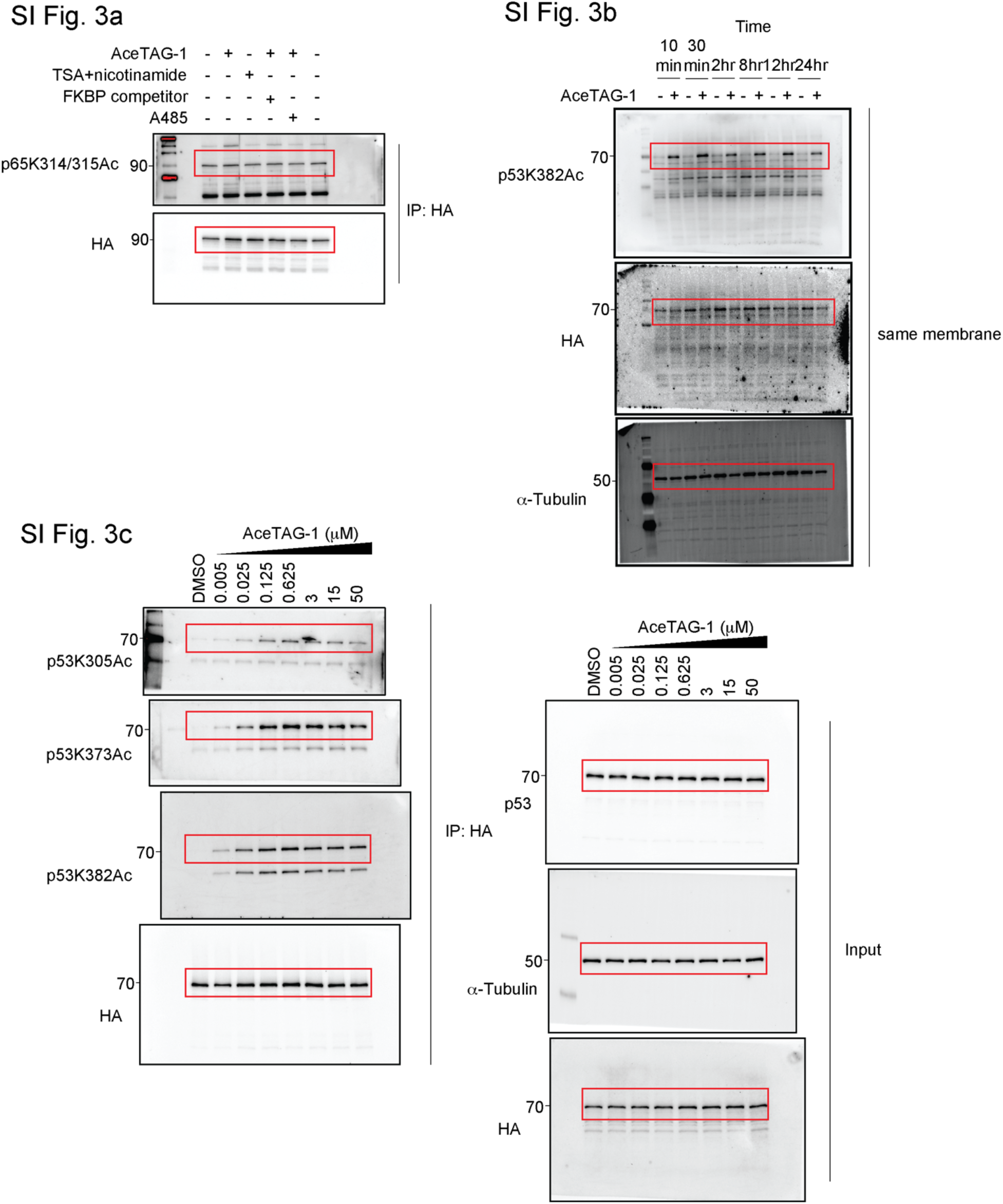
Uncropped immunoblots for SI Figure 3a, SI figure 3b, and SI Figure 3c.

**Supplementary Figure 15.**
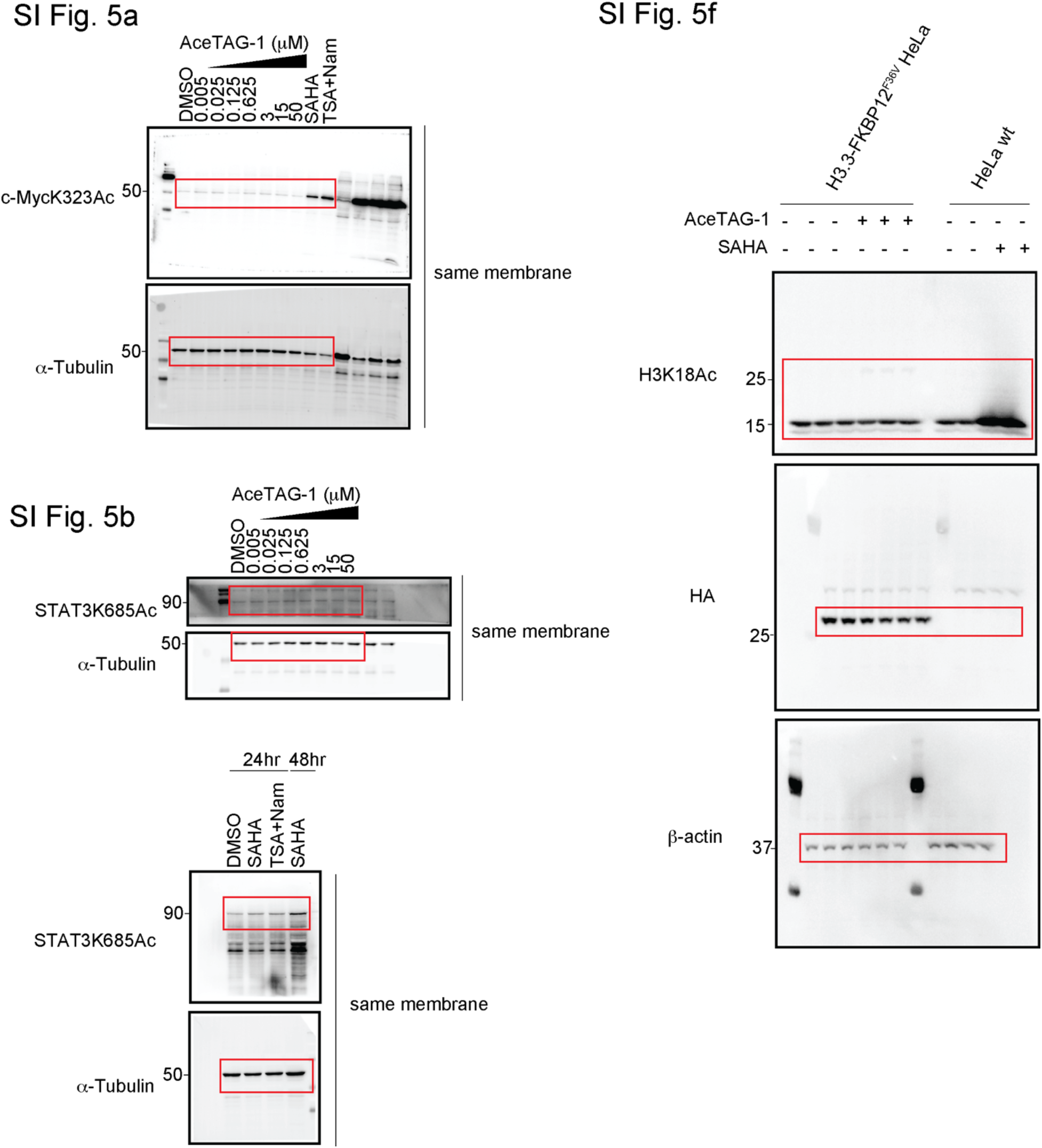
Uncropped immunoblots for SI Figure 5a, SI figure 5b, and SI Figure 5f.

## MATERIALS AND METHODS

### Materials

Mouse Anti-HA (2367S, 1:2000 dilution), rabbit anti-HA (3724S, 1:2000 dilution), rabbit anti-H3K9ac (9649S, 1:2000 dilution), rabbit anti-H4K14ac (7627S, 1:2000 dilution), rabbit anti-H3K18ac (13998S, 1:2000 dilution), rabbit anti-H3K27ac (8173S, 1:2000 dilution), rabbit anti-H3 (4499S, 1:2000 dilution), rabbit anti-acetyl lysine (9814S, 1:1000 dilution), rabbit anti-NFκBp65 (8242S, 1:2000 dilution), rabbit anti-p53K382ac (2525S, 1:1000 dilution), rabbit anti-Stat3K685ac (2523S, 1:2000 dilution), and anti-rabbit IgG HRP conjugate (7074P2, 1:10000 dilution) were from Cell Signaling Technology. Anti-tubulin hFAB Rhodamine (12004166, 1:3000 dilution) was from Bio-rad). Rabbit anti-H3K23ac (07-335, 1:2000 dilution), rabbit anti-H3K79 (SAB5600231-100UG, 1:2000 dilution), rabbit anti-c-MycK323ac (ABE26, 1:2000 dilution) were from Millipore Sigma. Rabbit anti-p65K310ac (ab19870, 1:1000 dilution), Rabbit anti-p65K314/315ac (PA5-114696, 1:500 dilution), rabbit anti-p53K373ac (ab62376, 1:1000 dilution), and rabbit anti-p53K305ac (ab109396, 1:1000 dilution) were from Abcam. Mouse anti-p53 (628202, 1:2000 dilution) was from Biolegend. Anti-mouse IgG (H+L) HRP conjugate (PA1-28568, 1:10000 dilution) was from Thermo Fisher Scientific.

### Cell Culture

HEK293T cells (ATCC) were cultured in DMEM supplemented with 10% FBS, 1% (v/v) penicillin/streptomycin, and 2 mM glutamine. H1299 (ATCC) were cultured in RPMI supplemented with 10% FBS, 1% (v/v) penicillin/streptomycin, 2 mM glutamine, 10 mM HEPES, and 1 mM sodium pyruvate. HeLa cells (ATCC) were cultured in DMEM supplemented with 10% FBS, 1% (v/v) penicillin/streptomycin, and 2mM glutamine. RelA KO HeLa cells (Abcam) we cultured in DMEM supplemented with 10% FBS, 1% (v/v) penicillin/streptomycin, and 2 mM glutamine. All cells were cultured at 37°C and 5% CO_2_.

### Lentivirus Plasmid Construction

pLEX_305-N-dTAG and pLEX305-C-dTAG empty vectors were kindly provided by Dr. Erb (The Scripps Research Institute). Gateway recombination cloning technology (Invitrogen) was used to clone targets of interest into pLEX_305-N-dTAG and pLEX_305-C-dTAG. H3.3 cDNA ORF clone in pGEM-T vector was purchased from SinoBiological (Cat# HG16451-G). p53 cDNA ORF clone in pCMV3-C-HA vector was purchased from SinoBiological (Cat# HG10182-CY). RelA cDNA ORF clone was purchased from SinoBiological (Cat# HG12054-G). H3.3, p53, and RelA were cloned into Gateway compatible donor vector pDONR221 using BP clonase after PCR with primers containing BP overhangs sequence from the vector described above. H3.3 was cloned into pLEX_305-C-dTAG and p53 as well as RelA were cloned into pLEX_305-N-dTAG using Gateway LR clonase II Enzyme mix kit (Invitrogen). The sequences of the primers are listed below.

Briefly, 100 ng pDONR221 vectors containing genes of interest were mixed with 150 ng of pLEX_305-C-dTAG or pLEX_305-N-dTAG in TE buffer. 1 μl of LR clonase II enzyme was added to the plasmid mix and incubated for 1h at room temperature, followed by addition of 1 μl Proteinase K and incubated for 10 min at 37°C to terminate the reaction. Samples were transformed using DH10B competent cells and plated on Ampicillin selective agar plates. After growing at 30°C overnight, colonies were picked and grown up in 5mL LB media supplemented with Ampicillin at 30°C overnight. Plasmid DNA was purified using Zyppy Plasmid Miniprep Kit (Genesee Scientific) and sequenced using primers as followed: pLEX_305-dTAG-seq-F: TGTTCCGCATTCTGCAAGCCTC and pLEX_305-dTAG-seq-R ACAAAGGCATTAAAGCAGCGTATCC.

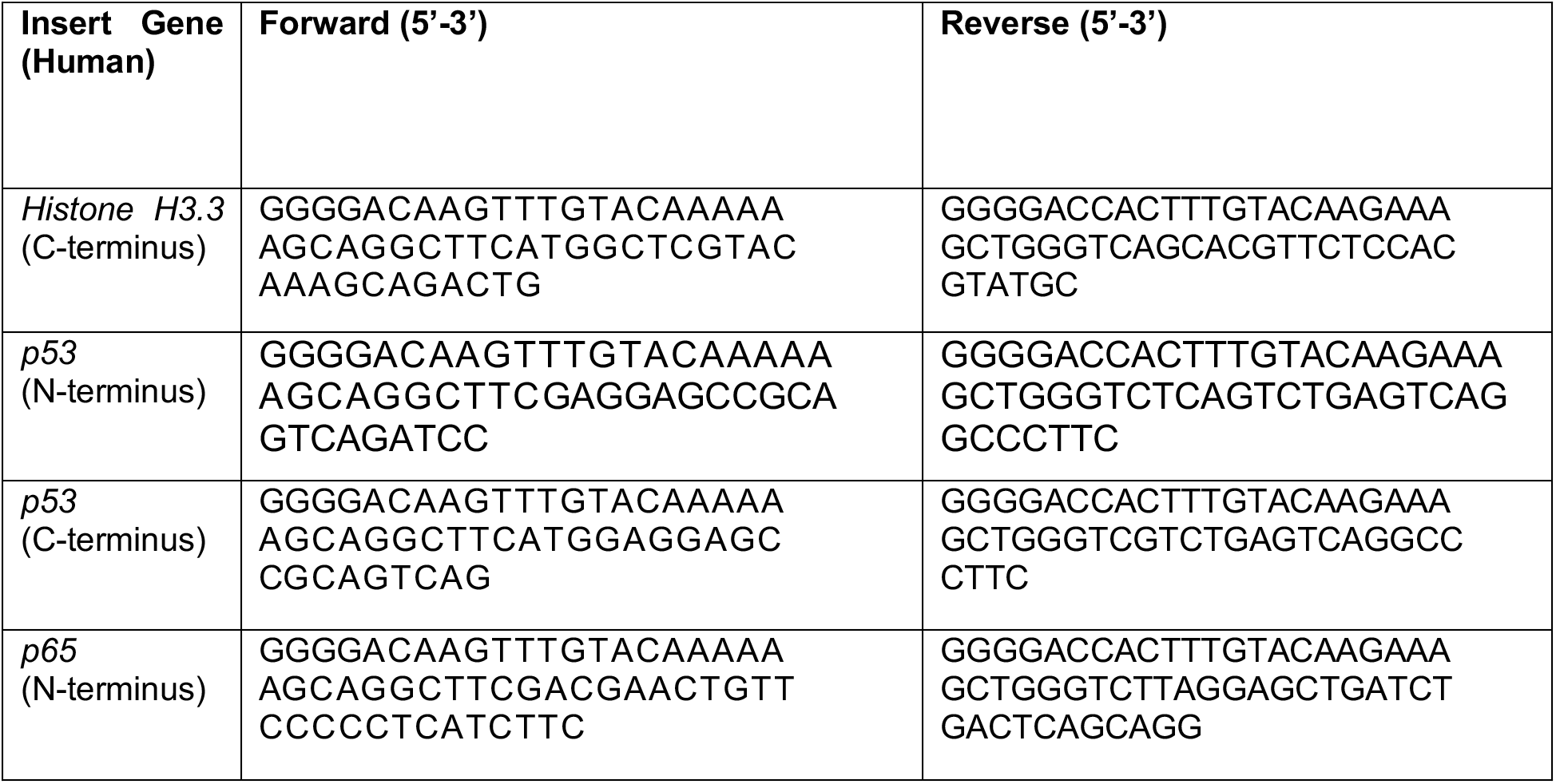

### Lentivirus Production and Transduction

Lentiviral production was performed using HEK293T cells, which were co-transfected with pMD2.G (Addgene, #12259), psPAX2 (Addgene, #12260), and AceTAG lentiviral plasmids using PEI (Polysciences Inc). Viral particles were collected 72 h after transfection and filtered through a 0.45 μM membrane. A range of dilutions of the lentivirus in DMEM complete and 10 µg/mL polybrene were added to HeLa, H1299, or HeLa RelA KO cell lines. Transduced cells were selected with 1 µg/mL or 2 µg/mL puromycin.

### Protein Expression

A construct containing residues 2-108 of FKBP12^F36V^ in pGEX4t-1 vector was overexpressed in E. coli BL21(DE3) in LB medium in the presence of 100 mg/mL carbenicillin. Cells were grown at 37°C to an OD of 0.6, induced with 100 μM isopropyl-1-thio-D-galatopyranoside (IPTG), incubated for 4h at 37°C, and collected by centrifugation. Cell pellets were suspended in lysis buffer (50 mM HEPES pH 8.0, 250 mM NaCl, 0.1%(v/v) Triton X-100, 1 mM TCEP), sonicated and then cleared by centrifugation at 11,000 g for 40 min at 4°C. FKBP12^F36V^ was first purified with Pro Affinity Concentration Kit GST (Amicon: ACR5000GS), and further purified on HiPrep 16/60 Sephacryl S-200 HR gel filtration column. Clean fraction was concentrated and buffer exchanged to storage buffer (50 mM HEPES, 250 mM NaCl, pH 8.0, 10% Glycerol), protein concentration was measured with Thermo 660nM kit, flash freezed and stored at −80°C.

### GST-FKBP12^F36V^/His-P300-BRD AlphaScreen Assay

GST-FKBP12^F36V^ and 6xHis-p300-BRD (active motif #81158) were diluted to 250 nM and 500 nM, in assay buffer (50 mM HEPES pH 7.4, 200 mM NaCl, 1 mM TCEP, and 0.1% BSA), 20 μL of protein mixture was added to each well of a 1/2 area 96-well OptiPlate (PerkinElmer). Compounds were then added from DMSO stock to protein mixture (0.4 μl/well) with 1% DMSO in final mixture. Plates were then shaken orbitally at room temperature for 1h. Nickel Chelate AlphaLISA Acceptor and Glutathione AlphaLISA Donor beads (PerkinElmer) diluted to 20 ng/μl in assay buffer and 20 μl beads mixture was added to each well. After a 1h orbital shaking at room temperature. lluminescence was measured on the Envision 2104 plate reader (PerkinElmer). Data were plotted and analyzed using GraphPad PRISM v8 and fit with ‘Bell-shaped’ dose response curve.

For ternary complex competition experiments, GST-FKBP12^F36V^ and 6xHis-p300-BRD (active motif #81158) were diluted to 250 nM and 500 nM, in assay buffer (50 mM HEPES pH 7.4, 200 mM NaCl, 1 mM TCEP, and 0.1% BSA). AceTAG-1 was added into protein mixture from 600 μM DMSO stock (final concentration 300 nM). 20 μL of protein mixture was added to each well of a 1/2 area 96-well OptiPlate (PerkinElmer). FKBP12^F36V^ competitor or p300-BRD competitor was then added from DMSO stock to protein mixture (0.4 μl/well) with 1% DMSO in final mixture. Plates were then shaken orbitally at room temperature for 1h. Nickel Chelate AlphaLISA Acceptor and Glutathione AlphaLISA Donor beads (PerkinElmer) diluted to 20 ng/µl in assay buffer and 20 μl beads mixture was added to each well. After a 1h orbital shaking at room temperature. lluminescence was measured on the Envision 2104 plate reader (PerkinElmer). Data were plotted using GraphPad PRISM v8 and fit with ‘log(inhibitor)-response, four parameters’ curve.

### AceTAG-1 Target Engagement

At 50%-70% confluency, HEK293t cells in 6-well plates were transfected with pLEX305-C-dTAG-H3.3. After 24h, cells were co-treated for 30 min with 0.5 ĿM FKBP-p probe and increasing concentration of AceTAG-1 in serum-free media. Cells then underwent UV-photocrosslinking (365 nm, energy 200 mJ/cm^2^) for 20 min at 4Ȣ, harvested in cold DPBS by scraping and centrifugation, cell pellets were washed with cold DPBS two times and aspirated. Cells were then lysed in 400 μL DPBS buffer supplemented with 0.2% SDS (w/v) and 1X Halt protease inhibitor cocktail. Lysate was then normalized to 1.5 mg/mL. To each sample (50 μL), 6 μL of a freshly prepared “click” reagent mixture containing 0.1 mM tris(benzyltriazolylmethyl)amine (TBTA) (3 μL/sample, 1.7 mM in 1:4 DMSO:*t-*ButOH), 1 mM CuSO_4_ (1 μL/sample, 50 mM in H_2_O), 25 μM Rhodamine-azide (1 μL/sample, 1.25 mM in DMSO), and freshly prepared 1 mM tris(2-carboxyethyl)phosphine HCl (TCEP) (1 μL/sample, 50 mM in H_2_O) was added to conjugate the fluorophore to probe-labeled proteins. Upon addition of the click mixture, each reaction was immediately mixed by vortexing and then allowed to react at room temperature for 1 h before quenching the reactions with SDS loading buffer (4X stock, 17 μL). Proteins (25 μg total protein loaded per gel lane) were resolved using SDS-PAGE (10% acrylamide) and visualized by in-gel fluorescence on Bio-Rad ChemiDoc MP Imaging System. Gel fluorescence and imaging was processed using Image Lab (v 6.0.1) software. Proteins were then transferred to PVDF membranes using Trans-Blot Turbo RTA Mini 0.45 mM LF PVDF Transfer Kit (Bio-Rad), and blotted with anti-HA antibody. Chemiluminescence was recorded with Bio-Rad ChemiDoc MP Imaging System and processed using Image Lab (v 6.0.1) software.

#### In-situ photocrosslinking with p300-BRD photocrosslinking probe and competition by AceTAG-1

At 50%-70% confluency, HEK293t cells in 6-well plates were transfected with pSG5-HA-p300. After 24 h, cells were co-treated for 30 min with 2.5 μM p300-p probe and increasing concentration of AceTAG-1 in serum-free media. Cells then underwent UV-photocrosslinking (365 nm, energy 200 mJ/cm^2^) for 20 min at 4Ȣ and harvested in cold DPBS by scraping and centrifugation. Cell pellets were washed with cold DPBS two times and aspirated. Cells were then lysed in 200 μL DPBS buffer supplemented with 1X Halt protease inhibitor cocktail. Lysate was fractionated by ultracentrifuge (100000 g, 45 min, 4Ȣ) to provide soluble fraction (supernatant) and membrane fraction (pellets). Soluble fraction was normalized to 2 mg/mL, underwent click reaction with Rhodamine-azide as previously described. Proteins (25 μg total protein loaded per gel lane) were resolved using SDS-PAGE (6% acrylamide). In-gel fluorescence visualization and immunoblot analysis was carried out as previously described.

### Immunoblotting

Cells were harvested in cold DPBS by scraping and washed with cold DPBS twice. Cell pellets were resuspended in DPBS supplemented with 5 mM sodium butyrate, 20 mM Nicotinamide, and 1X Halt Protease Inhibitor Cocktail and lysed by sonication (15 ms on, 40 ms off, 15% amplitude, 1s total on x 3). Protein concentration was determined using DC Protein Assay (Bio-Rad) and absorbance read using a Tecan, Infinite F500 plate reader following manufacturer’s instructions. Samples with equal protein content were boiled in 4X SDS gel loading buffer for 10 min. Proteins were separated by 10%, 12.5%, or 15% SDS-polyacrylamide gel electrophoresis and transferred to PVDF membranes using Trans-Blot Turbo RTA Mini 0.45μM LF PVDF Transfer Kit (Bio-Rad). Membranes were incubated for 1h at room temperature with blocking buffer, followed by incubating overnight at 4°C with primary antibodies. After washing in TBST, the secondary antibodies were incubated with the membranes at room temperature for 1h. The membranes were washed with TBST for three times and visualized on a Bio-Rad ChemiDoc MP Imaging System with Clarity Max Western ECL substrate (Bio-Rad) or SuperSignal West Femto Chemiluminescent substrate proprietary luminol and peroxide solution kit (Thermo Scientific Pierce).

### Immunoprecipitation

Cells were split in 10 cm dishes in indicated media. The probes were added once the cells grew to 70-80% confluency and incubated for indicated time. The cells were harvested and washed with cold DPBS twice. The pelleted cells were lysed in 1 mL lysis buffer, containing 50 mM Tris-HCl pH 7.5, 300 mM NaCl, 0.5% (v/v) NP40, 5 mM sodium butyrate, 20 mM Nicotinamide, and 1X Halt Protease Inhibitor, followed by clarification by centrifugation at 16,000 g for 20 min at 4°C. The protein concentration was normalized by DC Proein Assay (Bio-Rad) and equal amount of protein was subjected to enrichment. For HA tag fusion protein enrichment, Monoclonal Anti-HA Agarose antibody produced in mouse (Sigma-Aldrich) was pre-washed with lysis buffer and added to the clarified lysate. Enrichment was carried out at 4°C for 4h with rotating. After allowing immune complex binding, the beads were spun down at 2,000 rpm for 3 min at 4°C. The beads were washed with lysis buffer and DPBS and incubated with HA peptide elution buffer at 37°C for 15 min twice or 4X SDS gel loading buffer at 95°C for 10 min. Supernatant was collected for immunoblot analysis or mass spectrometry sample preparation.

### Chromatin Isolation

To isolate chromatin, 3X10^6^ cells were resuspended in 200 μL buffer A (10 mM HEPES, pH 7.9, 10 mM KCl, 1.5 mM MgCl_2_, 0.34 M sucrose, 10% glycerol, 1 mM DTT, 5 mM sodium butyrate, 20 mM Nicotinamide, and 1X Halt Protease Inhibitor). Cells were incubated for 5 min on ice with 0.1% (v/v) Triton X-100. Nuclei and soluble protein were separated by low-speed centrifugation at 1,300 g for 15 min at 4°C. The soluble protein was further clarified by high-speed centrifugation at 20,000 g for 15 min at 4°C. Nuclei was washed with buffer A, following by lysis in buffer B (3 mM EDTA, 0.2 mM EGTA, 1 mM DTT, 5 mM sodium butyrate, 20 mM Nicotinamide, and 1X Halt Protease Inhibitor). Insoluble chromatin and nuclear plasma were separated by centrifugation at 1,700 g for 4 min at 4°C. The insoluble chromatin was washed once in buffer B and resuspended in DPBS with 0.1% SDS. Protein concentration of chromatin, nuclear plasma, and soluble protein were normalized by DC Protein Assay (Bio-Rad) and equal amount of protein was subjected to immunoblotting analysis as described above.

### Quantitative Proteomic Analysis of H3.3 Acetylation

#### Sample preparation

Cells were split in 10 cm dishes in indicated media. The probes were added once the cells grew to 60%-70% confluency and incubated for indicated time. The cells were harvested and washed with cold DPBS twice. The pelleted cells were lysed in 1 mL lysis buffer, containing 10 mM Tris pH 7.5, 30 mM NaCl, 0.1% (v/v) NP40, 5 mM sodium butyrate, 20 mM Nicotinamide, and 1X Halt Protease Inhibitor, followed by centrifuging for 5 min at 1,000 g at 4°C. The pellets were resuspended in benzonase buffer, containing 50 mM Tris pH 7.5, 300 mM NaCl, 0.5% NP40, and 2.5 mM MgCl_2_. Protein concentration was normalized by DC Protein Assay (Bio-Rad) and equal amount of protein was subjected to benzonase nuclease (Millipore Sigma) treatment. 50 U/100 μl of benznase was added to each sample. Samples with benzonase were incubated on ice for 30 min and centrifuged at 300 g for 3 min at 4 °C to obtain soluble chromatin fraction. Samples were diluted with dilution buffer, containing 50mM Tris pH 7.5, 300 mM NaCl, 0.5% NP40, and 15 mM EDTA to quench the benzonase activity. Protein concentration was normalized again by DC Protein Assay (Bio-Rad) and equal amount of protein was used for HA-tag protein enrichment as described above. The beads were washed with lysis buffer and DPBS and incubated with 8 M urea and boiled in 4X SDS gel loading buffer at 95°C for 15 min. Supernatant was collected for in-gel digestion for mass-spectrometry sample preparation. HA-tag protein enriched samples were run on SDS-PAGE gel, flanked by MW markers, and stained with ProtoBlue. The desire protein bands were excised out and dice into approximately 1 mm^3^ pieces. The gels were washed with 500 μl 100 mM TEAB for 2 times and reduced in 10 mM TCEP solution at 60°C for 30 min. After reaction, TCEP solution was replaced with a solution of freshly prepared iodoacetamide (55 mM in 100 mM TEAB) and incubated for 30 min at room temperature while protected from light. A solution of 1:1 acetonitrile and 100 mM TEAB was added to wash the gels bands, followed by 100% acetonitrile to completely dry the gels. Sequencing-grade modified porcine trypsin (0.2 μg, 25 mM TEAB pH 8.5, 100 µM CaCl_2_) was added to the gel and incubated at 37 °C for 14 h with shaking. The digest was collected and the peptides within the gels were extracted with 25% acetonitrile/5% formic acid (MS-grade) for one time, 75% acetonitrile (MS-grade) for two times, and 100% acetonitrile (MS-grade) for two times. The peptide samples were dried under vacuum centrifugation.

#### TMT Labeling and Fractionation

Samples were dissolved in 100 mM TEAB containing 30 % acetonitrile (MS-grade) and labeled with respective TMT 10 plex isotope (8 μL, 20 µg/ µL) for 1 h with occasional vortexing at RT. To quench the reaction, hydroxylamine (6 μL, 5% *v/v*) was added to each sample, vortexed, and incubated for 15 min at RT. Formic acid (4 μL) was added to each tube to acidify and the samples were dried under vacuum centrifugation. Multiplexed samples were fractionated using the Pierce™ High pH Reversed-Phase Peptide Fractionation Kit according to manufacturer instructions with the following elution scheme (% acetonitrile in 0.1% TEA; F1: 5%, F2: 7.5%, F3: 10%, F4: 12.5%, F5: 15%, F6: 17.5%, F7: 20%, F8: 22.5%, F9: 25%, F10: 30%, F11: 50%, F12: 95%). Fractions were then combined into six fractions by the following scheme: F1+F7, F2+F8, F3+F9, F4+F10, F5+F11, F6+F12. Combined fractions were lyophilized prior to resuspension for MS analysis.

#### MS Analysis

TMT labeled samples were redissolved in MS buffer A (20 μL, 0.1% formic acid in water). 3 μL of each sample was loaded onto an Acclaim PepMap 100 precolumn (75 µm x 2 mm) and eluted on an Acclaim PepMap RSLC analytical column (75 µm x 15 cm) using the UltiMate 3000 RSLCnano system (Thermo Fisher Scientific). Buffer A (0.1% formic acid) and buffer B (0.1% formic acid in MeCN) were used in a 200 min gradient (flow rate 0.3 mL/min, 35 °C) of 2 % buffer B for 10 min, followed by an incremental increase to 25 % buffer B over 155 min, 25%-45% buffer B for 10 min, 45%-95 % buffer B for 5 min, hold at 95 % buffer B for 2 min, followed by descent to 2% buffer B for 1 min and re-equilibration at 2% for 6 min. The elutions were analyzed with a Thermo Fisher Scientific Orbitrap Fusion Lumos mass spectrometer with a cycle time of 3 s and nano-LC electrospray ionization source applied voltage of 2.0 kV. MS^1^ spectra were recorded at a resolution of 120,000 with an automatic gain control (AGC) value of 1×10^6^ ions, maximum injection time of 50 ms (dynamic exclusion enabled, repeat count 1, duration 20 s). The scan range was specified from 375 to 1,500 m/z. Peptide fragmentation MS^2^ spectra was recorded via collision-induced diffusion (CID) and quadrupole ion trap analysis (AGC 1.8×10^4^, 30 % collision energy, maximum inject time 120 ms, isolation window 0.7). MS^3^ spectra were generated by high-energy collision-induced dissociation (HCD) with collision energy of 65 %. Precursor selection included up to 10 MS^2^ ions for the MS^3^ spectrum.

Proteomic analysis was performed with the processing software Proteome Discoverer 2.4 (Thermo Fisher Scientific). Peptide sequences were identified by matching proteome databases with experimental fragmentation patterns via the SEQUEST HT algorithm. Fragment tolerances were set to 0.6 Da, and precursor mass tolerances set to 10 ppm with four missed cleavage sites allowed. Carbamidomethyl (C, +57.02146) and TMT-tag (N-terminal, +229.163) were specified in the static modifications. Oxidation (M, +15.995), TMT-tag (K, +229.163), and Acetylation (K, +42.001) were defined as dynamic modifications. Spectra were searched against the *Homo Sapiens* proteome database (42,358 sequences) using a false discovery rate of 1 % (Percolator). MS^3^ peptide quantitation was performed with a mass tolerance of 20 ppm. Peptide abundance of TMT ratios obtained by Proteome Discoverer were normalized to total protein abundance.

### Quantitative Acetylproteomics

#### Sample preparation

Frozen cell pellets were thawed on ice and lysed in a buffer containing 50 mM Tris-HCl (pH 7.5), 150 mM NaCl, 1 mM EDTA, 1% NP-40, 0.1% sodium deoxycholate, 5 mM sodium butyrate, 10 mM nicotinamide and 1X Pierce™ protease inhibitor cocktail. Cells were sonicated to ensure complete lysis, mixed with 5 M NaCl at a ratio of 1:10 with sample volume and incubated on ice for 15 min to release chromatin-bound proteins. Cell lysates were then sonicated again to shear genomic DNA. Cellular debris was cleared by centrifugation at 16,000 g for 20 min at 4°C, protein concentration was determined using a Pierce™ BCA assay and ∼10 mg of protein was aliquoted for acetyl-proteomic analysis. Proteins were denatured with 8 M urea in 50 mM HEPES, reduced with 5 mM dithiothreitol for 30 min at 56°C, then alkylated with 5 mM iodoacetamide for 20 min in the dark at RT. Following alkylation, proteins were precipitated using chloroform-methanol as previously described, and protein pellets were dried at 56°C for 15 min. Proteins pellets were resuspended in 1 M urea in 50 mM HEPES and digested in a two-step process with LysC and trypsin as previously described. Digested peptides were desalted using Phenomenex Strata-X polymeric reverse phase extraction cartridges using the following protocol. Cartridges were conditioned with 1 mL 100% acetonitrile, 1 mL 80% acetonitrile/0.5% AA, and 1 mL 0.1% TFA. Peptides were then loaded on to the cartridge at a reduced flow rate and desalted with 5 mL 0.1% TFA followed by 1 mL 0.5% AA. Desalted peptides were eluted using 1 mL 40% acetonitrile/0.5% AA followed by 1 mL 80% acetonitrile/0.5% AA and lyophilized. Digested peptides were quantified using the Pierce™ Quantitative Colorimetric Peptide Assay and 25 μg was aliquoted from each for total protein analysis while the remaining peptide was lyophilized and retained for acetyl-peptide enrichment.

#### Acetyl-peptide Enrichment

Acetyl-peptides were enriched using the PTMScan® Acetyl-Lysine Motif [Ac-K] Kit according to manufacturer instructions. Briefly, lyophilized peptides were resuspended in immuno-affinity purification buffer (IAP) and added to PBS-washed antibody-bead slurry. Acetyl-peptides were enriched on a rotator for 2h at 4°C. Bead-peptide complexes were then washed 2X with 1 mL IAP and 3X with 1 mL H2O. Acetyl-peptides were eluted from the beads with 2X treatments of 0.15% TFA (gently mixing at RT for 10 min for each elution), desalted with Phenomenex Strata-X polymeric reverse phase extraction cartridge as above and lyophilized prior to TMT labeling.

#### TMT Labeling and Fractionation

Peptide aliquots were resuspended in 30% dry acetonitrile, 200 mM HEPES, pH 8.5 and 8 μl of TMT labeling reagents was added to each sample. Labeling was allowed to proceed for 1h at RT after which TMT labels were quenched by the addition of 9 ul 5% hydroxylamine for 15 min before the samples were acidified with 50 ul 1% TFA. Samples were then pooled into respective 10plexes, lyophilized and resuspended in 0.1% TFA. Multiplexed samples were fractionated using the Pierce™ High pH Reversed-Phase Peptide Fractionation Kit according to manufacturer instructions with the following elution scheme (% acetonitrile in 0.1% TEA; F1: 7.5%, F2: 10%, F3: 12.5%, F4: 15%, F5: 17.5%, F6: 20%, F7: 22.5%, F8: 25%, F9: 27.5%, F10: 30%, F11: 32.5%, F12: 35%, F13: 37.5%, F14: 40%, F15: 42.5%, F16: 45%, F17: 47.5%, F18: 75%). Fractions were then combined into nine fractions by the following scheme: F1+F10, F2+F11, F3+F12, F4+F13, F5+F14, F6+F15, F7+F16, F8+F17, F9+18. Combined fractions were lyophilized prior to resuspension for MS analysis.

#### MS analysis

Acetylproteomic samples were subject to analysis via LC-MS^3^ as above with the following alterations. Samples were resuspended in 60 μL buffer A and 10 μL was loaded on to the analytical column. Samples were eluted with 5% buffer B for 10 min, a gradient of 5-20% buffer B over 160 min, 20-45% buffer B over 20 min, 45-95% buffer B over 5 min, 95% buffer B for 2 min, 5% buffer B for 2 min, 95% buffer B for 2 min and 5% buffer B for 10 min to re-equilibrate the column. Eluted peptides were analyzed with a Thermo Fisher Orbitrap Fusion mass spectrometer as above with the following alterations: MS^1^ scan range of 400-1700 m/z, dynamic exclusion of 15 sec, MS^1^ AGC target of 2×10^5^, MS^2^ CID collision energy 35%, MS^3^ HCD collision energy of 55%.

### Data availability

Mass spectrometry datasets are deposited on the ProteomeXchange database and will be made publicly available upon manuscript acceptance. All other raw data are available upon requests.

### Chemistry Materials

Chemicals and reagents were purchased from commercial vendors, including Sigma-Aldrich, Fisher Scientific, Combi-Blocks, MedChemExpress, Alfa Aesar and AstaTech, and were used as received without further purification, unless otherwise noted. Anhydrous solvents were purchased from Sigma-Aldrich in Sure/Seal™ formulations. All reactions were monitored by thin-layer chromatography (TLC, Merck silica gel 60 F-254 plates). The plates were stained either with p-anisaldehyde (2.5% p-anisaldehyde, 1% AcOH, 3.5% H2SO4 (conc.) in 95% EtOH), ninhydrin (0.3% ninhydrin (w/v), 97:3 EtOH-AcOH), KMnO4 (1.5g of KMnO4, 10g K2CO3, and 1.25mL 10% NaOH in 200mL water), iodine or directly visualized with UV light. Reaction purification was carried out using Flash chromatography (230 – 400 mesh silica gel), Biotage® or preparative thin layer chromatography (pTLC, Analtech, 500-2000 μm thickness). NMR spectra were recorded on Bruker DPX-400 MHz or Bruker AV-600 MHz spectrometers in the indicated solvent. Multiplicities are reported with the following abbreviations: s singlet; d doublet; t triplet; q quartet; p pentet; m multiplet; br broad; dd doublet of doublets; dt doublet of triplets; td triplet of doublets; Chemical shifts are reported in ppm relative to the residual solvent peak and J values are reported in Hz. Mass spectrometry data were collected on an Agilent 6120 single-quadrupole LC/MS instrument (ESI, low resolution) or an Agilent ESI-TOF instrument (ESI-TOF, HRMS).

## COMPOUND SYNTHESIS AND CHARACTERIZATION

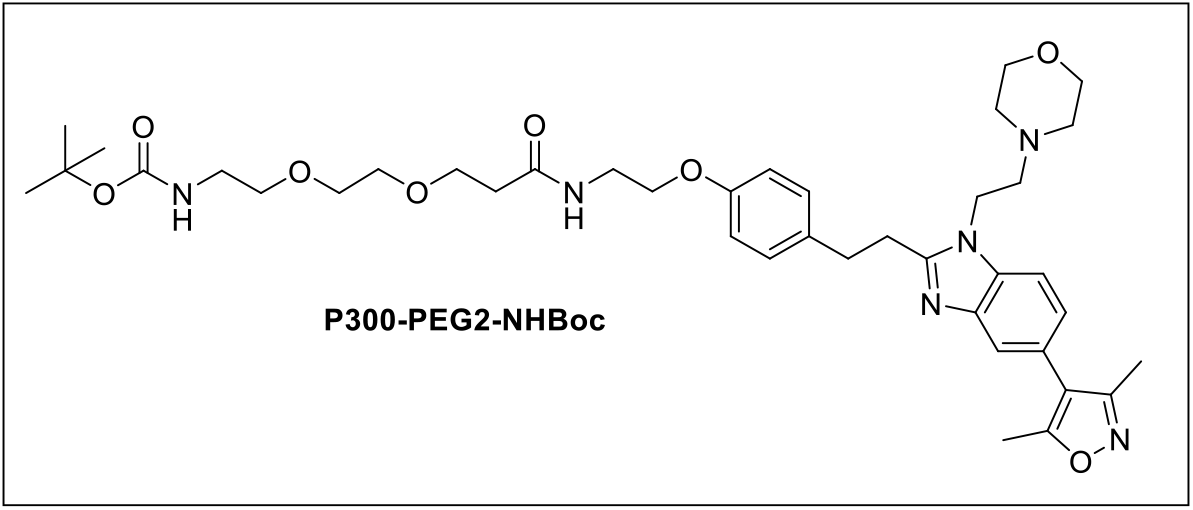

### tert-butyl (2-(2-(3-((2-(4-(2-(5-(3,5-dimethylisoxazol-4-yl)-1-(2-morpholinoethyl)-1H-benzo[d]imidazol-2-yl)ethyl)phenoxy)ethyl)amino)-3-oxopropoxy)ethoxy)ethyl)carbamate (P300-PEG2-NHBoc)

To a solution of 2-(4-(2-(5-(3,5-dimethylisoxazol-4-yl)-1-(2-morpholinoethyl)-1H-benzo[d]imidazol-2-yl)ethyl)phenoxy)ethan-1-amine (10.0 mg, 0.020 mmol, 1.0 eq) (synthesized according to previous method^1^) in DMF (1 mL), DIPEA (10.6 µL, 0.061 mmol, 3.0 eq) and t-Boc-N-amido-PEG2-acid (5.35 mg, 0.020 mmol, 1 eq) was added followed by HATU (8.54 mg, 0.022 mmol, 1.1 eq) were added at 0°C and resulting mixture was stirred for 5 minutes the corresponding starting material was fully consumed (indicated by TLC). The crude mixture was diluted with cold water and extracted in ethyl acetate (20 mL X 3) then combined organic extract was dried over sodium sulfate, filtered, and concentrated under reduced pressure. The crude material was purified by PTLC (DCM/Methanol, 19:1) to obtain P300-PEG2-NHBoc as a colorless sticky material (8.4 mg, 55 %).

**^1^H NMR:** (400 MHz, CDCl_3_) δ 7.62 (dd, J = 1.5, 0.6 Hz, 1H), 7.35 (dd, J = 8.3, 0.7 Hz, 1H), 7.18 – 7.10 (m, 3H), 6.88 – 6.80 (m, 3H), 4.13 (t, J = 6.9 Hz, 2H), 4.02 (t, J = 5.3 Hz, 2H), 3.75 – 3.63 (m, 10H), 3.62 – 3.56 (m, 4H), 3.27 – 3.20 (m, 2H), 3.19 – 3.13 (m, 2H), 2.62 (t, J = 6.8 Hz, 2H), 2.52 – 2.44 (m, 8H), 2.42 (s, 3H), 2.29 (s, 3H), 1.43 (s, 9H).

**^13^C NMR:** (101 MHz, CDCl_3_) δ 171.87, 170.88, 165.04, 159.02, 157.32, 155.38, 143.00, 134.27, 133.34, 129.41, 124.27, 123.46, 119.92, 117.10, 114.71, 109.44, 80.67, 70.24, 70.11, 67.19, 66.89, 66.84, 57.64, 54.06, 41.52, 38.88, 36.94, 36.22, 32.98, 29.88, 28.11, 11.60, 10.90.

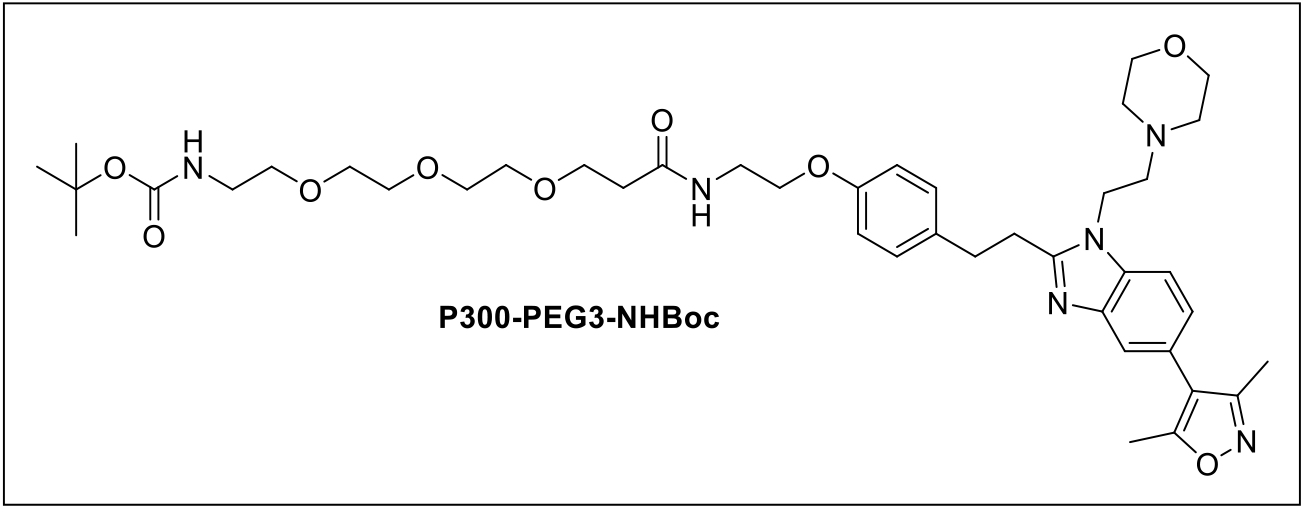

### tert-butyl (15-(4-(2-(5-(3,5-dimethylisoxazol-4-yl)-1-(2-morpholinoethyl)-1H-benzo[d]imidazol-2-yl)ethyl)phenoxy)-12-oxo-3,6,9-trioxa-13-azapentadecyl)carbamate (P300-PEG3-NHBoc)

To a solution of 2-(4-(2-(5-(3,5-dimethylisoxazol-4-yl)-1-(2-morpholinoethyl)-1H-benzo[d]imidazol-2-yl)ethyl)phenoxy)ethan-1-amine (10.0 mg, 0.020 mmol, 1.0 eq) (synthesized according to previous method^1^) in DMF (1 mL), DIPEA (10.6 µL, 0.061 mmol, 3.0 eq) and t-Boc-N-amido-PEG3-acid (6.56 mg, 0.020 mmol, 1 eq) was added followed by HATU (8.54 mg, 0.022 mmol, 1.1 eq) were added at 0°C and resulting mixture was stirred for 5 minutes the corresponding starting material was fully consumed (indicated by TLC). The reaction mixture was diluted with cold water and extracted in ethyl acetate (20 mL X 3) then combined organic extract was dried over sodium sulfate, filtered, and concentrated under reduced pressure. The crude material was purified by PTLC (DCM/Methanol, 19:1) to obtain P300-PEG2-NHBoc as a colorless sticky material (8.5 mg, 52 %).

**^1^H NMR:** (600 MHz, CDCl_3_) δ 7.62 (d, J = 1.5 Hz, 1H), 7.35 (d, J = 8.3 Hz, 1H), 7.16 – 7.13 (m, 2H), 7.12 (dd, J = 8.2, 1.5 Hz, 1H), 6.84 (d, J = 8.6 Hz, 2H), 6.76 (s, 1H), 5.07 (s, 1H), 4.13 (t, J = 6.9 Hz, 2H), 4.02 (t, J = 5.3 Hz, 2H), 3.74 (t, J = 5.8 Hz, 2H), 3.68 – 3.61 (m, 10H), 3.60 (s, 4H), 3.52 (t, J = 5.1 Hz, 2H), 3.33 – 3.27 (m, 2H), 3.22 – 3.13 (ddd, J = 9.7, 5.1, 2.1 Hz, 2H), 3.17 (ddd, J = 8.5, 7.0, 2.2 Hz, 2H), 2.62 (t, J = 6.9 Hz, 2H), 2.51 (t, J = 5.8 Hz, 2H), 2.46 (t, J = 4.6 Hz, 4H), 2.42 (s, 3H), 2.29 (s, 3H), 1.42 (s, 9H).

**^13^C NMR:** (151 MHz, CDCl_3_) δ 171.92, 165.06, 159.03, 157.26, 155.39, 142.97, 134.27, 133.36, 129.42, 124.25, 123.47, 119.89, 117.11, 114.71, 109.47, 70.56, 70.35, 70.20, 70.16, 70.04, 67.24, 66.84, 66.76, 57.63, 54.05, 41.51, 40.34, 38.92, 32.97, 29.86, 28.42, 11.61, 10.92.

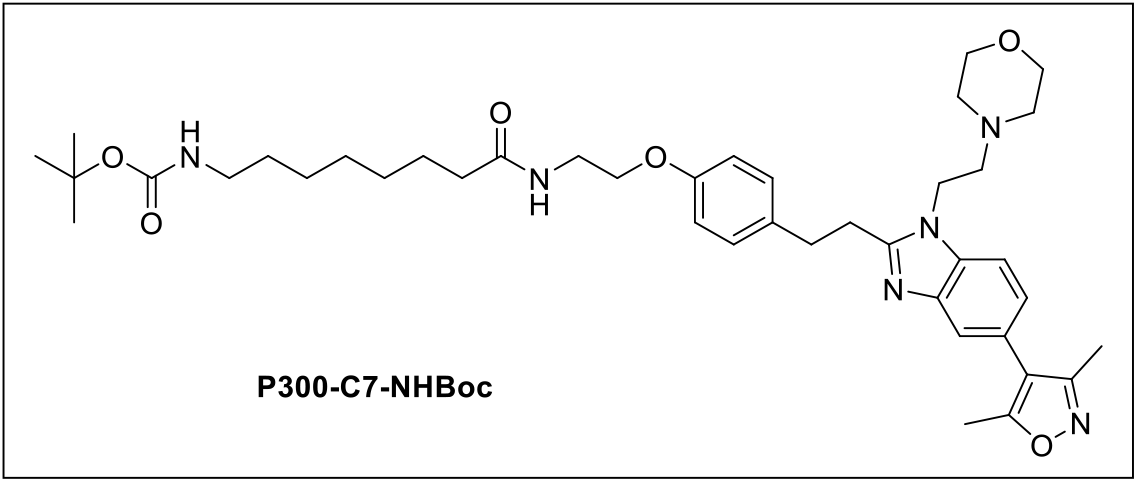

### tert-butyl (8-((2-(4-(2-(5-(3,5-dimethylisoxazol-4-yl)-1-(2-morpholinoethyl)-1H-benzo[d]imidazol-2-yl)ethyl)phenoxy)ethyl)amino)-8-oxooctyl)carbamate (P300-C7-NHBoc)

To a solution of 2-(4-(2-(5-(3,5-dimethylisoxazol-4-yl)-1-(2-morpholinoethyl)-1H-benzo[d]imidazol-2-yl)ethyl)phenoxy)ethan-1-amine (30.0 mg, 0.061 mmol, 1.0 eq) (synthesized according to previous method^1^) and Boc-8-Aoc-OH (17.5 mg, 0.067 mmol, 1.1 eq) in DCM (4 mL), EDC (18.0 mg, 0.092 mmol, 1.5 eq) HOBt (13.0 mg, 0.092mmol, 1.5 eq) and DIPEA (32 µL, 0.183 mmol, 3.0 eq) was added. The mixture was stirred at room temperature for 14 hours. After completion the reaction mixture was diluted with water (10 mL) and extracted in DCM (2 X 20 mL), the organic layer was washed with saturated sodium bicarbonate (aq), and brine, the combined organic layer was dried over sodium sulfate, filtered, and concentrated under reduced pressure. The crude material was purified by Biotage® Sfär Silica D 10 g column with a 0-5 % linear gradient of Methanol in Dichloromethane over 20 column volumes (CV), to obtain P300-C7-NHBoc as a colorless oil (29 mg, 65 %).

**^1^H NMR** (600 MHz, CDCl_3_) δ 7.62 (d, J = 1.5 Hz, 1H), 7.36 (s, 1H), 7.18 – 7.15 (m, 2H), 7.12 (dd, J = 8.2, 1.5 Hz, 1H), 6.86 – 6.82 (m, 2H), 5.93 (t, J = 5.9 Hz, 1H), 4.51 (s, 1H), 4.13 (t, J = 6.9 Hz, 2H), 4.01 (t, J = 5.1 Hz, 2H), 3.66 (td, J = 4.9, 2.8 Hz, 6H), 3.27 – 3.21 (m, 2H), 3.20 – 3.14 (m, 2H), 3.08 (q, J = 6.7 Hz, 2H), 2.62 (t, J = 6.9 Hz, 2H), 2.46 (t, J = 4.6 Hz, 4H), 2.42 (s, 3H), 2.29 (s, 3H), 2.21 – 2.16 (m, 2H), 1.63 (t, J = 7.3 Hz, 3H), 1.43 (s, 9H), 1.33 – 1.28 (m, 6H).

**^13^C NMR** (151 MHz, CDCl_3_) δ 173.26, 165.06, 159.03, 157.18, 156.01, 155.35, 143.03, 134.28, 133.51, 129.47, 124.26, 123.46, 119.95, 117.10, 114.66, 109.42, 66.94, 66.84, 57.64, 54.06, 41.50, 40.52, 38.89, 36.65, 32.94, 29.98, 29.87, 29.72, 29.11, 28.91, 28.44, 26.56, 25.52, 11.61, 10.92.

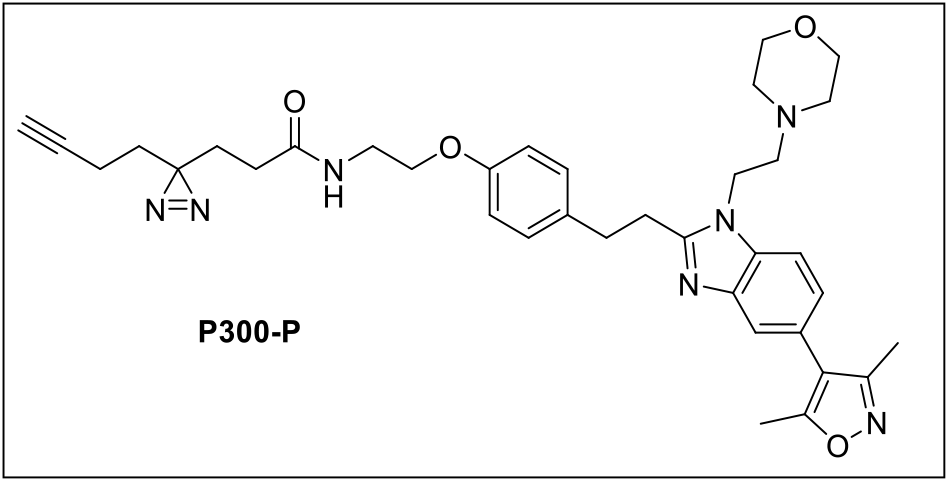

### 3-(3-(but-3-yn-1-yl)-3H-diazirin-3-yl)-N-(2-(4-(2-(5-(3,5-dimethylisoxazol-4-yl)-1-(2-morpholinoethyl)-1H-benzo[d]imidazol-2-yl)ethyl)phenoxy)ethyl)propenamide (P300-P)

To a solution of 2-(4-(2-(5-(3,5-dimethylisoxazol-4-yl)-1-(2-morpholinoethyl)-1H-benzo[d]imidazol-2-yl)ethyl)phenoxy)ethan-1-amine (15.0 mg, 0.030 mmol, 1.0 eq) (synthesized according to previous method^1^) and 3-(3-(but-3-yn-1-yl)-3H-diazirin-3-yl)propanoic acid (6.0 mg, 0.033 mmol, 1.1 eq) in DCM (2 mL), EDC (9.0 mg, 0.046 mmol, 1.5 eq) HOBt (7.0 mg, 0.046 mmol, 1.5 eq) and DIPEA (16 µL, 0.092 mmol, 3.0 eq) was added. The mixture was stirred at room temperature for 12 hours, then diluted with DCM. The organic layer was washed with saturated sodium bicarbonate (aq), water and brine. The combined organic layer was then dried over sodium sulfate, filtered, and concentrated under reduced pressure. The crude material was purified by Biotage® Sfär Silica D 10 g with a 0-5 % linear gradient of Methanol in Dichloromethane over 20 column volumes (CV), to obtain P300-diazirine tag as a colorless oil (12.7 mg, 65 %).

**^1^H NMR** (400 MHz, CDCl_3_) δ 7.62 (d, J = 1.6 Hz, 1H), 7.35 (d, J = 8.3 Hz, 1H), 7.19 – 7.09 (m, 3H), 6.86 – 6.79 (m, 2H), 6.10 – 5.91 (m, 1H), 4.13 (t, J = 6.8 Hz, 2H), 4.01 (t, J = 5.1 Hz, 2H), 3.70 – 3.61 (m, 6H), 3.23 (dd, J = 8.9, 5.8 Hz, 2H), 3.20 – 3.12 (m, 2H), 2.61 (t, J = 6.8 Hz, 2H), 2.45 (t, J = 4.6 Hz, 4H), 2.42 (s, 3H), 2.29 (s, 3H), 2.03 – 1.93 (m, 5H), 1.84 (dd, J = 8.9, 6.6 Hz, 2H), 1.63 (t, J = 7.3 Hz, 2H).

**^13^C NMR** (101 MHz, CDCl_3_) δ 171.30, 165.04, 159.02, 157.09, 155.34, 143.03, 134.28, 133.56, 129.47, 124.22, 123.45, 119.92, 117.10, 114.63, 109.43, 82.71, 69.26, 66.83, 66.73, 57.64, 54.05, 41.50, 39.04, 32.92, 32.35, 30.31, 29.85, 28.35, 27.85, 13.28, 11.60, 10.92.

**HRMS** (ESI-TOF) calcd for C_36_H_44_N_7_O_4_, 638.3449 (M+H^+^), found 638.3451.

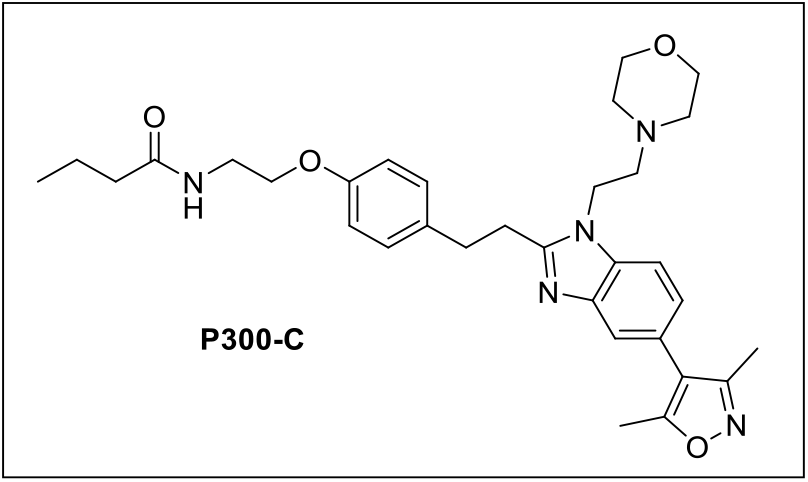

### N-(2-(4-(2-(5-(3,5-dimethylisoxazol-4-yl)-1-(2-morpholinoethyl)-1H-benzo[d]imidazol-2-yl)ethyl)phenoxy)ethyl)pentanamide (P300-C)

To a solution of 2-(4-(2-(5-(3,5-dimethylisoxazol-4-yl)-1-(2-morpholinoethyl)-1H-benzo[d]imidazol-2-yl)ethyl)phenoxy)ethan-1-amine (synthesized according to previous method^1^) (15.0 mg, 0.030 mmol, 1.0 eq) and butyric acid (3.0 mg, 0.034 mmol, 1.1 eq) in DCM (2 mL), EDC (9.0 mg, 0.046 mmol, 1.5 eq) HOBt (7.0 mg, 0.046 mmol, 1.5 eq) and DIPEA (16 µL, 0.092 mmol, 3.0 eq) was added. The mixture was stirred at room temperature for 12 hours, then diluted with DCM. The organic layer was washed with saturated sodium bicarbonate (aq), water and brine. The combined organic layer was then dried over sodium sulfate, filtered, and concentrated under reduced pressure. The crude material was purified by PTLC (DCM/Methanol, 19:1) to afford P300-C as a colorless oil (10.0 mg, 58%).

**^1^H NMR:** (400 MHz, MeOD) δ 7.61 – 7.54 (m, 2H), 7.24 (dd, J = 8.3, 1.6 Hz, 1H), 7.17 – 7.09 (m, 2H), 6.86 (d, J = 8.6 Hz, 2H), 4.20 (t, J = 6.7 Hz, 2H), 4.01 (t, J = 5.5 Hz, 2H), 3.64 (t, J = 4.7 Hz, 4H), 3.55 (t, J = 5.5 Hz, 2H), 3.27– 3.24 (m, 2H), 3.23 – 3.14 (m, 2H), 2.54 (t, J = 6.7 Hz, 2H), 2.49 – 2.41 (m, 7H), 2.29 (s, 3H), 2.20 (dd, J = 7.9, 6.9 Hz, 2H), 1.69 – 1.60 (m, 2H), 0.94 (t, J = 7.4 Hz, 3H).

**^13^C NMR:** (151 MHz, MeOD) δ 175.06, 165.34, 158.83, 157.56, 156.11, 142.02, 134.11, 132.96, 129.19, 124.07, 123.51, 118.43, 116.96, 114.36, 110.25, 66.45, 66.15, 57.06, 53.63, 40.86, 38.67, 37.49, 32.95, 29.16, 18.96, 12.53, 10.00, 9.33.

**HRMS** (ESI-TOF) calculated for C_32_H_41_N_5_O_4,_ 560.3231 (M+H^+^), found 560.3247.

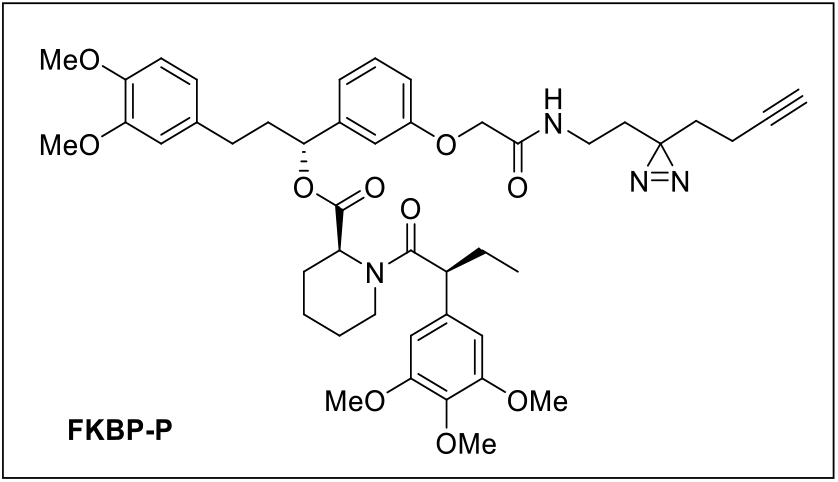

### 3) (R)-1-(3-(2-((2-(3-(but-3-yn-1-yl)-3H-diazirin-3-yl)ethyl)amino)-2-oxoethoxy)phenyl)-3-(3,4-dimethoxyphenyl)propyl (S)-1-((S)-2-(3,4,5-trimethoxyphenyl)butanoyl)piperidine-2-carboxylate (FKBP-P)

To a solution of 2-(3-((R)-3-(3,4-dimethoxyphenyl)-1-(((S)-1-((S)-2-(3,4,5-trimethoxyphenyl)butanoyl)piperidine-2-carbonyl)oxy)propyl)phenoxy)acetic acid (synthesized according to previous method^2^) (10.0 mg, 0.014 mmol, 1.0 eq) and 2-(3-(but-3-yn-1-yl)-3H-diazirin-3-yl)ethan-1-amine (2.20 mg, 0.016 mmol, 1.1 eq) in DCM (2 mL), EDC (4.15 mg, 0.022 mmol, 1.5 eq), HOBt (2.94 mg, 0.022 mmol, 1.5 eq) and DIPEA (8.0 µL, 0.043 mmol, 3.0 eq) was added. The mixture was stirred at room temperature for 12 hours, after completion the reaction mixture was diluted with DCM (15 mL). The organic layer was washed with saturated sodium bicarbonate (aq), water and brine, the combined organic layer was dried over sodium sulfate, filtered, and concentrated under reduced pressure. The crude material was purified by PTLC (DCM/Methanol, 19:1) to afford FKBP-P as a colorless oil (8.0 mg, 58%).

**^1^H NMR** (600 MHz, CDCl_3_) δ 7.33 (t, J = 7.9 Hz, 0.25H), 7.19 (t, J = 7.9 Hz, 1H), 7.01– 6.99 (m, 0.25H), .6.93– 6.92 (m, 0.25H), 6.88 – 6.86 (m, 0.46H), 6.82 – 6.80 (m, 1.60H), 6.79 – 6.76 (m, 0.25H), 6.69 – 6.66 (m, 1.37H), 6.66 – 6.62 (m, 1.20H), 6.42 (s, 0.5H), 6.40 (s, 2H), 5.81 (dd, J = 7.8, 6.0 Hz, 0.23H), 5.63 (dd, J = 8.3, 5.4 Hz, 1H), 5.47 – 5.44 (m, 1H), 4.66 (d, J = 5.4 Hz, 0.22H), 4.61 – 4.58 (m, 0.26H), 4.50 (s, 0.5H), 4.48 (s, 2H), 3.87 – 3.83 (m, 10H), 3.82 – 3.78 (m, 1H), 3.78 (s, 3H), 3.67 (s, 6H), 3.59 – 3.57 (m, 1H), 3.39 – 3.37 (m, 0.26H), 3.28 – 3.21 (m, 2.5H), 2.79 (td, J = 13.4, 3.1 Hz, 1H), 2.63 – 2.53 (m, 1.5H), 2.50 – 2.44 (m, 1H), 2.42 – 2.40 (m, 0.23H), 2.30 (dd, J = 12.0, 2.6 Hz, 1H), 2.13 – 2.04 (m, 2.61H), 2.02 – 1.98 (m, 4H), 1.96 – 1.91 (m, 1H), 2.04 1.77 – 1.67 (m, 6H), 1.66 – 1.61 (m, 3H), 1.47 – 1.39 (m, 1H), 1.31 – 1.25 (m, 3H), 0.90 (t, J = 7.3 Hz, 3H), 0.86 (t, J = 7.3 Hz, 1H). *Note: rotomeric isomers observed*.

**^13^C NMR** (101 MHz, CDCl_3_) δ 172.67, 170.65, 168.26, 157.31, 153.51, 153.20, 148.89, 147.37, 142.41, 136.64, 135.32, 133.34, 130.14, 129.85, 120.21, 120.11, 120.07, 119.92, 113.79, 113.72, 113.55, 113.08, 111.69, 111.59, 111.34, 111.27, 104.98, 104.58, 82.60, 75.60, 69.45, 67.37, 67.31, 60.92, 60.80, 56.33, 55.97, 55.94, 55.88, 55.86, 55.69, 52.06, 51.25, 50.81, 43.49, 39.69, 38.24, 38.01, 34.05, 34.02, 32.66, 32.63, 32.05, 31.46, 31.30, 29.72, 28.40, 28.37, 26.79, 26.74, 26.36, 25.34, 24.54, 20.91, 20.68, 13.22, 12.79, 12.58. *Note: rotomeric isomers observed*.

**HRMS** (ESI-TOF) calculated for C_45_H_57_N_4_O_10,_ 813.4069 (M+H^+^), found 813.4072.

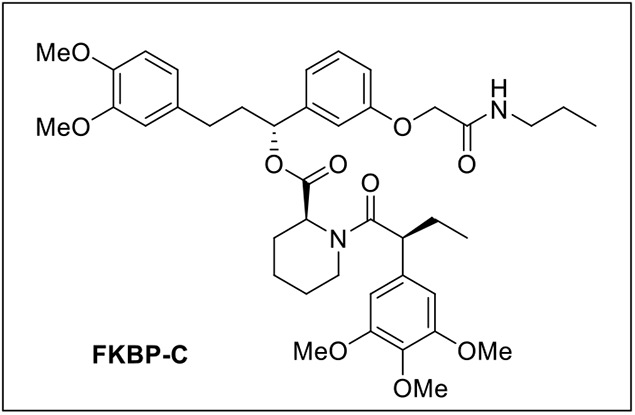

### (R)-3-(3,4-dimethoxyphenyl)-1-(3-(2-oxo-2-(propylamino)ethoxy)phenyl)propyl (S)-1-((S)-2-(3,4,5-trimethoxyphenyl)butanoyl)piperidine-2-carboxylate (FKBP-C)

To a solution of 2-(3-((R)-3-(3,4-dimethoxyphenyl)-1-(((S)-1-((S)-2-(3,4,5-trimethoxyphenyl)butanoyl)piperidine-2-carbonyl)oxy)propyl)phenoxy)acetic acid (synthesized according to previous method^2^) (15.0 mg, 0.030 mmol, 1.0 eq) and Propylamine (1.3 µL, 0.034 mmol, 1.1 eq) in DCM (2 mL), EDC (9.0 mg, 0.046 mmol, 1.5 eq), HOBt (7.0 mg, 0.046 mmol, 1.5 eq) and DIPEA (16 µL, 0.092 mmol, 3.0 eq) was added. The mixture was stirred at room temperature for 12 hours. After completion the reaction mixture was diluted with DCM (15 mL), the organic layer was washed with saturated sodium bicarbonate (aq), water and brine. The combined organic layer was dried over sodium sulfate, filtered, and concentrated under reduced pressure. The crude material was purified by PTLC (DCM/Methanol, 19:1) to obtain P300-C as a colorless oil (5.8 mg, 54 %).

**^1^H NMR** (400 MHz, MeOD) δ 7.34 (t, J = 7.8 Hz, 0.21H), 7.20 (t, J = 7.9 Hz, 1H), 7.07 – 6.94 (m, 1H), 6.95 – 6.84 (m, 2.24H), 6.81 – 6.79 (m, 1.28H), 6.77 – 6.72 (m, 1.36H), 6.69 (dd, J = 8.2, 2.0 Hz, 1H), 6.63 – 6.55 (m, 3.30H), 5.84 (dd, J = 7.9, 5.7 Hz, 0.2H), 5.61 (dd, J = 8.3, 5.4 Hz, 1H), 5.45 – 5.35 (m, 1H), 4.98 – 4.90 (m, 1H), 4.54 (s, 0.5H), 4.51 (s, 2H), 4.14 – 4.05 (m, 1.25H), 3.93 – 3.85 (m, 2H), 3.84 (s, 1.38H), 3.82 (s, 3H), 3.81 (s, 3H), 3.77 (s, 1H), 3.71 (s, 3H), 3.70 – 3.65 (m, 6H), 3.61 – 3.58 (m, 0.5H), 3.30 – 3.22 (m, 2.5H), 2.78 – 2.70 (m, 1.22H), 2.67 – 2.53 (m, 2H), 2.49 – 2.41 (m, 1.5H), 2.33 – 2.29 (m, 1.3H), 2.20 – 2.11 (m, 0.6H), 2.10 – 1.98 (m, 3H), 1.97 – 1.85 (m, 1.5H), 1.79 – 1.85 (m, 5H), 1.56 – 1.48 (m, 4H), 1.37 – 1.24 (m, 4H), 0.94 – 0.89 (m, 6H), 0.85 (t, J = 7.3 Hz, 1H). Note: rotomeric isomers observed.

**^13^C NMR** (101 MHz, MeOD) δ 173.76, 173.60, 170.48, 170.02, 169.51, 158.00, 157.79, 153.48, 153.19, 149.05, 147.48, 142.21, 136.55, 136.13, 135.52, 133.81, 133.75, 129.60, 129.44, 120.39, 120.31, 119.69, 119.30, 114.13, 113.68, 113.16, 112.89, 112.27, 111.88, 105.19, 104.61, 76.72, 75.74, 66.89, 59.68, 55.96, 55.15, 55.06, 52.23, 50.24, 49.87, 43.57, 37.84, 31.20, 30.86, 28.00, 26.21, 24.91, 20.44, 19.63, 12.70, 11.58, 11.27. *Note: rotomeric isomers observed*.

**HRMS** calculated for C_41_H_55_N_2_O_10_, 749.4008 (M+H^+^), found 749.4010.

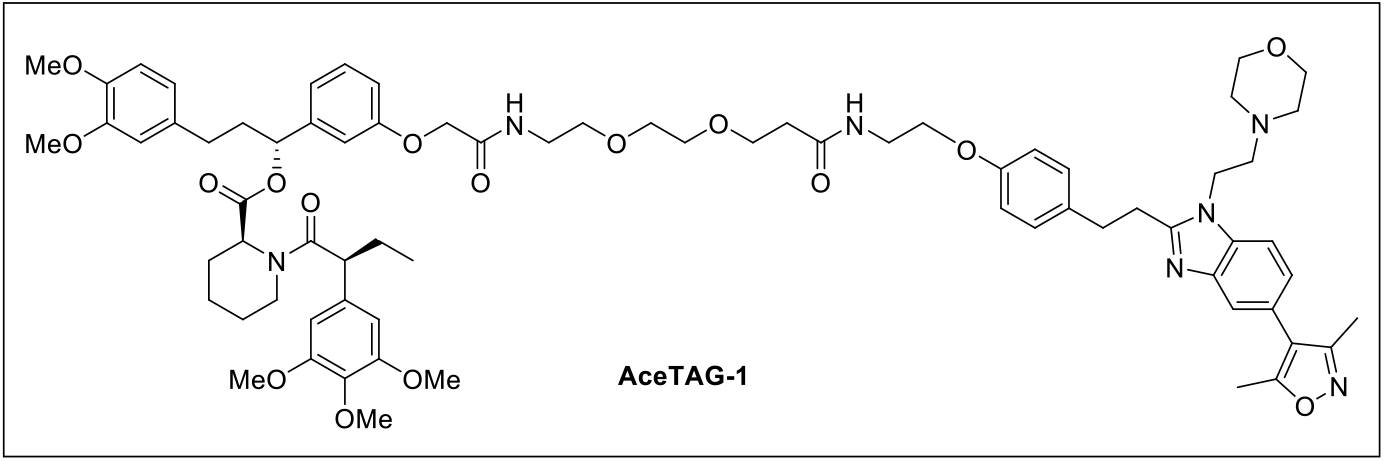

### (R)-3-(3,4-dimethoxyphenyl)-1-(3-((15-(4-(2-(5-(3,5-dimethylisoxazol-4-yl)-1-(2-morpholinoethyl)-1H-benzo[d]imidazol-2-yl)ethyl)phenoxy)-2,12-dioxo-6,9-dioxa-3,13-diazapentadecyl)oxy)phenyl)propyl (S)-1-((S)-2-(3,4,5-trimethoxyphenyl)butanoyl)piperidine-2-carboxylate (AceTAG-1)

To a solution of tert-butyl (2-(2-(3-((2-(4-(2-(5-(3,5-dimethylisoxazol-4-yl)-1-(2-morpholinoethyl)-1H-benzo[d]imidazol-2-yl)ethyl)phenoxy)ethyl)amino)-3-oxopropoxy)ethoxy)ethyl)carbamate (P300-PEG_2_-NHBoc) (20 mg, 0.027 mmol) in DCM (1 mL), a solution of 20 % TFA in DCM (1 mL) was added at 0 °C and resulting mixture was stirred at room temperature for 2 h. After completion (monitored by TLC), the reaction mixture was evaporated under reduced pressure to obtained the corresponding TFA salt of 3-(2-(2-aminoethoxy)ethoxy)-N-(2-(4-(2-(5-(3,5-dimethylisoxazol-4-yl)-1-(2-morpholinoethyl)-1H-benzo[d]imidazol-2-yl)ethyl)phenoxy)ethyl)propenamide (P300-PEG_2_-NH_2_.TFA), was obtained were used in next step without further purification. To a solution of corresponding TFA salt of 3-(2-(2-aminoethoxy)ethoxy)-N-(2-(4-(2-(5-(3,5-dimethylisoxazol-4-yl)-1-(2-morpholinoethyl)-1H-benzo[d]imidazol-2-yl)ethyl)phenoxy)ethyl)propenamide (5.0 mg, 0.0077 mmol, 1.0 eq) in DMF (1 mL), DIPEA (6.70 µL, 0.038 mmol, 5.0 eq.) and 2-(3-((*R*)-3-(3,4-dimethoxyphenyl)-1-(((*S*)-1-((*S*)-2-(3,4,5-trimethoxyphenyl)butanoyl)piperidine-2-carbonyl)oxy)propyl)phenoxy)acetic acid (synthesized as previously described^2^) (5.34 mg, 0.0077 mmol, 1 eq) was added followed by HATU (3.22 mg, 0.0085 mmol, 1.1 eq) were added at 0°C and resulting mixture was stirred for 5 minutes the corresponding starting material was fully consumed (indicated by TLC). The crude mixture was diluted with cold water and extracted in ethyl acetate (20 mlL X 3) then combined organic extract was dried over sodium sulfate, filtered, and concentrated under reduced pressure. The crude material was purified by PTLC (DCM/Methanol, 19:1) to obtain AceTAG-1 as a colorless sticky material (5.4 mg, 53 %).

**^1^H NMR** (600 MHz, CDCl_3_): δ 7.55 (d, J = 1.5 Hz, 1H), 7.29 (d, J = 8.2 Hz, 1H), 7.25 (t, J = 8.0 Hz, 0.25H), 7.14 – 7.05 (m, 4H), 7.03 – 6.98 (m, 1H), 6.94 (dt, J = 7.6, 1.2 Hz, 0.23H), 6.83 (dd, J = 2.6, 1.5 Hz, 0.22H), 6.80 – 6.77 (m, 0.27H), 6.77 – 6.74 (m, 2H), 6.73 – 6.71 (m, 0.36H), 6.70 – 6.68 (m, 2.6H), 6.67 – 6.63 (m, 1H), 6.62 – 6.57 (m, 3.3H), 6.35 (s, 0.4H), 6.34 (s, 2H), 5.72 (dd, J = 7.8, 6.0 Hz, 0.2H), 5.55 (dd, J = 8.3, 5.4 Hz, 1H), 5.41 – 5.33 (m, 1H), 4.59 (d, J = 5.6 Hz, 0.2H), 4.44 – 4.38 (m, 2H), 4.09 – 4.04 (m, 2H), 3.96 – 3.91 (m, 2H), 3.84 – 3.74 (m, 8H), 3.74–3.71 (m, 0.5H), 3.71 (s, 11H), 3.67 – 3.63 (m, 2.3H), 3.62 – 3.60 (m, 5H), 3.60 – 3.58 (m, 4.4H), 3.53 – 3.49 (m, 10H), 3.30 – 3.23 (m, 0.3H), 3.17 –3.14 (m, 2H), 3.12 – 3.06 (m, 2H), 2.73 (td, J = 13.4, 3.0 Hz, 1H), 2.63 – 2.53 (m, 2.2H), 2.52 – 2.45 (m, 1.4H), 2.44 – 2.38 (m, 7H), 2.36 (s, 3.3H), 2.26 – 2.20 (s, 4.3H), 2.05 – 1.95 (m, 2.6H), 1.88 – 1.82 (m, 2H), 1.69 – 1.58 (m, 4H), 1.56 – 1.51 (m, 1.6H), 1.42 – 1.34 (m, 3H), 1.31 (t, J = 7.4 Hz, 1H), 0.83 (t, J = 7.3 Hz, 3H), 0.78 (t, J = 7.3 Hz, 1.3H). *Note: rotomeric isomers observed*.

**^13^C NMR** (151 MHz, CDCl_3_): δ 172.70, 172.38, 171.66, 171.59, 170.62, 168.34, 168.10, 165.06, 159.03, 157.41, 157.30, 157.24, 157.21, 155.38, 155.35, 153.50, 153.20, 148.97, 148.89, 147.52, 147.38, 142.99, 142.37, 141.89, 136.96, 136.62, 135.98, 135.34, 134.28, 133.43, 133.33, 133.28 130.16, 129.88, 129.44, 124.25, 123.46, 120.20, 120.12, 119.97, 119.91, 119.81, 117.11, 114.68, 114.14, 113.82, 113.70, 112.98, 111.71, 111.48, 111.28, 109.46, 104.97, 104.58, 75.66, 70.27, 70.13, 69.80, 67.39, 67.17, 66.84, 65.91, 60.91, 60.79, 57.64, 56.31, 55.97, 55.93, 55.88, 55.86, 55.76, 54.88, 54.06, 52.08, 51.24, 50.79, 43.49, 41.52, 39.70, 38.92, 38.85, 38.82, 38.27, 37.99, 36.89, 36.86, 32.94, 31.94, 31.50, 31.30, 29.84, 29.72, 29.34, 28.37, 26.81, 26.50, 25.33, 24.52, 22.71, 20.94, 20.76, 17.58, 17.06, 14.15, 12.74, 12.59, 11.61, 10.92, 10.24. *Note: rotomeric isomers observed*.

**HRMS** calculated for C_73_H_94_N_7_O_16_, 1324.6752 (M+H^+^), found 1324.6749.

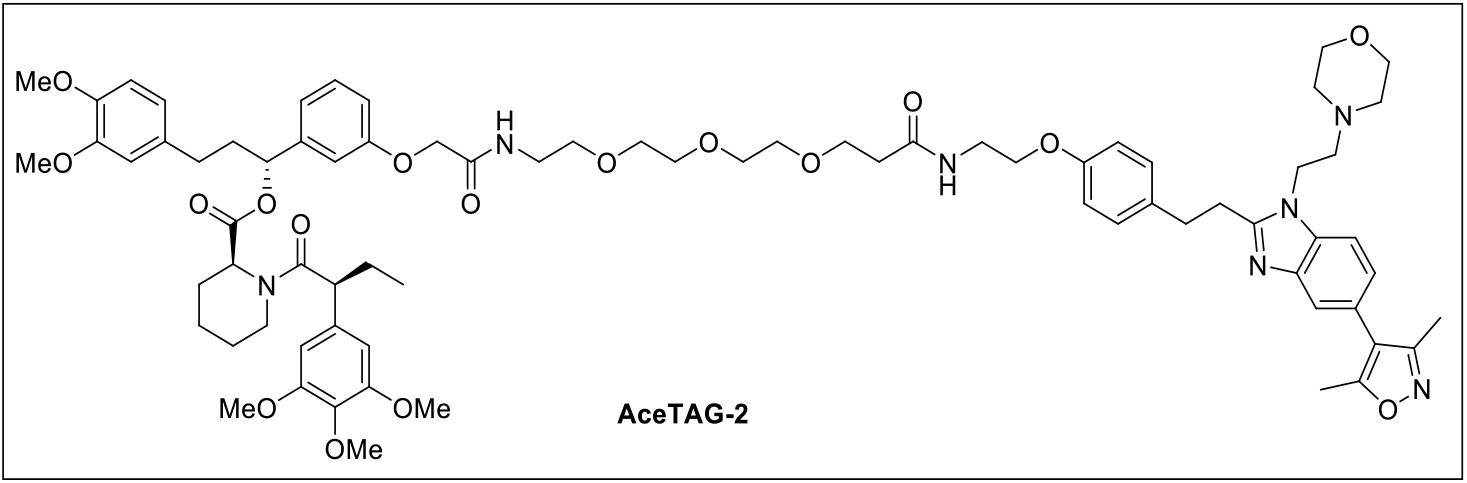

### (R)-3-(3,4-dimethoxyphenyl)-1-(3-((18-(4-(2-(5-(3,5-dimethylisoxazol-4-yl)-1-(2-morpholinoethyl)-1H-benzo[d]imidazol-2-yl)ethyl)phenoxy)-2,15-dioxo-6,9,12-trioxa-3,16-diazaoctadecyl)oxy)phenyl)propyl (S)-1-((S)-2-(3,4,5-trimethoxyphenyl)butanoyl)piperidine-2-carboxylate (AceTAG-2)

To a solution of tert-butyl (15-(4-(2-(5-(3,5-dimethylisoxazol-4-yl)-1-(2-morpholinoethyl)-1H-benzo[d]imidazol-2-yl)ethyl)phenoxy)-12-oxo-3,6,9-trioxa-13-azapentadecyl)carbamate (P300-PEG_3_-NHBoc) (10 mg, 0.013 mmol) in DCM (1 mL), a solution of 20 % TFA in DCM (1 mL) was added at 0 °C and resulting mixture was stirred at room temperature for 2 h. After completion (monitored by TLC), the reaction mixture was evaporated under reduced pressure to obtained the corresponding TFA salt of 3-(2-(2-(2-aminoethoxy)ethoxy)ethoxy)-N-(2-(4-(2-(5-(3,5-dimethylisoxazol-4-yl)-1-(2-morpholinoethyl)-1H-benzo[d]imidazol-2-yl)ethyl)phenoxy)ethyl)propenamide, was obtained were used in next step without further purification. To a solution of corresponding TFA salt of 3-(2-(2-(2-aminoethoxy)ethoxy)ethoxy)-N-(2-(4-(2-(5-(3,5-dimethylisoxazol-4-yl)-1-(2-morpholinoethyl)-1H-benzo[d]imidazol-2-yl)ethyl)phenoxy)ethyl)propanamide (6.0 mg, 0.0087 mmol, 1 eq.) in DMF (1 mL), DIPEA (7.52 µL, 0.043 mmol, 5 eq.) and 2-(3-((*R*)-3-(3,4-dimethoxyphenyl)-1-(((*S*)-1-((*S*)-2-(3,4,5-trimethoxyphenyl)butanoyl)piperidine-2-carbonyl)oxy)propyl)phenoxy)acetic acid (synthesized as previously described^2^) (6.0 mg, 0.0087 mmol, 1 eq) was added followed by HATU (3.16 mg, 0.0083 mmol, 1.1 eq) were added at 0°C and resulting mixture was stirred for 5 minutes, the corresponding starting material was fully consumed (indicated by TLC). The crude mixture was diluted with cold water and extracted in ethyl acetate (20 mL X 3) then combined organic extract was dried over sodium sulfate, filtered, and concentrated under reduced pressure. The crude material was purified by PTLC (DCM/Methanol, 19:1) to obtain AceTAG-2 as a colorless sticky material (5.6 mg, 48 %).

**^1^H NMR** (600 MHz, CDCl_3_): δ 7.62 (d, J = 1.4 Hz, 1H), 7.35 (d, J = 8.2 Hz, 1H), 7.32 (t, J = 7.9 Hz, 0.3H), 7.22 – 7.11 (m, 4H), 7.08 (d, J = 5.9 Hz, 1H), 7.00 (dt, J = 7.6, 1.2 Hz, 0.3H), 6.90 (d, J = 2.4 Hz, 0.3H), 6.86 (dd, J = 8.4, 2.7, Hz, 0.5H), 6.85 – 6.80 (m, 3H), 6.80 – 6.74 (m, 3H), 6.71 – 6.63 (m, 3H), 6.41 (s, 0.4H), 6.40 (s, 2H), 5.81 – 5.76 (m, 0.23H), 5.62 (dd, J = 8.3, 5.4 Hz, 1H), 5.46 – 5.42 (m, 1H), 4.66 – 4.60 (m, 0.5H), 4.50 – 4.45 (m, 2.2H), 4.13 (t, J = 6.8 Hz, 2H), 4.01 (t, J = 5.4 Hz, 2H), 3.87 – 3.82 (m, 8.5H), 3.80 – 3.76 (m, 3.2H), 3.72 – 3.69 (m, 2.5H), 3.69 – 3.67 (m, 5H), 3.67 – 3.64 (m, 4.3H), 3.63 – 3.61 (m, 3H), 3.61 – 3.56 (m, 11.4H), 3.55 – 3.51 (m, 2.2H), 3.35 – 3.32 (m, 0.4H), 3.25 – 3.20 (m, 2H), 3.19 – 3.14 (m, 2.2H), 2.81 – 2.76 (m, 1H), 2.62 (t, J = 6.9 Hz, 2H), 2.58 – 2.51 (m, 1.3H), 2.49 – 2.43 (m, 7H), 2.43 – 2.40 (m, 3.4H), 2.33 – 2.26 (m, 4H), 2.17 (s, 0.25H), 2.13 – 2.04 (m, 2H), 2.02 – 1.98 (m, 1H), 1.95 – 1.88 (m, 1H), 1.83 – 1.79 (m, 0.3H), 1.61 – 1.59 (m, 1.3H), 1.45 – 1.40 (m, 1H), 0.89 (t, J = 7.3 Hz, 3H), 0.84 (t, J = 7.3 Hz, 1H). *Note: rotomeric isomers observed*.

**^13^C NMR** (151 MHz, CDCl_3_) δ 172.67, 172.36, 171.74, 171.69, 170.61, 170.49, 168.21, 168.00, 165.05, 159.03, 157.44, 157.31, 157.26, 155.38, 153.49, 153.20, 148.97, 148.89, 147.52, 147.37, 143.06, 142.34, 141.86, 136.95, 136.62, 135.99, 135.34, 134.30, 133.39, 133.34, 133.08, 130.14, 129.86, 129.42, 124.23, 123.44, 120.19, 120.12, 119.94, 119.77, 117.11, 114.68, 114.28, 113.94, 113.61, 112.88, 111.70, 111.60, 111.35, 111.28, 109.44, 104.97, 104.57, 75.66, 70.51, 70.37, 70.31, 70.27, 69.74, 67.42, 67.38, 67.20, 66.84, 66.80, 60.91, 60.79, 57.65, 56.31, 55.97, 55.93, 55.88, 55.86, 55.77, 54.07, 52.07, 51.24, 50.80, 43.48, 41.52, 39.69, 38.88, 38.83, 38.29, 38.00, 36.90, 32.94, 31.50, 31.29, 30.96, 29.87, 29.72, 28.38, 26.82, 25.34, 24.52, 22.71, 22.63, 20.95, 20.77, 12.74, 12.59, 11.61, 10.92. *Note: rotomeric isomers observed*.

**HRMS** calculated for C_75_H_98_N_7_O_17_, 1368.7014 (M+H^+^), found 1368.7018.

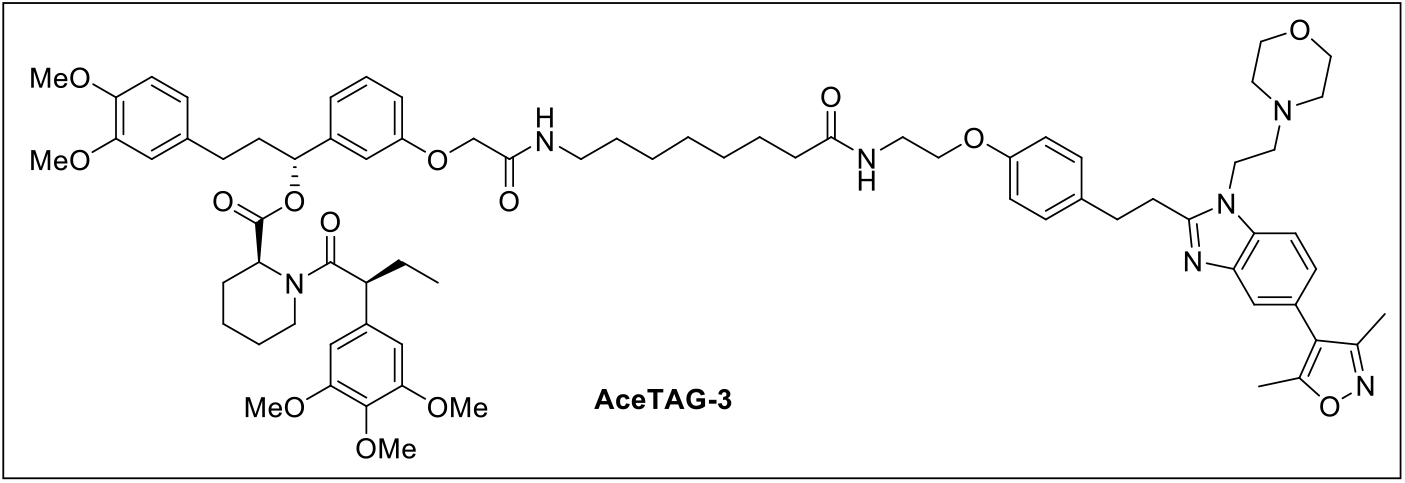

### (R)-3-(3,4-dimethoxyphenyl)-1-(3-(2-((8-((2-(4-(2-(5-(3,5-dimethylisoxazol-4-yl)-1-(2-morpholinoethyl)-1H-benzo[d]imidazol-2-yl)ethyl)phenoxy)ethyl)amino)-8-oxooctyl)amino)-2-oxoethoxy)phenyl)propyl (S)-1-((S)-2-(3,4,5-trimethoxyphenyl)butanoyl)piperidine-2-carboxylate (AceTAG-3)

To a solution of tert-butyl (8-((2-(4-(2-(5-(3,5-dimethylisoxazol-4-yl)-1-(2-morpholinoethyl)-1H-benzo[d]imidazol-2-yl)ethyl)phenoxy)ethyl)amino)-8-oxooctyl)carbamate (P300-C7-NHBoc) (15 mg, 0.020 mmol) in DCM (1 mL), a solution of 20 % TFA in DCM (1 mL) was added at 0 °C and resulting mixture was stirred at room temperature for 2 h. After completion (monitored by TLC), the reaction mixture was evaporated under reduced pressure to obtained the corresponding TFA salt of 8-amino-N-(2-(4-(2-(5-(3,5-dimethylisoxazol-4-yl)-1-(2-morpholinoethyl)-1H-benzo[d]imidazol-2-yl)ethyl)phenoxy)ethyl)octanamide was obtained, were used in next step without further purification. To a solution of corresponding TFA salt of 8-amino-N-(2-(4-(2-(5-(3,5-dimethylisoxazol-4-yl)-1-(2-morpholinoethyl)-1H-benzo[d]imidazol-2-yl)ethyl)phenoxy)ethyl)octanamide (8.0 mg, 0.013 mmol, 1 eq.) in DCM (1 mL), DIPEA (7.52 µL, 0.043 mmol, 5 eq.) and 2-(3-((*R*)-3-(3,4-dimethoxyphenyl)-1-(((*S*)-1-((*S*)-2-(3,4,5-trimethoxyphenyl)butanoyl)piperidine-2-carbonyl)oxy)propyl)phenoxy)acetic acid (synthesized as previously described^2^) (8.8 mg, 0.013 mmol, 1 eq.) was added followed by EDC (3.65 mg, 0.019 mmol, 1.5 eq.) and HOBt (2.60 mg, 0.019 mmol, 1.5 eq.) were added at 0°C and resulting mixture was stirred for 10 hours, the corresponding starting material was fully consumed (indicated by TLC). The crude mixture was diluted with cold water and extracted in DCM (20 mL X 3) then combined organic extract was dried over sodium sulfate, filtered, and concentrated under reduced pressure. The crude material was purified by PTLC (DCM/Methanol, 19:1) to obtain AceTAG-3 as a colorless sticky material (7.8 mg, 47 %).

**^1^H NMR** (600 MHz, CDCl_3_): δ 7.64 (s, 1H), 7.36 (s, 1H), 7.34 – 7.31 (m, 0.4H), 7.23 – 7.11 (m, 4H), 7.00 (d, J = 7.6 Hz, 0.25H), 6.91 (s, 0.25H), 6.87 – 6.82 (m, 2.3H), 6.81 – 6.77 (m, 0.3H), 6.78 – 7.74 (m, 3H), 6.71 – 6.60 (m, 4H), 6.41 (s, 0.4H), 6.40 (s, 2H), 6.01 – 5.99 (m, 1H), 5.81 – 5.79 (m, 0.2H), 5.62 (dd, J = 8.3, 5.4 Hz, 1H), 5.48 – 5.43 (m, 1H), 4.70 – 4.64 (m, 0.25H), 4.66 – 4.58 (m, 0.6H), 4.50 – 4.44 (m, 2H), 4.17 – 4.09 (m, 2H), 4.01 (t, J = 5.1 Hz, 2H), 3.86 – 3.81 (m, 9H), 3.80 – 3.76 (s, 3.5H), 3.67 – 3.63 (m, 11H), 3.60 – 3.56 (m, 1H), 3.36 – 3.28 (m, 2.5H), 3.26 – 3.21 (m, 2.5H), 3.20 – 3.15 (m, 2.3H), 2.82 – 2.76 (m, 1H), 2.65 – 2.60 (m, J = 6.9 Hz, 2.4H), 2.58 – 2.52 (m, 1.6H), 2.49 – 2.44 (m, 4.5H), 2.43 – 2.39 (m, 4.2H), 2.33 – 2.25 (m, 4.5H), 2.20 – 2.14 (m, 2.4H), 2.13 – 2.03 (m, 2.4H), 1.95 – 1.89 (m, 1H), 1.74– 1.66 (m, 4H), 1.63 – 1.59 (m, 6H), 1.55 – 1.49 (m, 3H), 1.47 – 1.41 (m, 1.3H), 1.33 – 1.23 (m, 12H), 0.90 (t, J = 7.3 Hz, 3H), 0.85 (t, J = 7.3 Hz, 1.3H). *Note: rotomeric isomers observed*.

**^13^C NMR** (151 MHz, CDCl_3_) δ 173.30, 172.70, 172.39, 170.66, 170.47, 168.03, 167.78, 165.06, 159.03, 157.43, 157.21, 155.39, 153.52, 153.22, 148.98, 148.91, 147.55, 147.40, 143.06, 142.39, 141.92, 137.00, 136.64, 136.01, 135.35, 134.31, 133.50, 133.37, 133.11, 130.17, 129.86, 129.47, 124.25, 123.46, 120.23, 120.15, 120.03, 119.95, 119.82, 117.12, 114.68, 113.83, 113.53, 113.09, 111.74, 111.65, 111.32, 109.45, 104.99, 104.61, 76.57, 67.39, 66.93, 66.85, 60.92, 60.80, 57.66, 56.33, 55.96, 55.88, 55.74, 54.08, 52.09, 51.26, 50.82, 43.49, 41.53, 39.69, 39.00, 38.92, 38.25, 37.97, 36.58, 32.96, 31.95, 31.31, 29.88, 29.73, 29.54, 29.09, 28.85, 28.40, 26.82, 26.63, 26.48, 25.53, 25.35, 24.54, 22.72, 20.93, 20.74, 14.16, 12.76, 12.61, 11.62, 10.93. *Note: rotomeric isomers observed*.

**HRMS** calculated for C_74_H_96_N_7_O_14_, 1306.7010 (M+H^+^), found 1306.7008.

## Spectra

### ^1^H NMR (P300-PEG2-NHBoc)

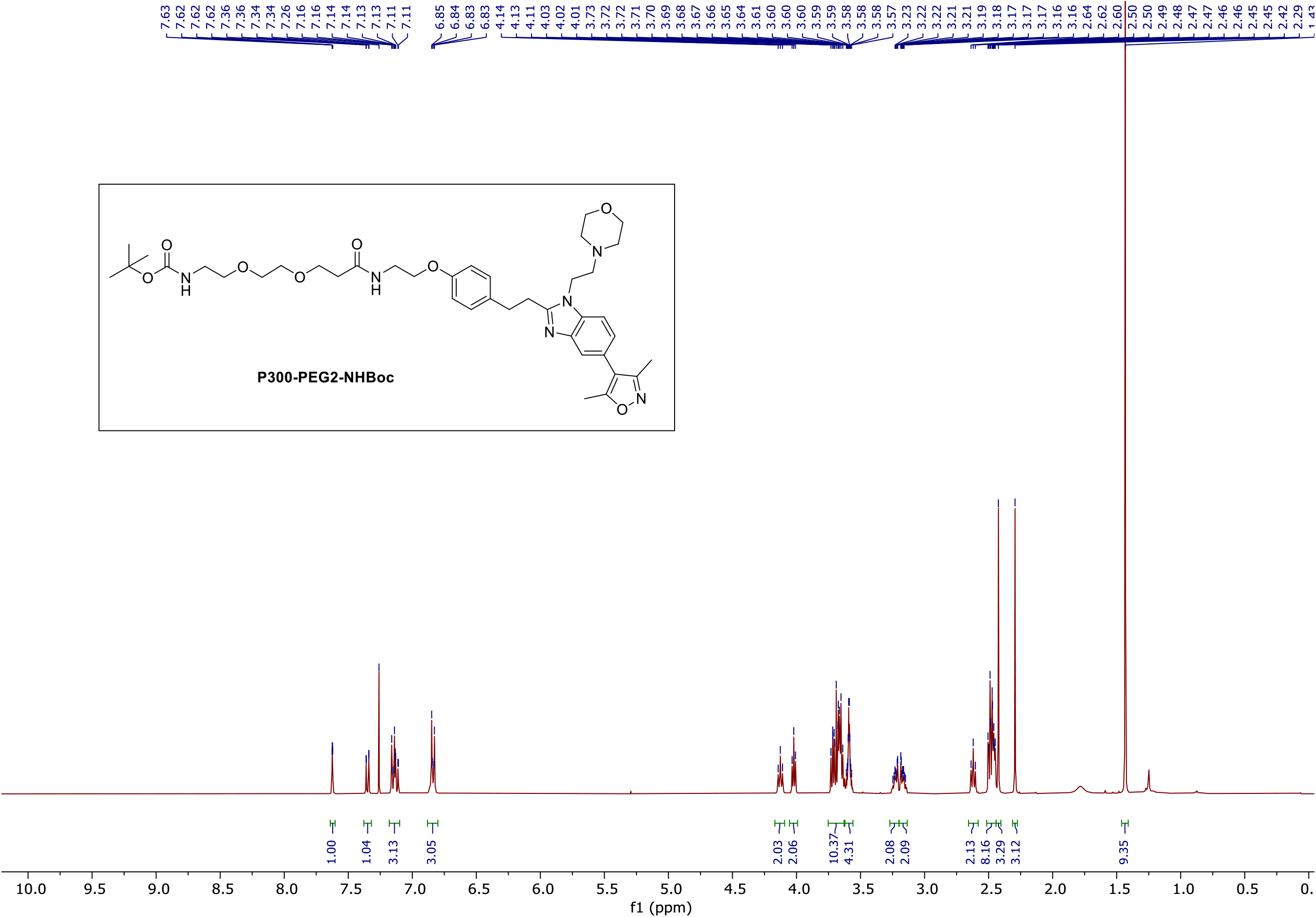

### ^13^C NMR (P300-PEG2-NHBoc)

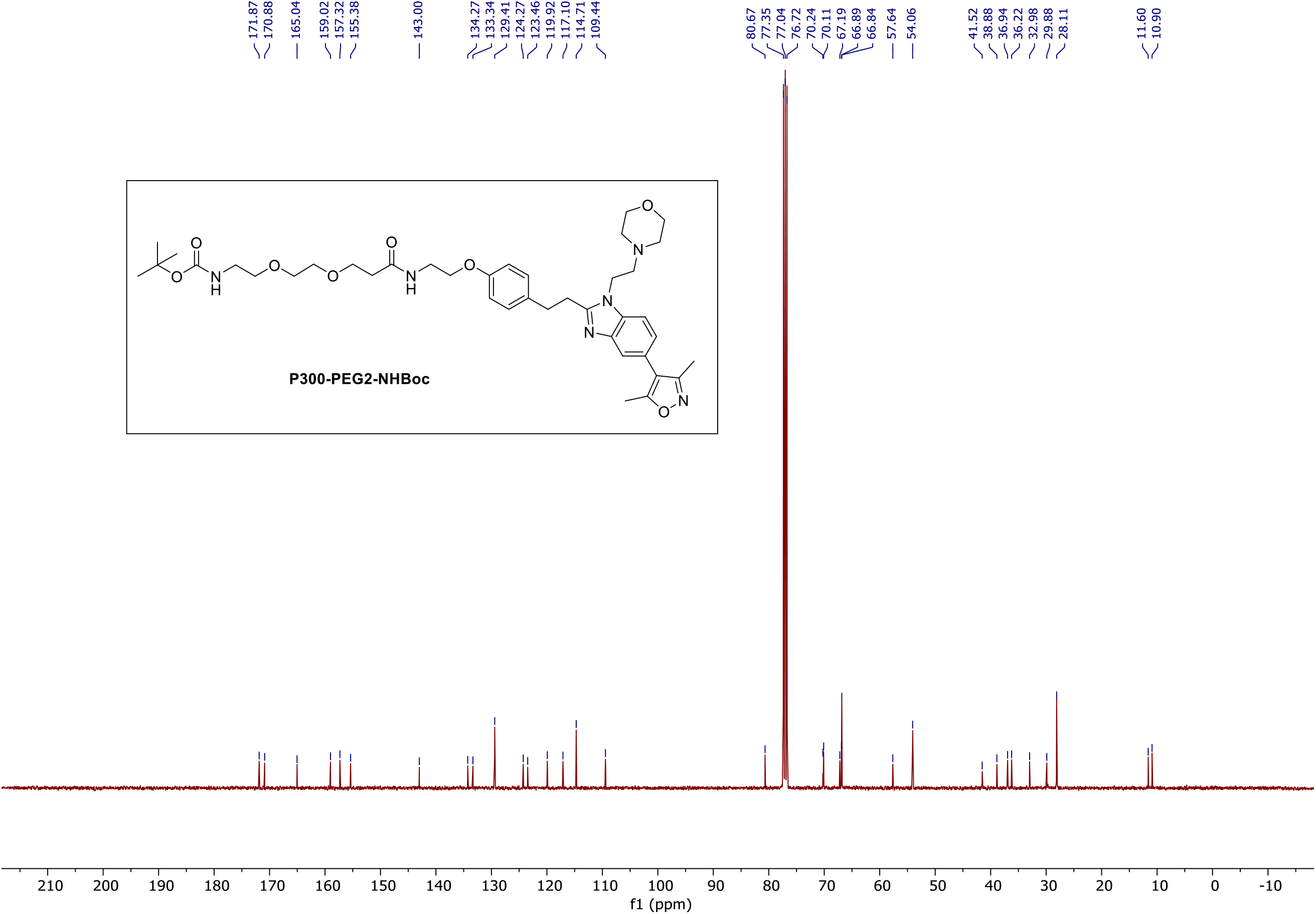

### ^1^H NMR (P300-PEG3-NHBoc)

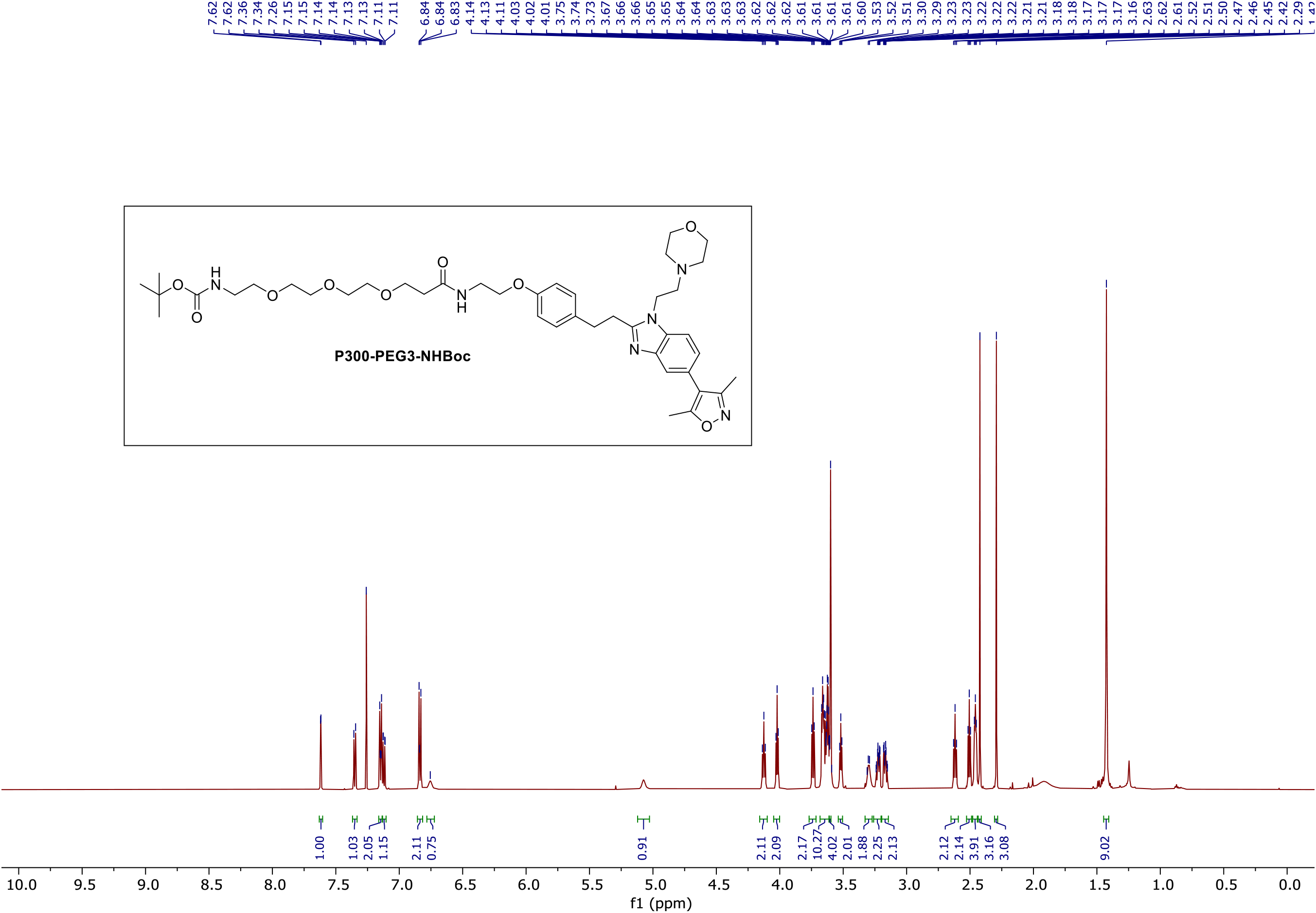

### ^13^C NMR (P300-PEG3-NHBoc)

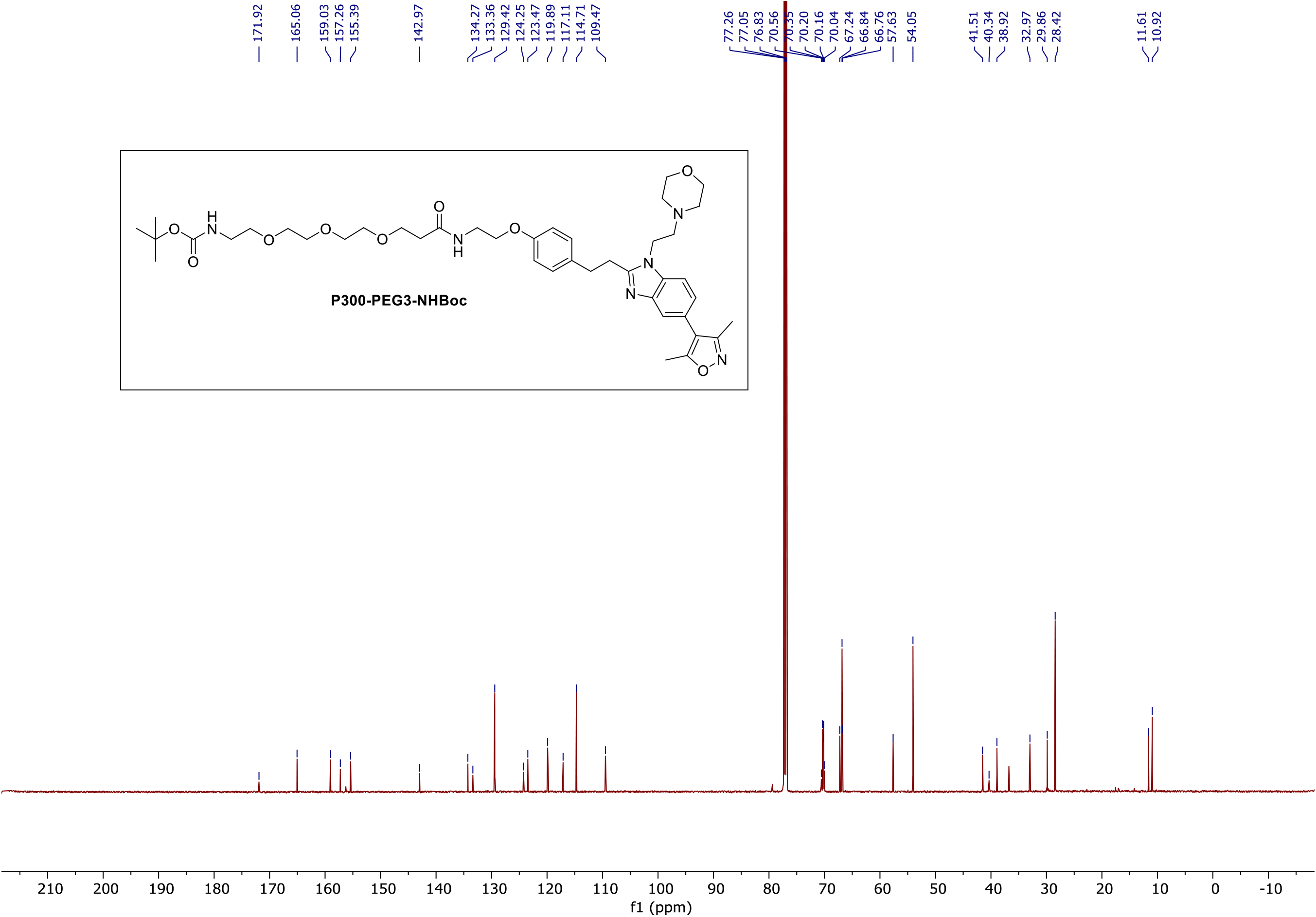

### ^1^H NMR (P300-C7-NHBoc)

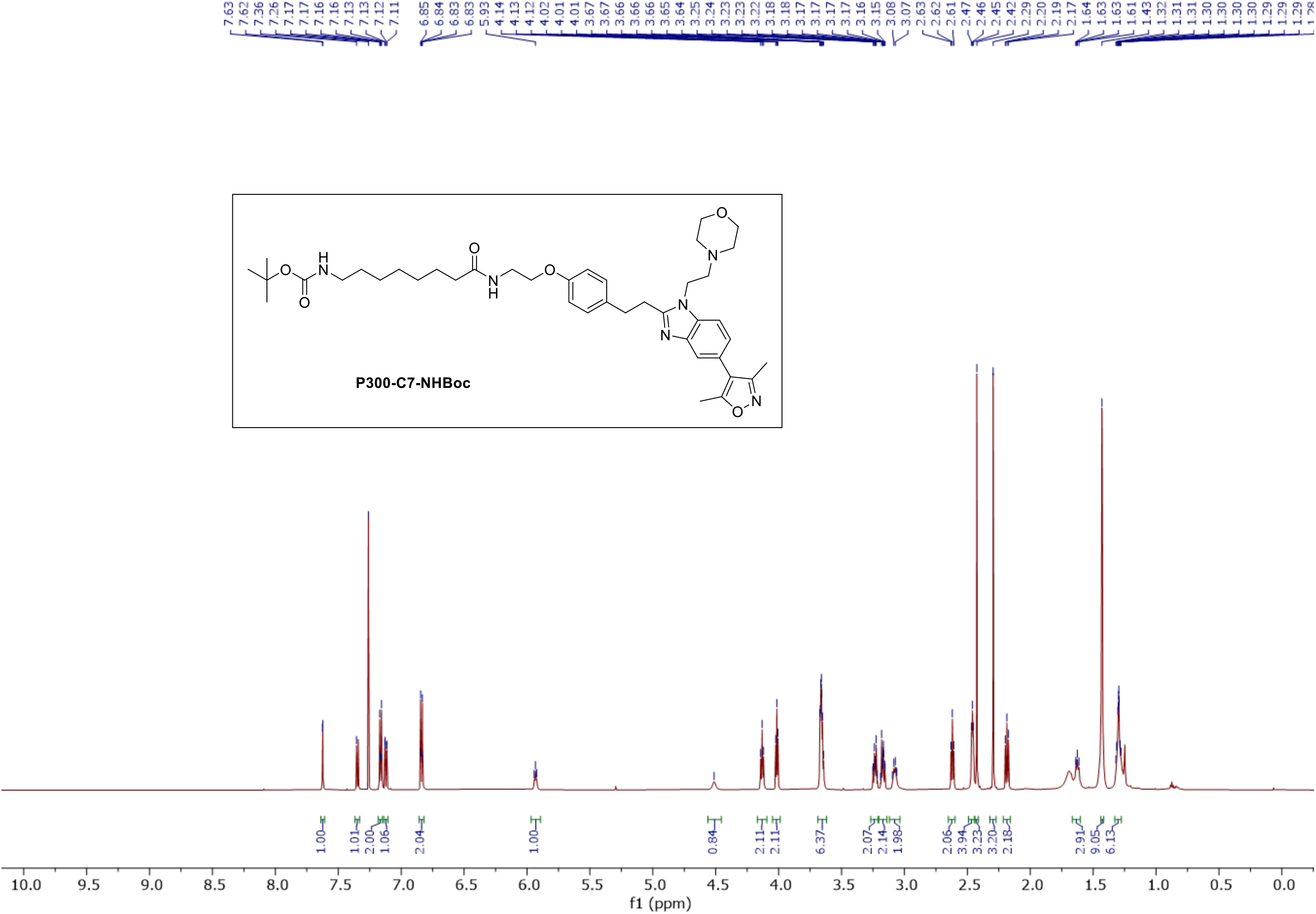

### ^13^C NMR (P300-C7-NHBoc)

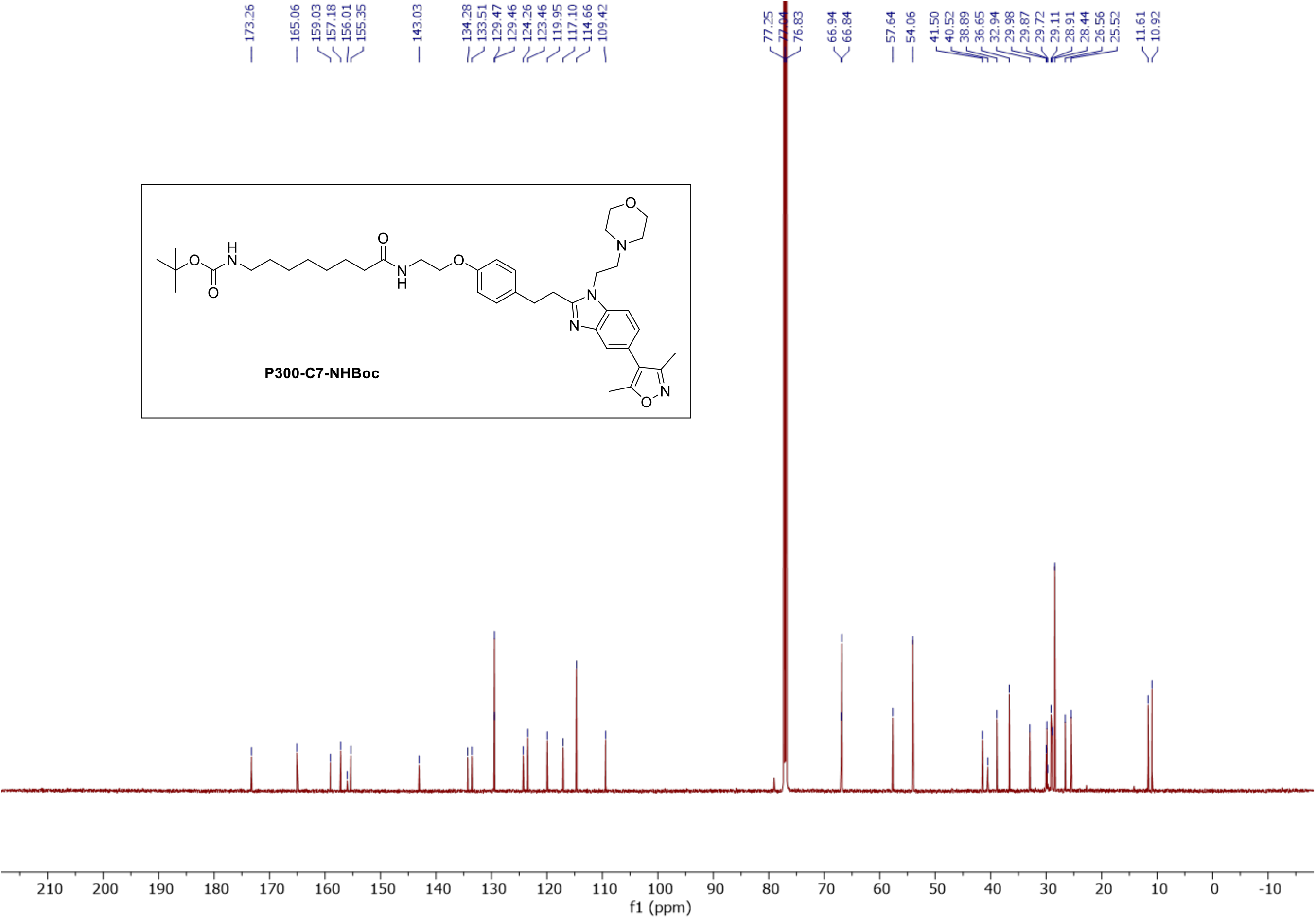

### ^1^H NMR (P300-P)

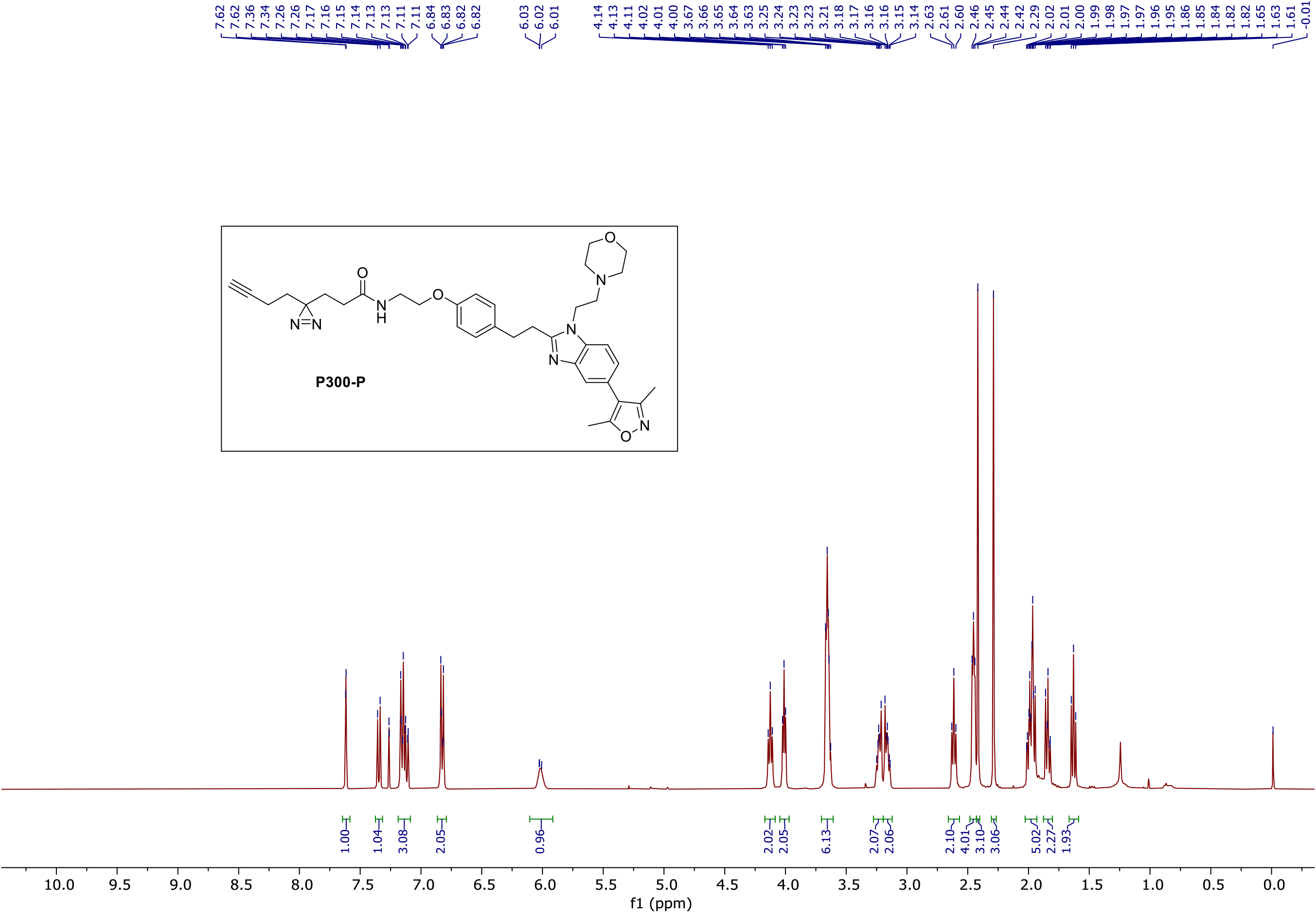

### ^13^C NMR (P300-P)

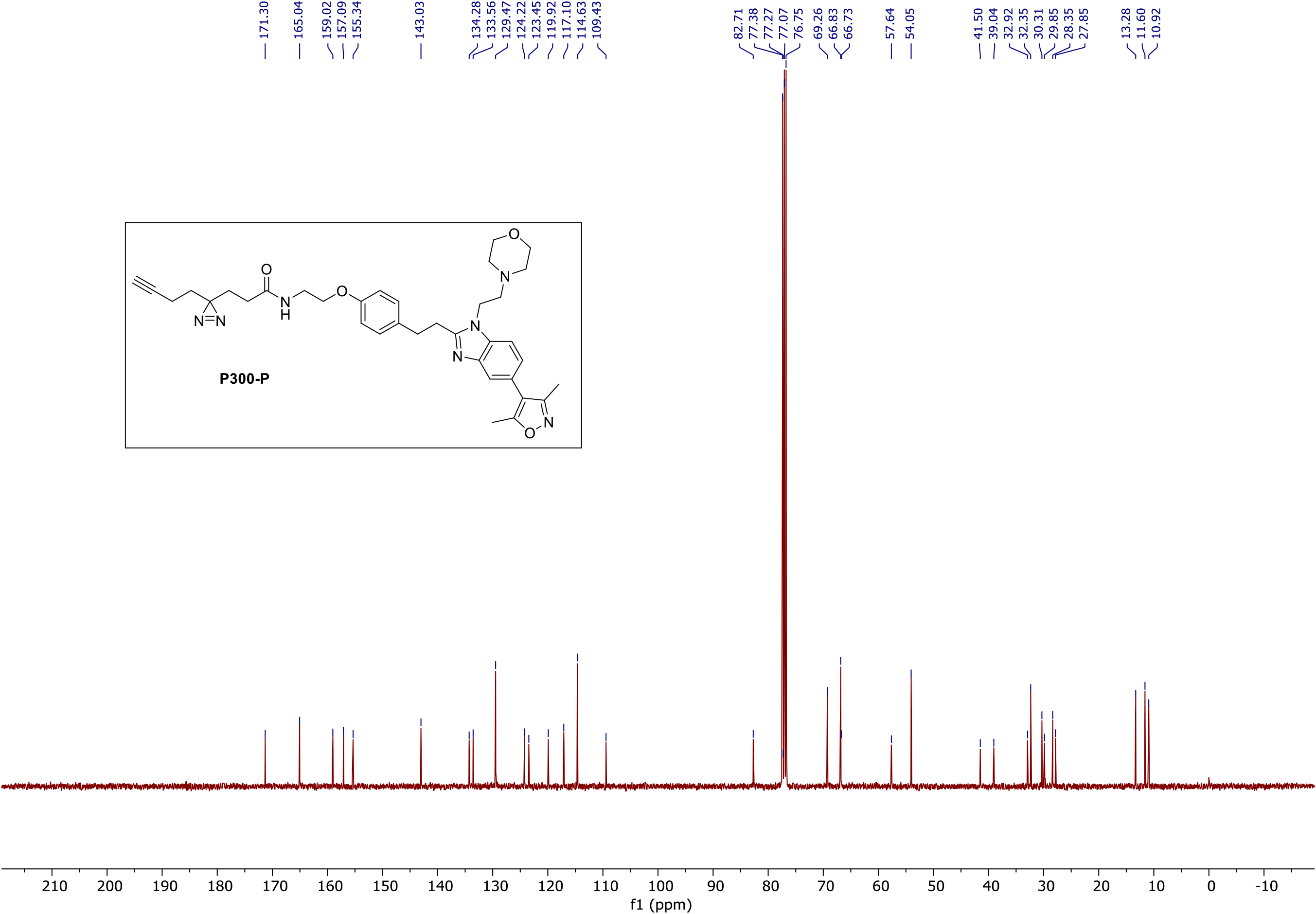

### ^1^H NMR (P300-C)

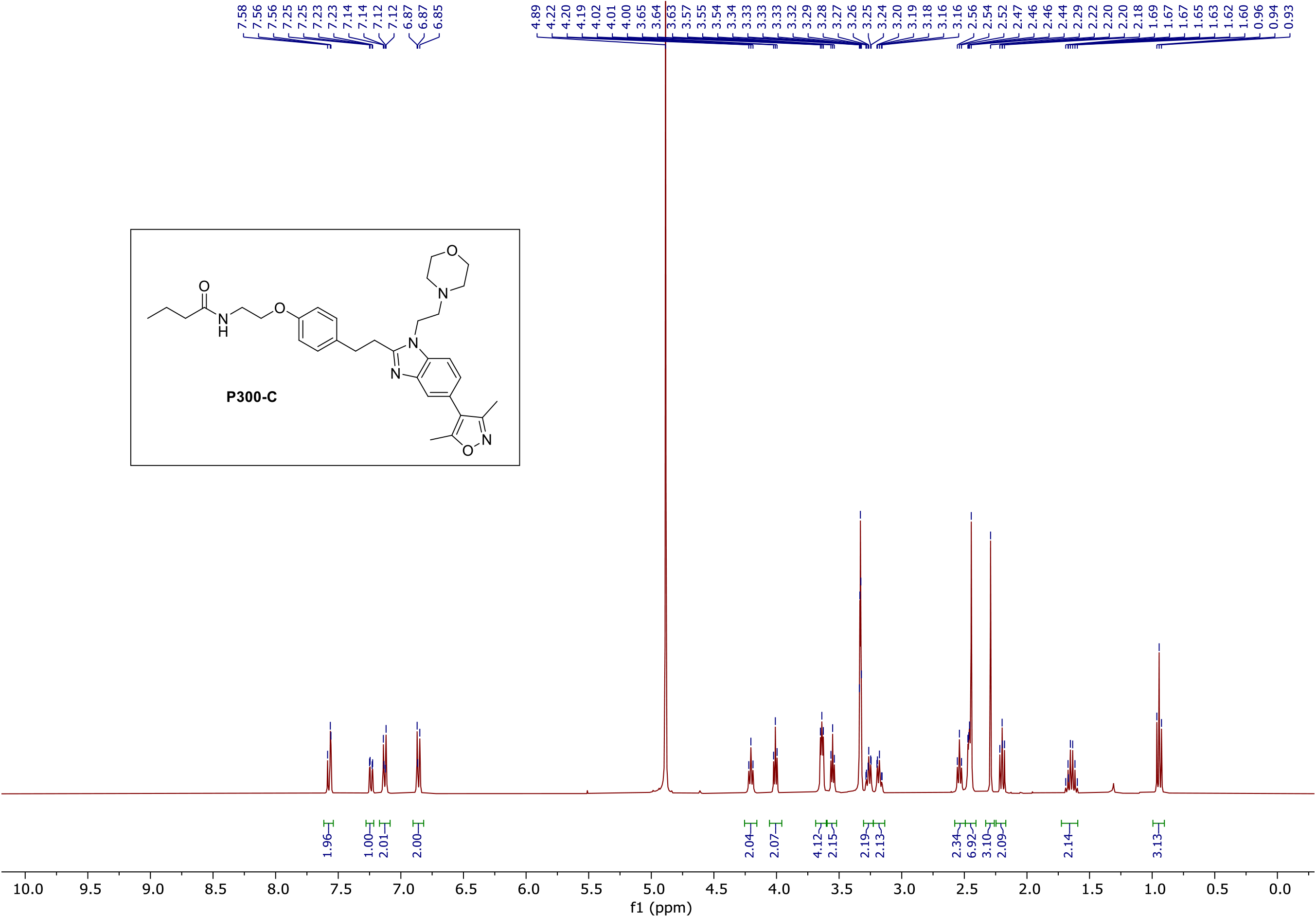

### ^13^C NMR (P300-C)

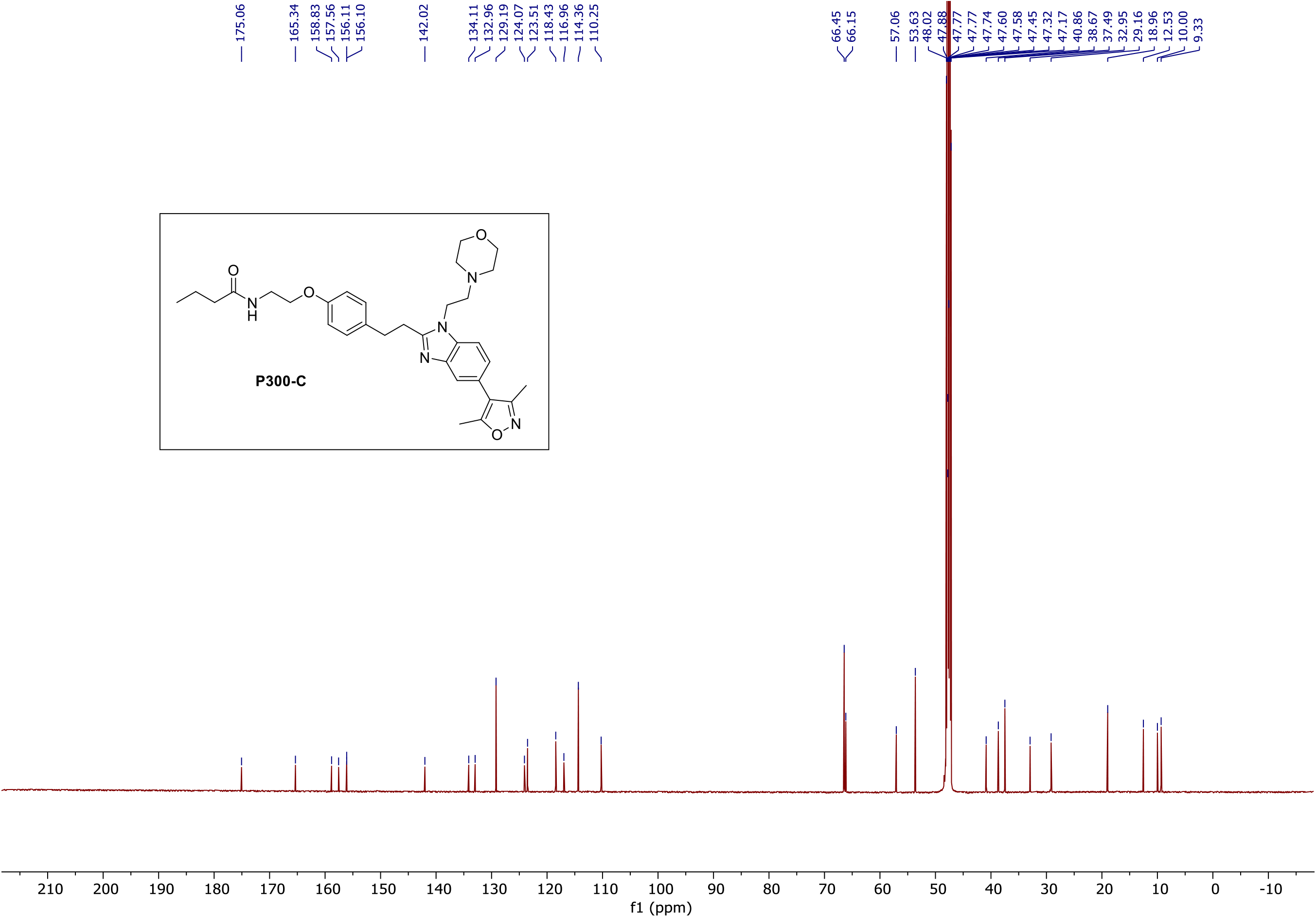

### ^1^H NMR (FKBP-P)

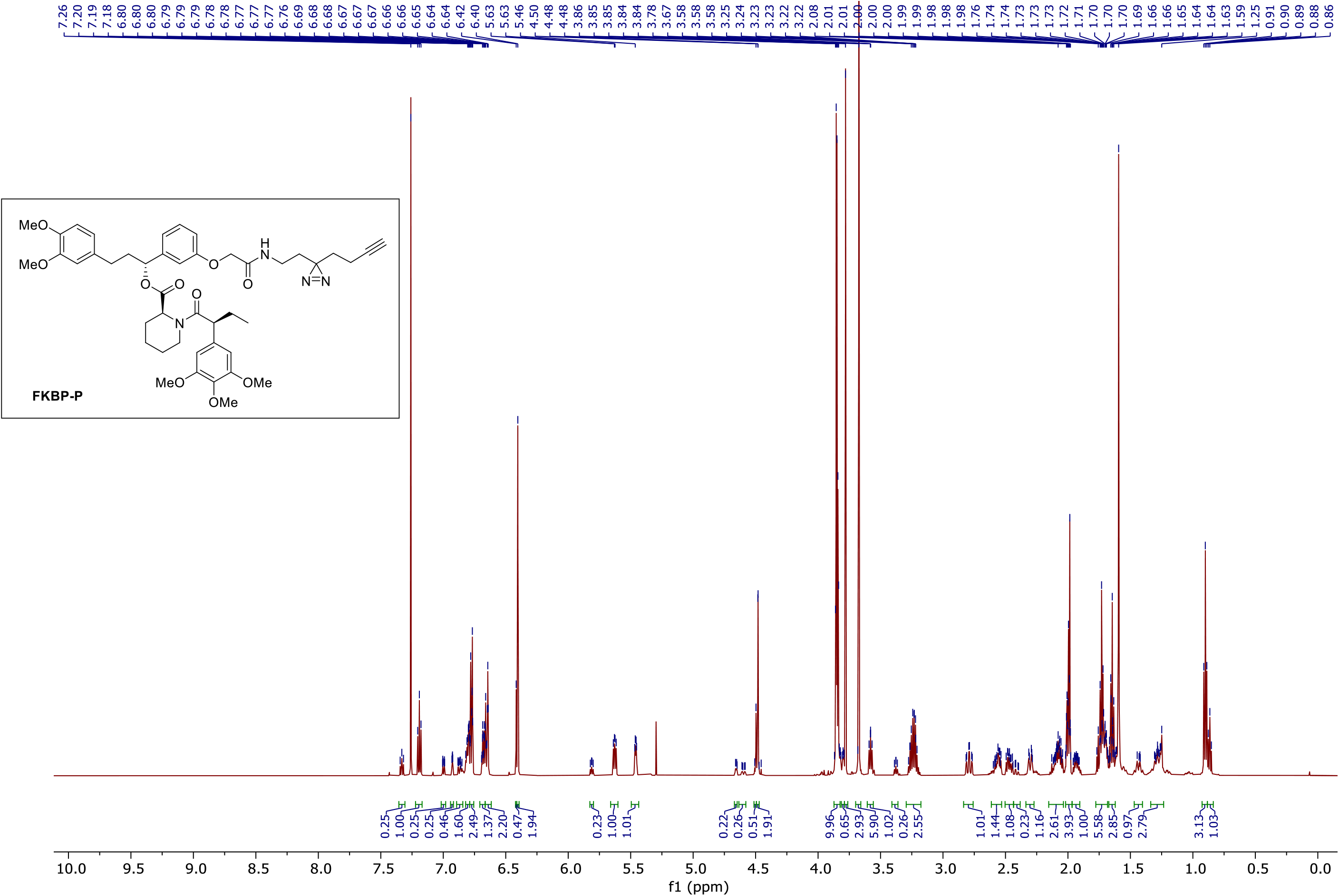

### ^13^C NMR (FKBP-P)

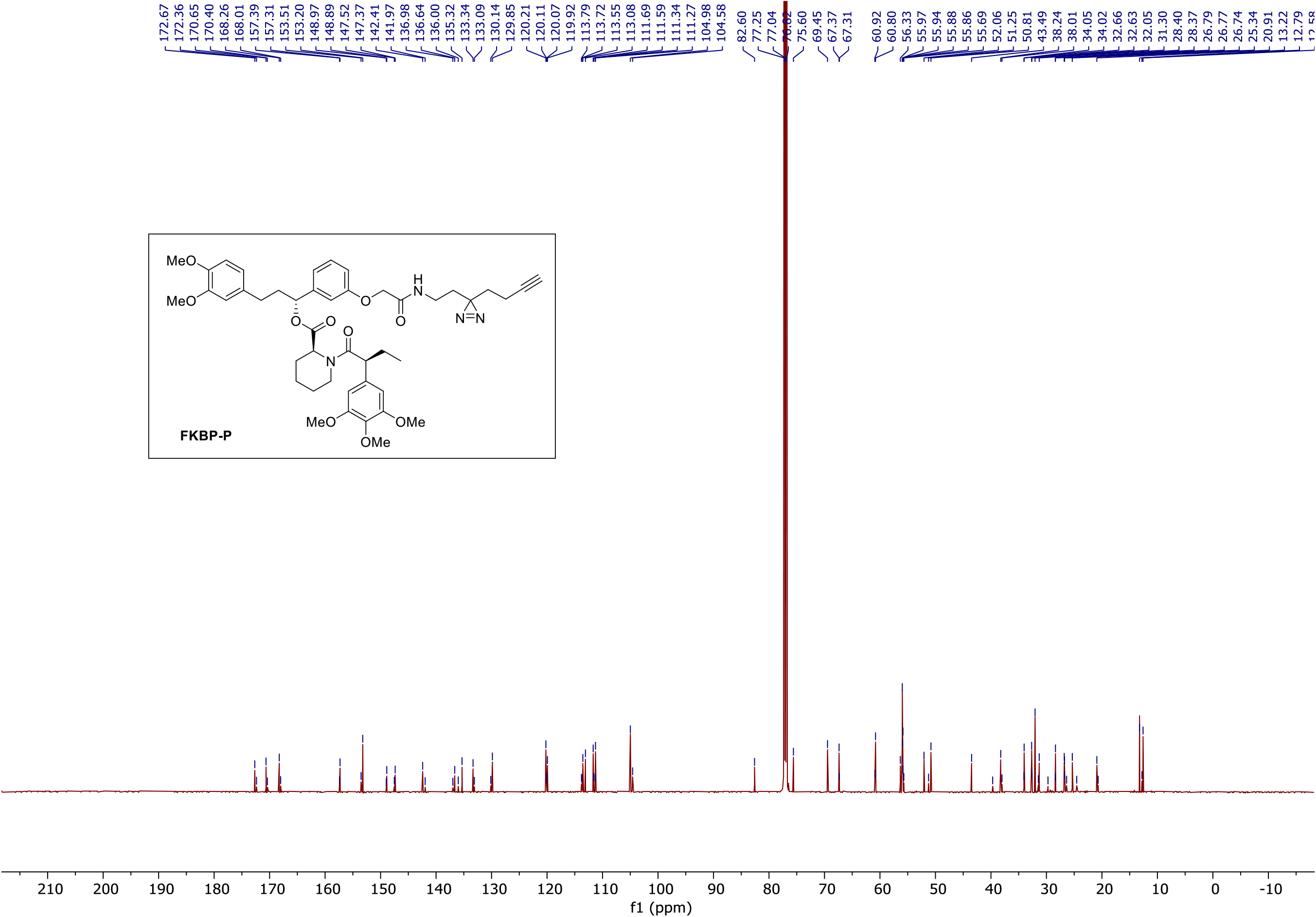

### ^1^H NMR (FKBP-C)

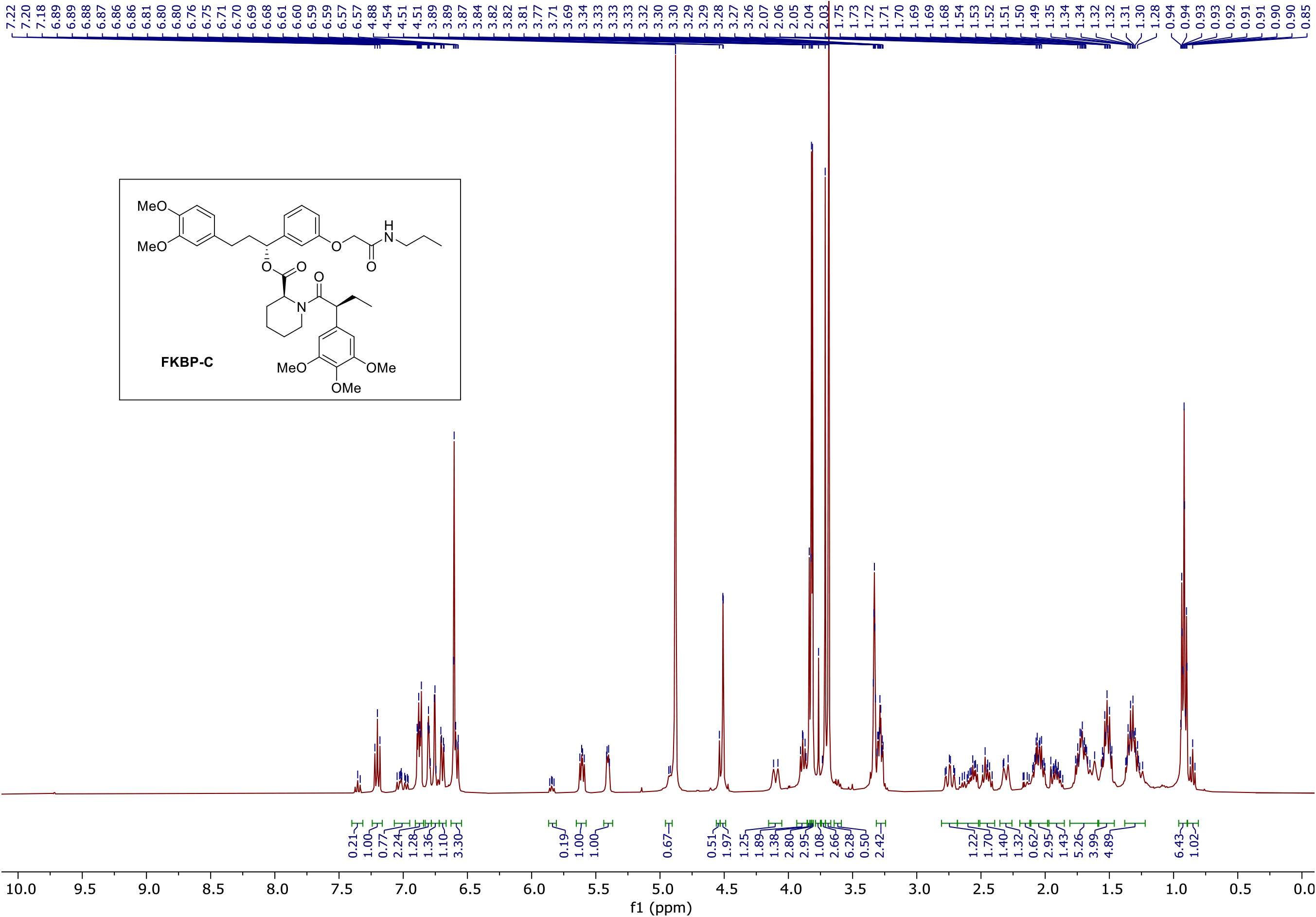

### ^13^C NMR (FKBP-C)

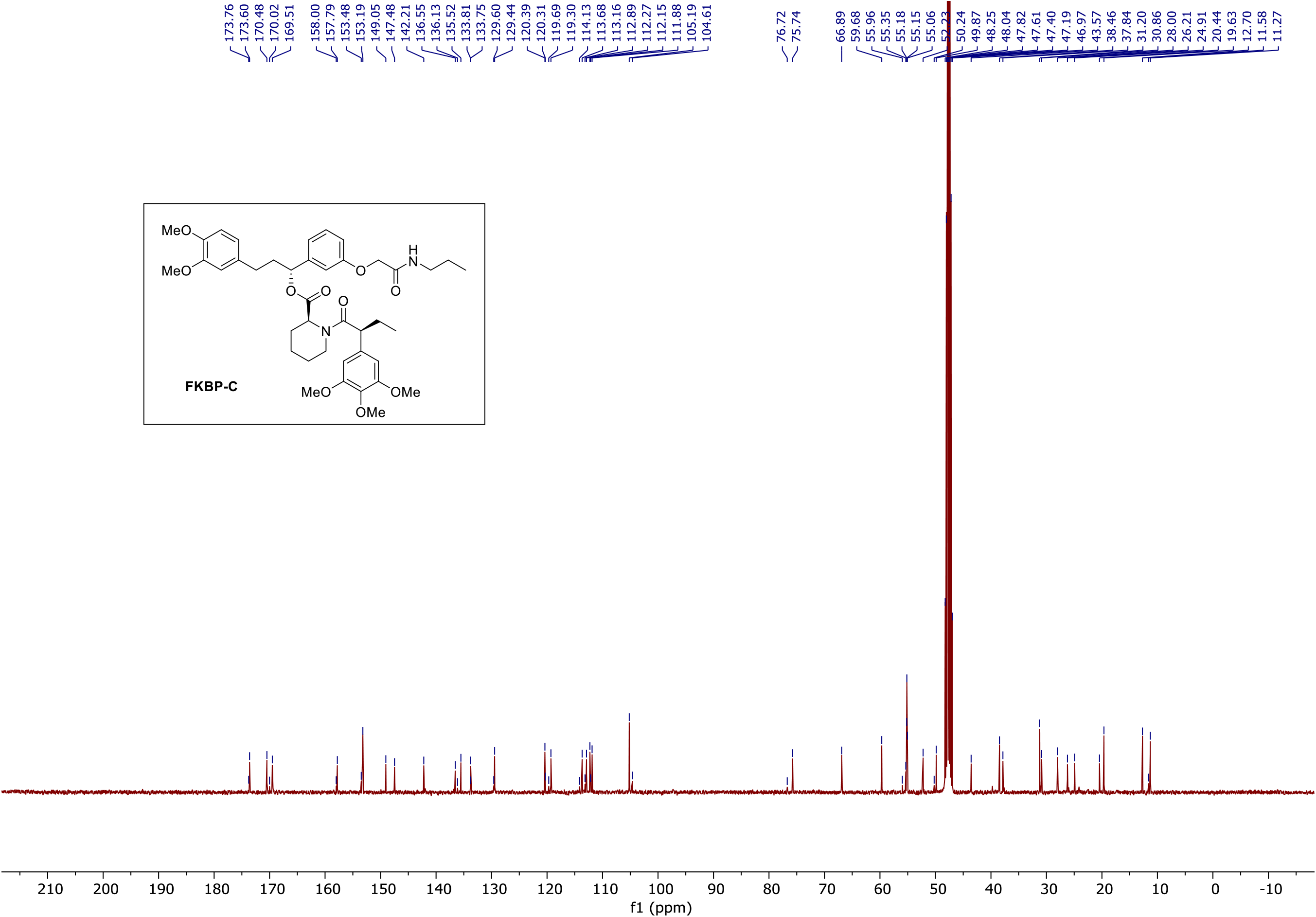

### ^1^H NMR (AceTAG-1)

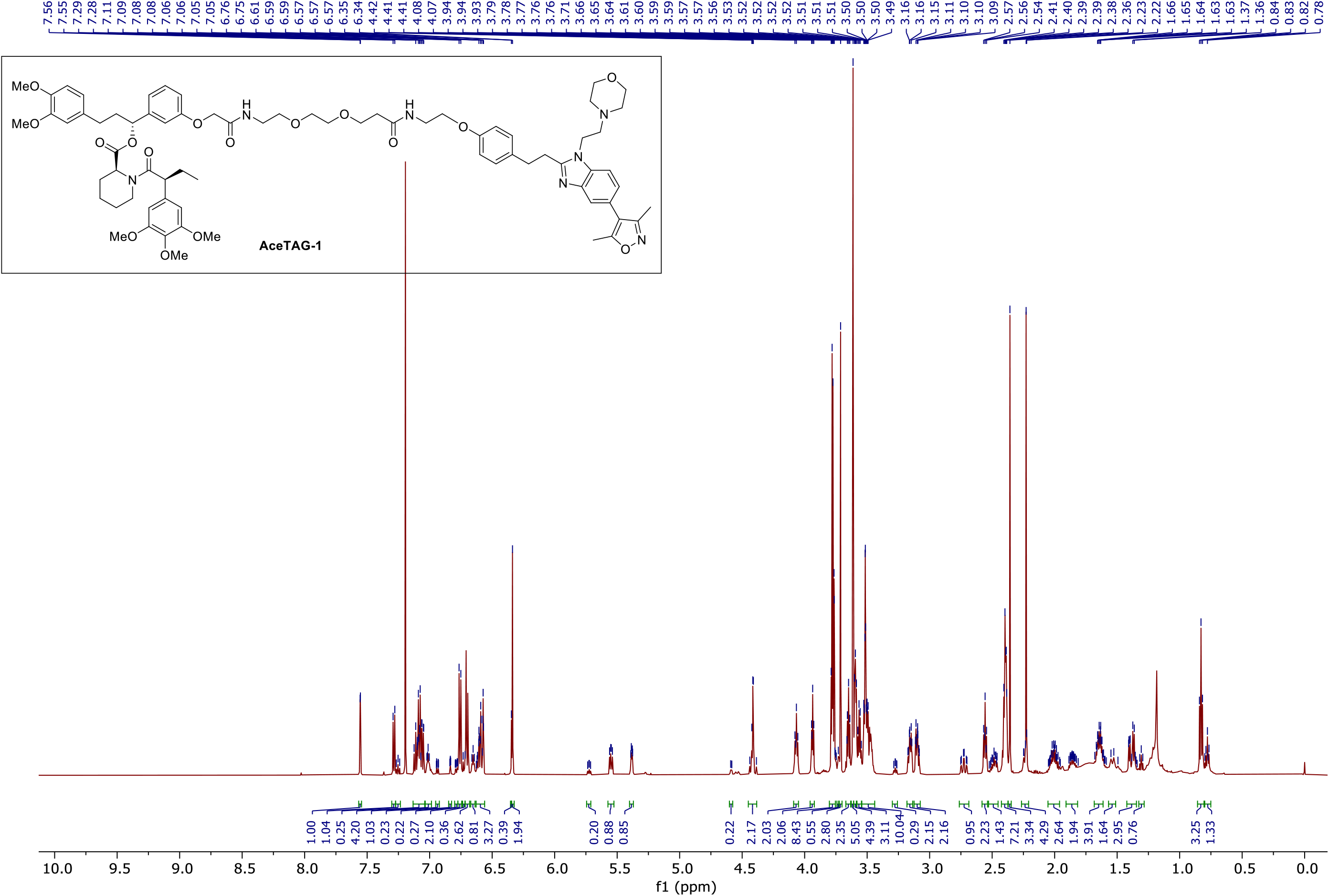

### ^13^C NMR (AceTAG-1)

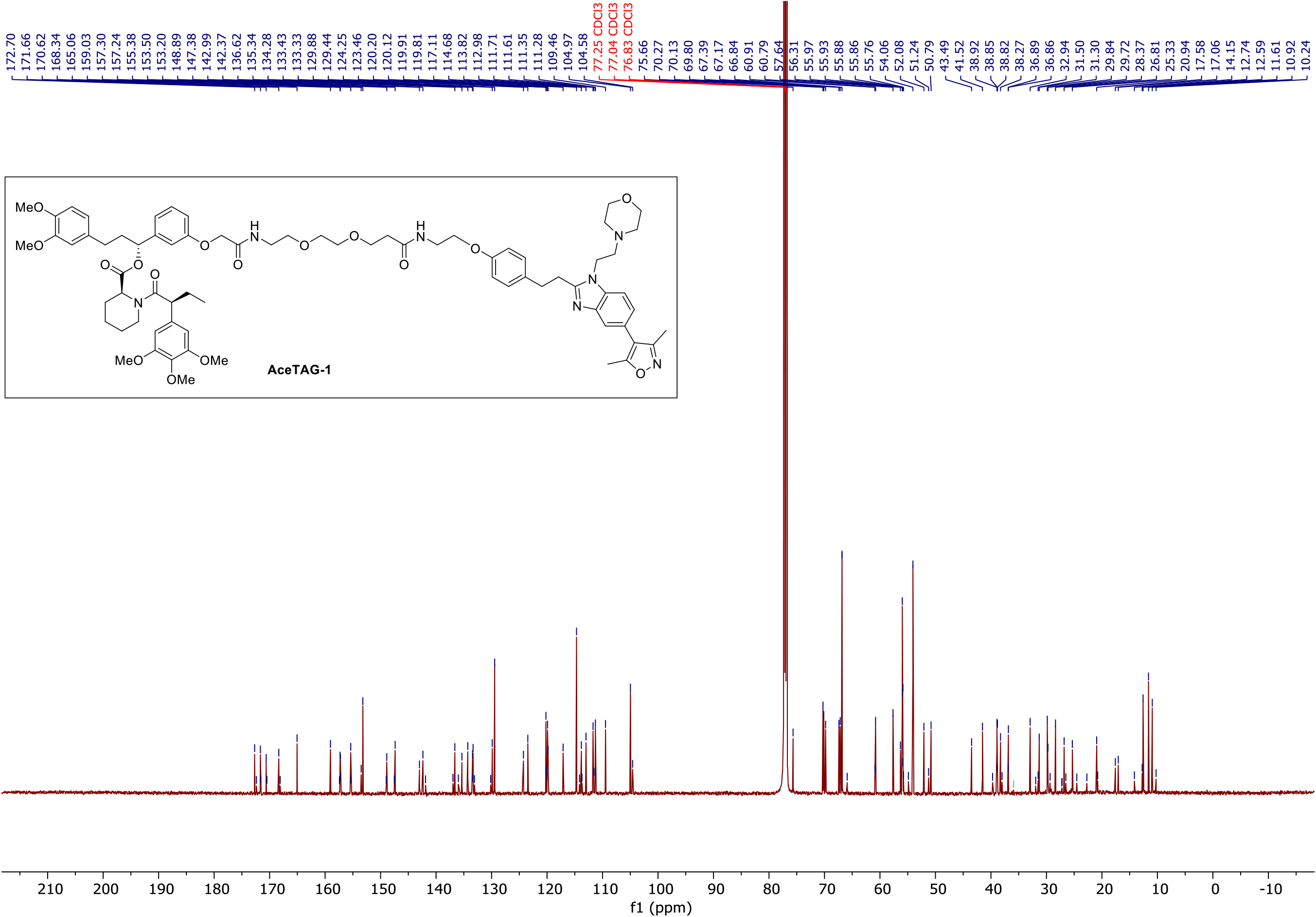

### ^1^H NMR (AceTAG-2)

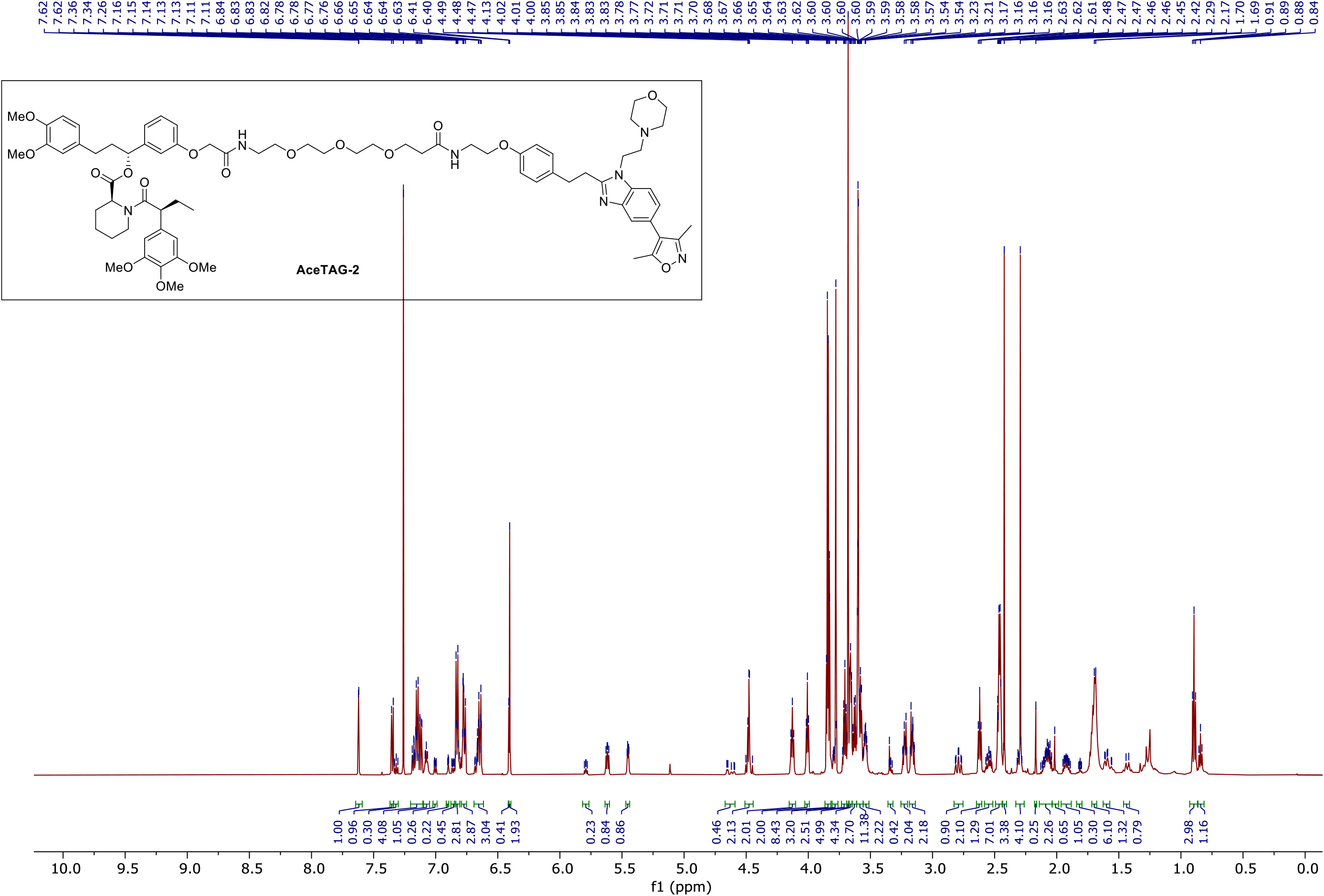

### ^13^C NMR (AceTAG-2)

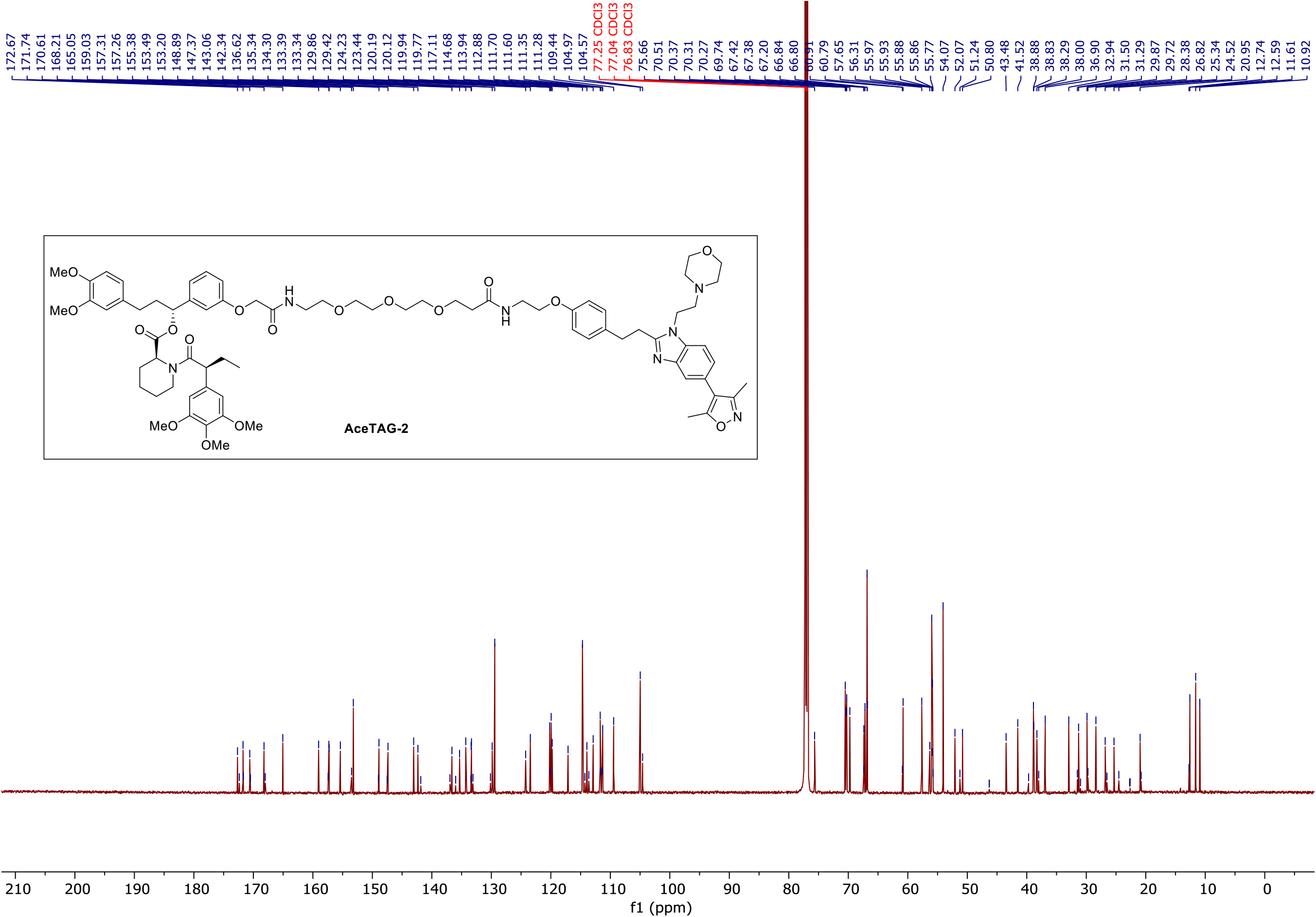

### ^1^H NMR (AceTAG-3)

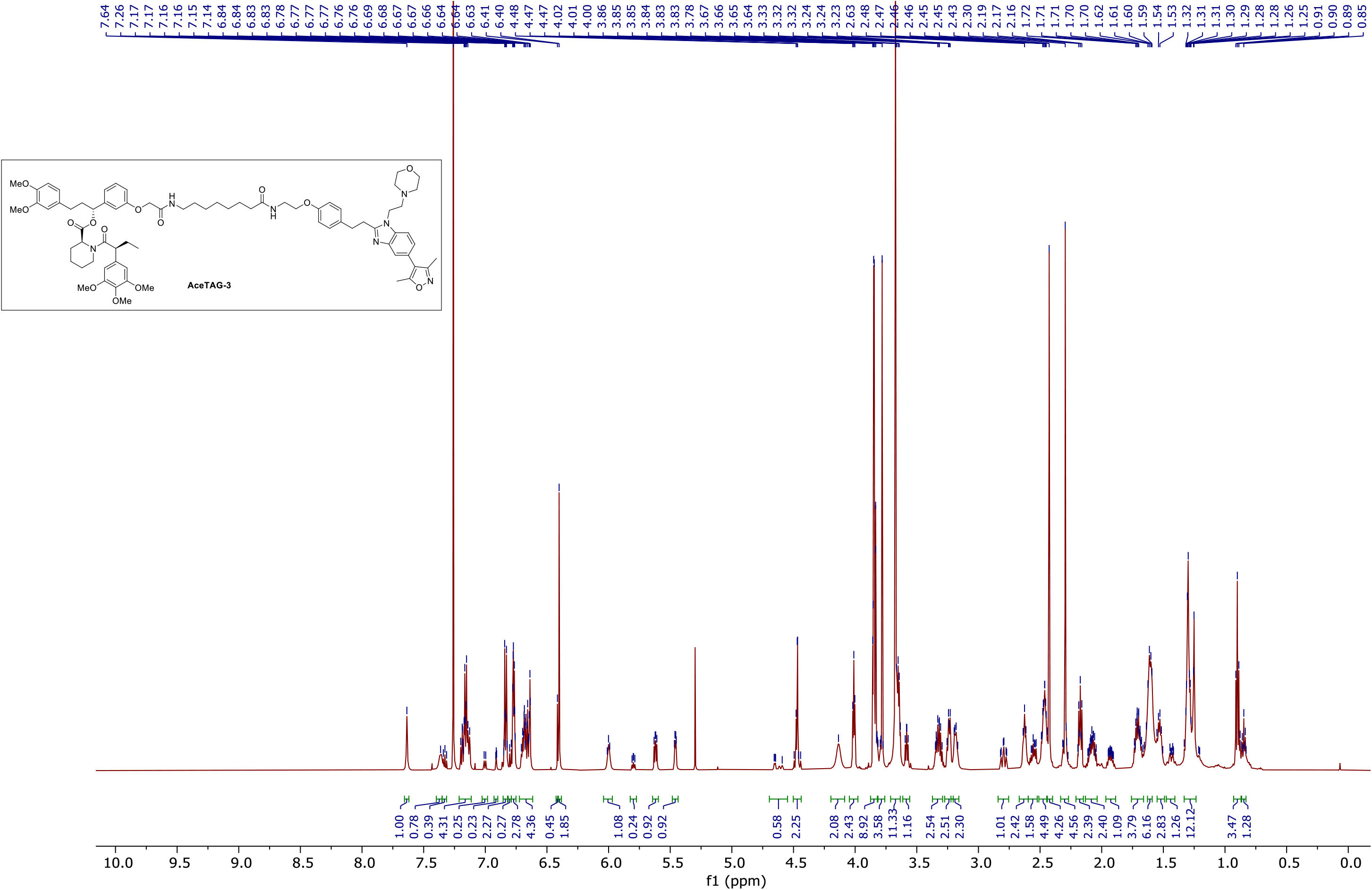

### ^13^C NMR (AceTAG-3)

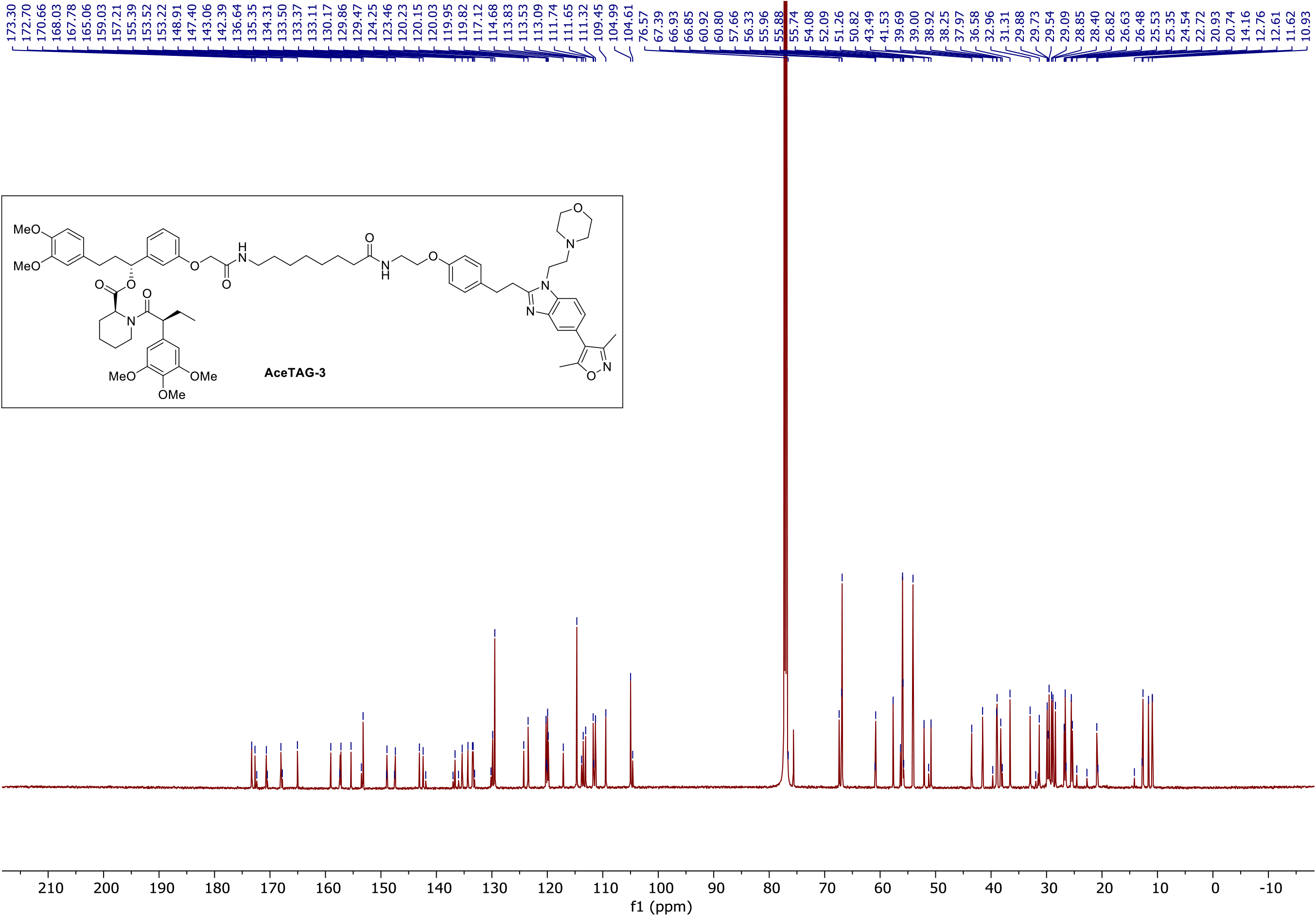

